# MetaGut: Insights into gut microbiomes in stem cell transplantation by comprehensive shotgun long-read sequencing

**DOI:** 10.1101/2023.03.10.531901

**Authors:** Philipp Spohr, Sebastian Scharf, Anna Rommerskirchen, Birgit Henrich, Paul Jäger, Gunnar W. Klau, Rainer Haas, Alexander Dilthey, Klaus Pfeffer

## Abstract

The gut microbiome is a diverse ecosystem, dominated by bacteria; however, fungi, phages/viruses, archaea, and protozoa are also important members of the gut microbiota. Up to recently, exploration of taxonomic compositions beyond bacteria as well as an understanding of the interaction between the bacteriome with the other members was limited due to 16S rDNA sequencing. Here, we developed MetaGut, a method enabling the simultaneous interrogation of the gut microbiome (bacteriome, mycobiome, archaeome, eukaryome, DNA virome) and of antibiotic resistance genes based on optimized long-read shotgun metagenomics protocols and custom bioinformatics. Using MetaGut we investigated the longitudinal composition of the gut microbiome in an exploratory clinical study in patients undergoing allogeneic hematopoietic stem cell transplantation (alloHSCT; n = 31). Pre-transplantation microbiomes exhibited a 3-cluster structure, associated with *Bacteroides*/*Phocaeicola*, mixed composition and *Enterococcus* abundances. MetaGut revealed substantial inter-individual and temporal variabilities of microbial domain compositions, human DNA, and antibiotic resistance genes during the course of alloHSCT. Interestingly, viruses and fungi accounted for substantial proportions of microbiome content in individual samples (up to >50% and >20%, respectively). After leukopenia, strains were stable or newly acquired. Our results demonstrate the disruptive effect of alloHSCT on the gut microbiome and pave the way for future studies based on long-read metagenomics.

## Introduction

Allogeneic hematopoietic stem cell transplantation (alloHSCT) is a potentially curative treatment for patients with high-risk hematological malignancies encompassing acute myeloid leukemia, high-risk myelodysplastic syndromes (MDS), and lymphoid leukemia (Rafiee et al. 2021, Xu and Huang 2021). It is well-established that the gut bacteriome has an influence on the outcome of alloHSCT, including the occurrence of adverse events such as graft-versus-host-disease (GvHD) or life-threatening infections. Recently, adverse outcomes, including the risk of GvHD, have been linked to reduced bacterial diversity of the gut microbiome (Ilett et al. 2020, Peled et al. 2020, Schluter et al. 2020, Taur et al. 2014), while the risk of bloodstream infections has been linked to the domination of individual taxa within the gut bacteriome (Ilett et al. 2020, Montassier et al. 2015, Peled et al. 2020, Taur et al. 2014). A set of microbial taxa, including *Enterobacteriaceae*, *Clostridiales,* and *Blautia* have been implicated in alloHSCT success (Holler et al. 2014, Jenq et al. 2015, Peled et al. 2017, Taur et al. 2014). While a number of explanations for the observed associations have been put forward, including modulation of the immune system by microbiota-derived components and microbiome-host crosstalk at the level of metabolites, the observed associations - apart from an association of overall diversity with outcome, which has been replicated in international multi-center studies (van der Velden et al. 2013) - often remain inconsistent (Ilett et al. 2020, Shono et al. 2016) and incompletely understood.

The vast majority of microbiome studies were performed using ribosomal DNA (rDNA) based sequencing methods designed for bacterial and fungal 16/23S, 18/28S or ITS region rDNA sequences, respectively, thus neglecting the archaeome, non-fungal eukaryome, and the DNA virome (Borrel et al. 2020, Kurilshikov et al. 2021, Richard and Sokol 2019, Sen and Thummer 2022, Stockdale and Hill 2021, Vandeputte et al. 2017, Yalcin et al. 2022, Zhernakova et al. 2016). rDNA sequencing is a well-established, scalable, and cost-effective technology; it has, however, important limitations as it bears potential amplification-induced biases in bacteriome/mycobiome composition estimates and challenges in reliably assigning accurate species - or genus-level labels (Edgar 2018). 16S/18S rDNA sequencing is thus ill-suited to characterize the potential contributions to alloHSCT outcomes of other domains of microbial life. For example, in the context of infection prevention, fungi and protozoan parasites like *Cryptosporidium* spp. and *Toxoplasma gondii* can become relevant in the clinical management of alloHSCT (Danziger-Isakov 2014, Rauwolf et al. 2021); and *Candida* has been associated with GvHD (van der Velden et al. 2013) and survival (Malard et al. 2021, Rolling et al. 2021). With regard to viruses, increases in persistent DNA viruses and reduced bacteriophage richness were observed to be associated with enteric GvHD (Legoff et al. 2017).

A challenge in characterizing associations between microbiome and alloHSCT outcomes consists in the highly dynamic nature of the gut microbiome over the course of alloHSCT (Taur et al. 2012). The gut microbiome undergoes substantial temporal variation even in healthy control individuals (Vandeputte et al. 2021); In the context of alloHSCT, patient microbiomes have often been impacted by multiple cycles of cytostatic and antiinfective therapeutic treatment prior to the initiation of alloHSCT (Dethlefsen and Relman 2011), and continue to be biased by the effects of anti-bacterial, anti-fungal, and anti-viral prophylaxis, in addition to the effects of myeloablation and the subsequent establishment of a “new” immune system by the transplanted allograft (Le Bastard et al. 2021).

Longitudinal studies taking the aforementioned aspects into account are sparse, thus, we developed a novel method (MetaGut) to enable whole-microbiome profiling of patient microbiomes, covering all domains of microbial life (with the exception of RNA viruses). Interrogation of non-bacterial domains of microbial life from shotgun metagenomics is challenging (Hayes et al. 2017, Lind and Pollard 2021, Rose et al. 2016); reasons for this include contamination (Lu and Salzberg 2018, Steinegger and Salzberg 2020) and coverage gaps in the relevant reference databases, likely remaining despite significant recent expansion efforts (Camarillo-Guerrero et al. 2021, Nayfach et al. 2021). We thus chose to implement the shotgun metagenomics step of MetaGut using long-read sequencing, based on the Oxford Nanopore technology, as the accuracy of taxonomic assignment generally increases with read length (Dilthey et al. 2019), and assembled a comprehensive 292 Gb reference database (MetaGut database v 1.0) as the basis of the MetaGut bioinformatics workflow. Furthermore, k-mer-based classification is known to be sensitive to mis-classification in the presence of out-of-database genomes (Dilthey et al. 2019), which, the utilization of a large reference database notwithstanding, remains a relevant concern in the gut microbiome context. We thus propose to complement initial k-mer-based classification with a mapping-based verification approach to reduce the rate of false-positive taxonomic detections. Tailored bioinformatics increased the analytical accuracy and reduced the rate of false-positive taxonomic detections. Moreover, bioinformatic tools for the enumeration of antibiotic resistance genes were implemented. Based on this, we applied MetaGut to characterize microbiome dynamics over the course of alloHSCT in an explorative clinical study including patients (n = 31) before alloHSCT, during the phase of leukopenia, and hematological reconstitution. In addition, a comprehensive characterization of gut microbiomes was performed for a group of healthy volunteers (n = 11) and compared to patient microbiomes pre-Tx. MetaGut proved to be a valuable tool for the comprehensive characterization of microbiomes. Furthermore, a 3-cluster structure of pre-Tx patient microbiomes was detected and the acquisition and replacement of bacterial strains during the course of alloHCST could be successfully monitored.

## Results

### MetaGut, a method for the accurate and comprehensive characterization of gut microbiomes

To enable the characterization of the gut microbiome encompassing the bacteriome, mycobiome, archaeome, DNA virome (bacteriophages/viruses), and protozoa of hematological patients from stool samples at high-resolution, we developed MetaGut, an integrated method for robust microbiome characterization (Figure 1). MetaGut comprises the following components: (i) protocols for stool sample processing and DNA extraction, based on a modified version of the Human Microbiome Project (HMP) protocol (Anon (n.d.), Integrative HMP (iHMP) Research Network Consortium 2019) for robust sample handling and DNA extraction and suitable for processing samples at different degrees of stool consistency (Methods); (ii) long-read sequencing and compositional analysis of microbiota, based on the Oxford Nanopore platform and a custom bioinformatics approach integrating Kraken 2-based read assignments (Wood et al. 2019) with a newly developed mapping-based validation to ensure that sufficient-quality pairwise alignments exist between reads and the taxonomic entities they are assigned to; taxa with low rates of mapping-based read validation are flagged and not included in many downstream analyses and visualizations (see Methods for details); (iii) targeted antibiotic resistance gene (ARG); and (iv) crAssphage analyses, based on read mapping to specific databases comprising (a) ARGs and (b) recently assembled crAssphage strain sequences (Gulyaeva et al. 2022); (v) longitudinal tracking of strain dynamics: short-read Illumina sequencing data are used to detect single-nucleotide variants (SNVs) in bacterial reference genomes, and the temporal dynamics of the detected SNVs over multiple samples from the same individual are investigated to infer the maintenance, acquisition and loss of bacterial strains. Extraction of sufficient amounts of DNA for metagenomic sequencing is challenging, which is particularly relevant during the period immediately following alloHSCT, when patient stool samples are known to vary in consistency over the course of alloHSCT, rendering the exhibit very low biomass. We thus developed and incorporated a modified version of the HMP gut microbiome protocol (Anon (n.d.), Integrative HMP (iHMP) Research Network Consortium 2019) into MetaGut, which proved to work well in our study across the course of alloHSCT. We carried out two experiments as an initial validation of MetaGut.

**Figure 1:**
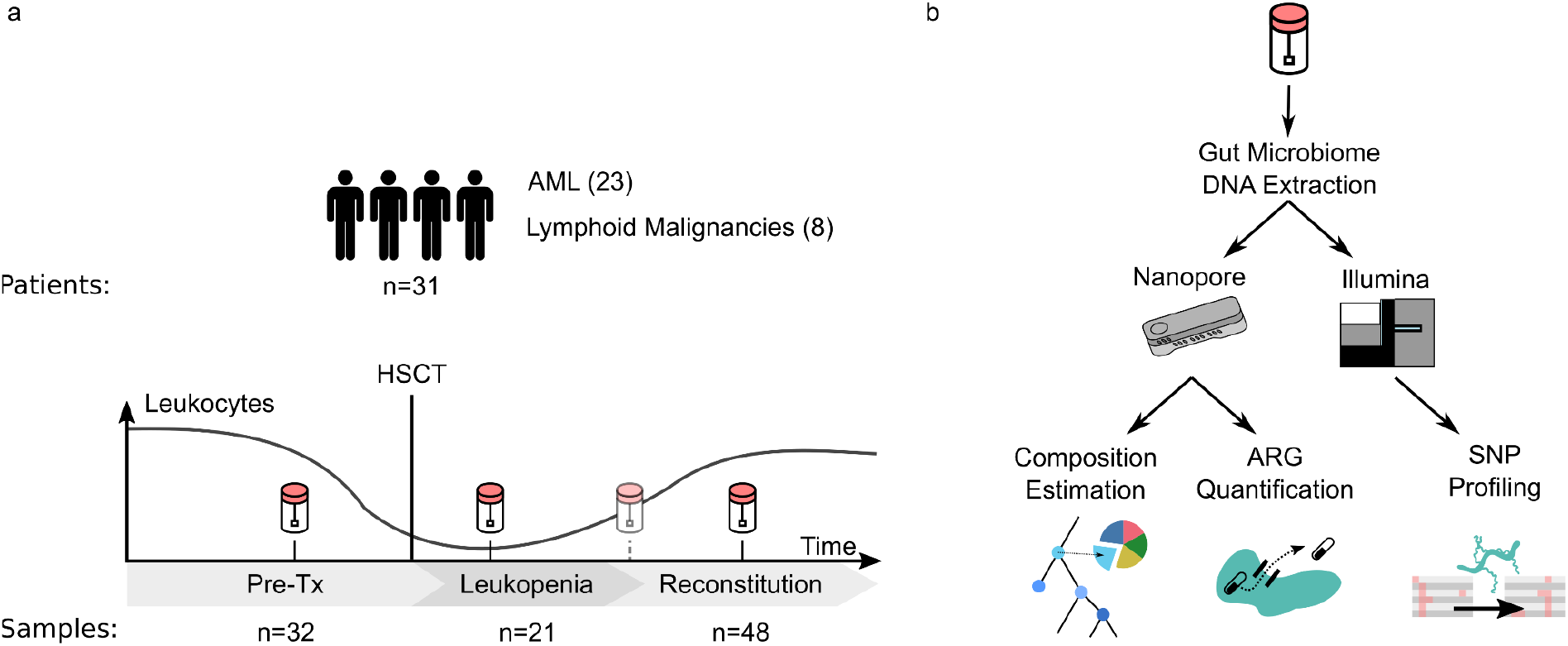
Overview of the exploratory clinical study (a) and the MetaGut method (b). **a)** 31 patients undergoing alloHSCT were recruited for an exploratory study and stool samples were collected longitudinally before transplantation (pre-Tx), during leukopenia (defined as white blood count of ≤ 1,000 /µL), and during reconstitution. **b)** MetaGut comprises optimized protocols for sample preparation and DNA extraction, long-read metagenomics using the Oxford Nanopore technology, and custom bioinformatics for taxon validation, quantification of antibiotic resistance genes / crAssphage sequences, and tracking of strain dynamics based on short-read Illumina data.

First, we applied MetaGut to a well-defined microbial community standard (Zymo Gut Microbiome Standard) and observed good concordance at the genus level between the theoretical and the MetaGut-inferred composition (Pearson’s r = 0.80, retaining the category “Unassigned At Level”; Supplementary Figure 1). Of note, an initial analysis of the Zymo data at the species level identified an elevated proportion of false-positive species hits, accounting for 32.43% of total abundance and driven by reads from *Veillonella rogosa* and *Prevotella corporis* being misassigned to different species of their respective genera (Supplementary Table 1). Therefore, all subsequent analyses were carried out at the genus level, which exhibited a markedly lower rate of false-positive hits (accounting for 5.55% of total read abundance).

Second, we applied MetaGut to a cohort of 10 healthy volunteers. Here, we observed good agreement of high-level compositional metrics with the WGS-based component of LifeLines-DEEP (Tigchelaar et al. 2015) (Supplementary Figure 2). Of note, at a median frequency of 34.28% (range 11.72% - 66.98%), the MetaGut-inferred abundance of *Bacteroides* was higher in the investigated cohort of healthy volunteers compared to the LifeLines-DEEP cohort with a median frequency of 20.95% (range 1.57% - 56.70%).

### MetaGut enabled reliable characterization of gut microbiomes over the course of alloHSCT

To characterize the microbiomes of alloHSCT patients and to investigate the dynamic changes of the gut microbiome over the course of alloHSCT, we recruited a cohort of 31 patients, diagnosed with defined hematological malignancies (see Supplementary Note 1 and Methods for a description of the cohort and recruitment process, and Supplementary Table 2 for patient characteristics), undergoing alloHSCT at Düsseldorf University Hospital.

Based on 101 stool samples collected longitudinally at defined time points over the course of alloHSCT (pre-Tx, leukopenia, reconstitution / Figure 1), we assessed the ability of MetaGut to enable microbiome characterization at different stages of the treatment cycle (Figure 2). First, we observed that DNA yields in the hematological cohort differed significantly from the healthy cohort (patients 0.47 µg/g stool, healthy 5.95 µg/g stool; p = 0.000011) and also displayed significant variation between treatment time points (Kruskall-Wallis p-value: 0.0031 for the 3 treatment phases). DNA yields were typically lowest during leukopenia (median = 0.06 µg/g stool), compared to the pre-Tx and reconstitution periods (medians = 0.6 µg/g stool and 1.1 µg/g stool, respectively). Second, the utilized protocols enabled the generation of more than 100,000 reads per sample (89 samples with >= 100,000 reads). We observed that median read counts were lowest for samples taken during leukopenia (median = 128,330). Third, the median of the median sample read lengths was 653 base pairs (bp) with the longest read spanning 888,591 bp. DNA extraction and sequencing data statistics are summarized in Supplementary Table 3. Fourth, at median per-genus validation rates of 79.29% (bacteria) and 57.86% (viruses) across treatment periods and in the healthy cohort showed generally high rates of mapping-based validation (Supplementary Figure 3). Fungi exhibited lower validation rates (median = 6.67%); we observed pronounced differences in fungal validation rates between the healthy and alloHSCT samples, and also between the characterized time points, with the healthy samples generally exhibiting the lowest fungal validation rates (Supplementary Figure 3).

**Figure 2:**
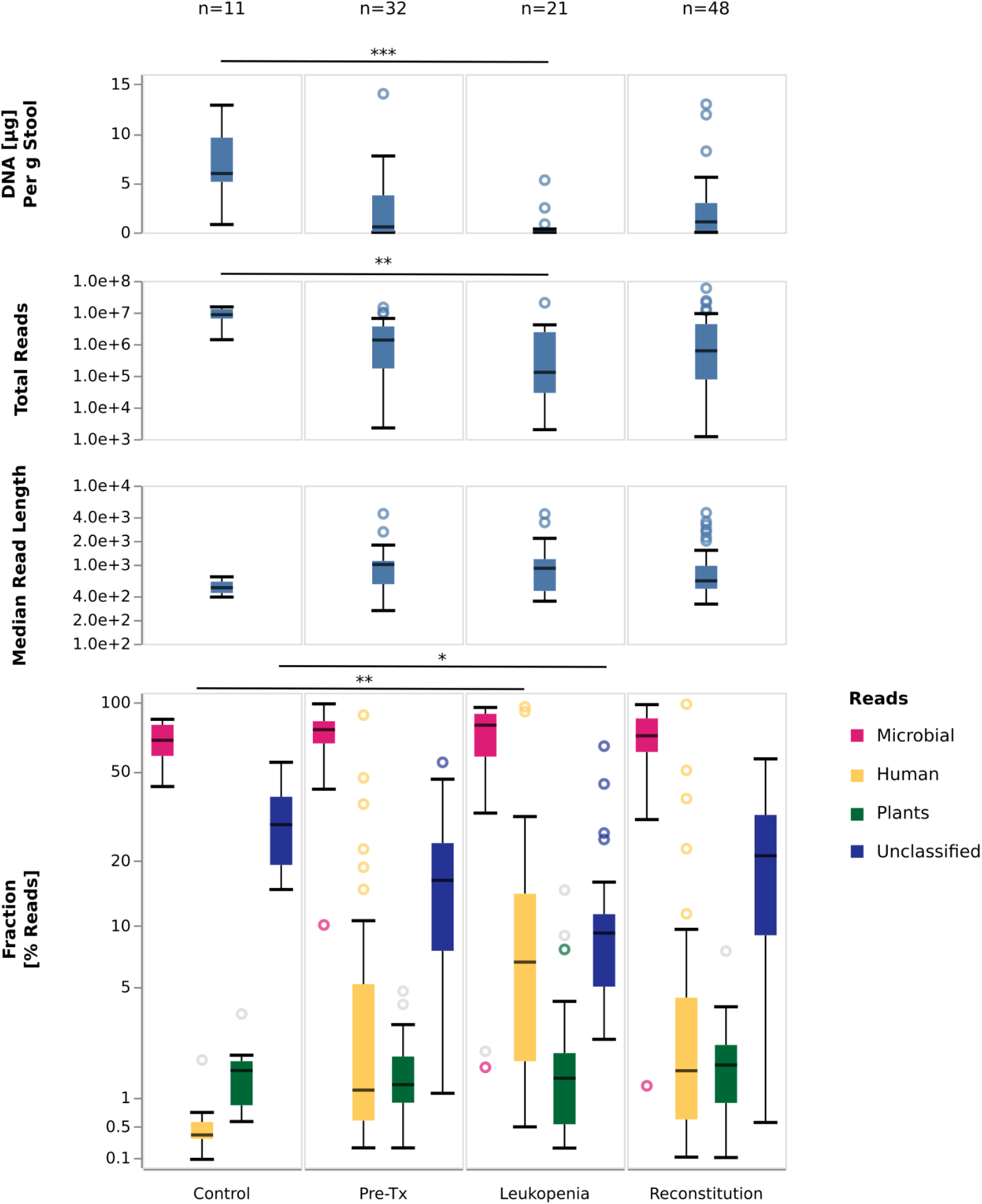
DNA amounts, sequencing metadata, and high-level stool sample compositions. Shown are the amounts of extracted DNA per gram of stool; the number of generated Nanopore sequencing reads; median read lengths; and the relative proportions of microbial-, human-, and plant-assigned reads, as well as the proportions of reads remaining unclassified by the initial Kraken 2-based step of MetaGut. Gray circles in the bottom panel indicate outliers for which the underlying taxa could not be validated using the mapping-based validation step of MetaGut. For each panel, the most significant difference between included categories is indicated (Mann-Whitney U test; * [p<1.19×10^-3^], ** [p<2.38×10^-4^] or *** [p<2.38×10^-5^]). Raw data are shown in Supplementary Table 3 and Supplementary Table 4.

Human-derived DNA is often ignored in microbiome studies, but we noted that the amount of human DNA differed significantly between the control group and the patient cohort, and also between the sampling time points within the hematological cohort. The relative proportion of human DNA was highest during the leukopenic period, possibly reflecting the effect of myeloablative therapy on the gut mucosa (Figure 2). We also observed that the per-read validation rates of the human reads were lowest for the control samples, i.e. indicating that a substantial proportion of the reads assigned by Kraken 2 to the genus “Homo” in these samples may reflect false-positive assignments (Supplementary Figure 3).

Per-genus validation rates for archaea and non-fungal eukaryotes were close to zero in most control and alloHSCT samples and also across the characterized time points. Analyses of the occurrence and abundance of individual genera were thus limited to the bacterial, viral and fungal components of the microbiome. Of note, the presence of plant-based DNA could also sometimes be validated, in particular during the leukopenic period (Supplementary Figure 3).

### alloHSCT microbiomes exhibited dynamic and diverse compositions

Having established and validated MetaGut as an accurate and innovative method, we investigated high-level gut microbiome compositions (Figure 3). The number of normalized genera was highest in the samples from the control group and lowest for samples during leukopenia (Figure 3, top panel). Besides, ARG-reads significantly differed between control samples (low) and alloHSCT samples (high) (Figure 3, middle panel). The frequency of ARG reads of the control samples was significantly lower compared to alloHSCT samples (p = 0.0006281). Bacteria accounted for > 90% of microbial reads in 89/101 hematological samples and in 11/11 of control samples; bacteria thus dominated the large majority of the investigated samples. At median relative abundances of 0.53%, 0.11% and 0.02%, fungi, viruses, and archaea accounted for low, but non-negligible proportions of the characterized alloHSCT and control microbiomes; median abundances of these groups were generally comparable between control and alloHSCT samples and between alloHSCT time points, with the exception of fungi, which showed an increased abundance in the alloHSCT samples in general and during leukopenia in particular (median = 1.32% vs 0.36% for control) (Figure 3, bottom panel). Non-fungal eukaryotes were also estimated to account for small fractions of the characterized microbiomes (median = 0.49%).

**Figure 3:**
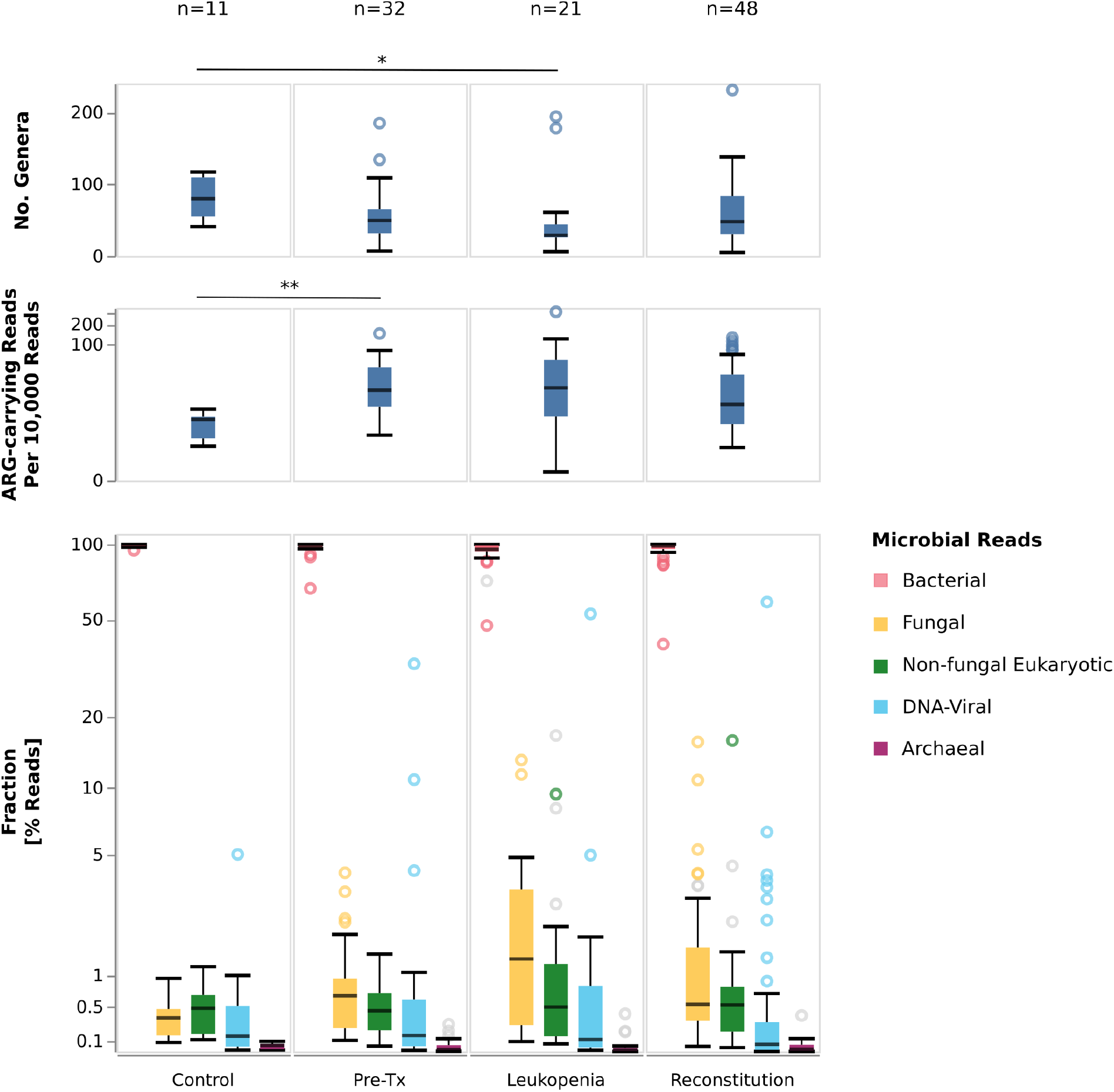
Diversity, ARG-carrying reads, and microbiome composition in stool samples from controls and alloHSCT patients at indicated phases. Shown are the numbers of detected genera, normalized to the sample with the lowest read count; the rate of reads that carry an ARG element; and the relative proportions of bacterial-, fungal-, archaeal-assigned reads as well as the proportion of reads assigned to DNA viruses (incl. bacteriophages), and non-fungal eukaryotes. Gray circles in the bottom panel indicate outliers for which the underlying taxa could not be validated using the mapping-based validation step of MetaGut. For each panel, the most significant difference between included categories is indicated (Mann-Whitney U test; * [p<1.19×10^-3^], ** [p<2.38×10^-4^] or *** [p<2.38×10^-5^]). Raw data are shown in Supplementary Table 3 and Supplementary Table 4

The dominance of bacteria in most samples notwithstanding, individual microbiome samples were found to exhibit increased frequencies of non-bacterial taxa. For example, in two alloHSCT samples from one patient, the abundance of viral DNA was found to be ≥ 50%, accounted for by the genus crAssphage; in two additional samples from different patients, it was ≥ 10%. Interestingly, in one control sample, crAssphage abundance approached 5% (Figure 3, bottom panel). Similarly, in five samples, fungal DNA was found at abundances ≥ 5%, accounted for mostly by the genera *Saccharomyces*, *Malassezia*, and *Candida* (Supplementary Table 6). In all of these instances, taxon presence was validated with read-mapping based verification.

### Pre-Tx alloHSCT microbiomes could be grouped into 3 distinct clusters

To investigate the structure of the pre-transplantation gut microbiome in alloHSCT patients, we carried out a PCoA analysis and found that pre-Tx alloHSCT and control samples fell into 3 distinct clusters (Figure 4). Cluster 1, comprising 18 pre-Tx alloHSCT samples from distinct patients and all 11 control samples, was characterized by high abundances of *Bacteroides* and *Phocaeicola,* accompanied by relatively high normalized numbers of detected genera per sample; specifically, *Bacteroides* and *Phocaeicola* accounted for ≥ 50% of reads in 10/18 of Cluster 1 samples, while the median number of normalized detected genera per sample was 55, showing a high diversity. Cluster 3, comprising 6 pre-Tx alloHSCT samples, was characterized by *Enterococcus* domination and exhibited the lowest diversity in terms of the number of detected genera per sample; in 5/6 Cluster 3 samples, *Enterococcus* accounted for > 50% of sequencing reads, at a median number of 26 detected normalized genera per sample. Cluster 2, comprising 8 pre-Tx alloHSCT samples, was the most diverse in terms of the number of detected normalized genera per sample (median = 62) and was also characterized by more uniform genus abundance distributions. In Cluster 2, a median of 3 genera were required to account for ≥ 50% of sequencing reads, compared to medians of 2 and 1 genera in Clusters 1 and 3, respectively. Notably, Cluster 2 genera contained in individual samples included *Citrobacter* (max. abundance 51%)*, Lactobacillus* (max. abundance 44%), and *Escherichia* (max. abundance 31%); by contrast, the genera characteristic for Cluster 1 and 3 *(Bacteroides / Phocaeicola*, *Enterococcus*) were present at only low abundances in Cluster 2 (abundances ≤ 10 % for 6/8 samples of Cluster 2).

**Figure 4:**
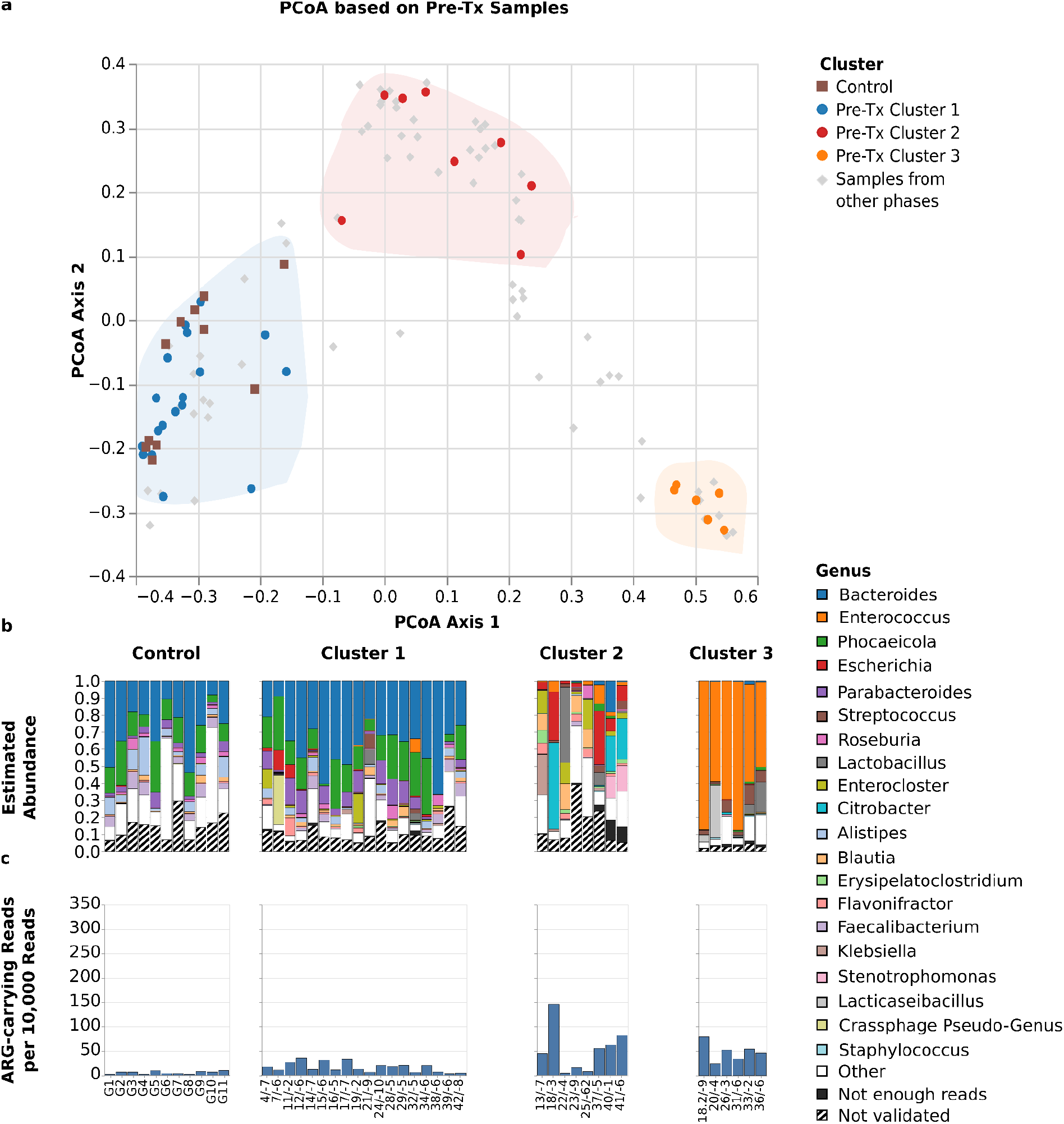
Microbiome structures and compositions of pre-Tx alloHSCT samples and healthy controls. The Figure shows **a)** the positions of pre-Tx and control samples, displayed as colored dots, in the joint PCoA space of all samples, as well as the positions of pre-Tx Cluster 1, Cluster 2, and Cluster 3 in PCoA space (shaded areas), **b)** bar plots visualizing the microbiome compositions of healthy control and pre-Tx alloHSCT samples, stratified by pre-Tx cluster membership, and **c)** the rate of reads that carry an ARG element, separately for control and pre-Tx alloHSCT samples and stratified by pre-Tx cluster membership. The bar plots show the 20 genera that attained the highest aggregated sum of frequency across all time points, and, of these, within each sample, only the genera that 1.) were assigned at least 20 reads to and 2.) passed the mapping-based taxon validation step of MetaGut were depicted. The combined abundances of genera with fewer than 20 reads is shown in the category “not enough reads”; the category “not validated” shows the combined abundances of genera with more than 20 Kraken2-assigned reads that did not pass the mapping-based taxon validation step of MetaGut. Sample labels below the bar plots specify patient or control sample ID, followed, for alloHSCT samples, by the day of sampling relative to the stem cell transplantation time point.

Suggestive structural differences between the 3 pre-Tx clusters were also apparent at the level of the fungal and viral microbiome components (Figure 7). *Saccharomyces* (validated presence in 21 samples and >10% relative within-fungal abundance in 16 samples) and *Candida* (validated presence in 3 samples and >10% within-fungal relative abundance in 3 samples) were the most abundant fungal genera detected during the pre-Tx period; the presence of *Candida*, however, was limited to Cluster 2 and 3 samples. At the level of the DNA virome, *Skunavirus* dominated (>50% relative abundance) 8 of 14 Cluster 2 and Cluster 3 samples, but only 2 of 18 Cluster 1 samples. Conversely, crAssphage was found mostly in Cluster 1 samples (6 validated detections in total, 5 of which occurred in Cluster 1 samples); and 2 Cluster 1 samples were crAssphage-dominated, but no samples from Clusters 2 or 3.

### Microbiomes in leukopenia exhibited decreased diversity and increased abundances of fungi and various bacterial genera

We proceeded to investigate the structure of microbiome samples collected during the leukopenic period, reflecting the combined effects of myeloablation, anti-infective prophylaxis, and recent alloHSCT. Consistent with an assumed bottleneck effect of anti-infective prophylaxis on the population of gut microbes, the leukopenic period exhibited the lowest median number of normalized detected microbial genera per sample (29; compared to 46 and 48 for the pre-Tx and reconstitution periods, respectively; Figure 3). The proportion of the microbiome accounted for by fungal taxa (mycobiome), however, was substantially increased during leukopenia (median per sample: 1.32% of reads, compared to 0.65 and 0.54 for pre-Tx and reconstitution, respectively; Figure 2); the main detected fungal genera were *Saccharomyces* (1.52% mean absolute abundance across leukopenia samples) and *Candida* (0.04% absolute mean abundance).

Changes in microbiome composition compared to the pre-Tx period were also apparent at the level of individual bacterial genera; *Enterococcus*, *Streptococcus*, *Lactobacillus*, and *Erysipelatoclostridium* were detected at abundances ≥ 5% in 57%, 38%, 28%, and 14% of samples during leukopenia, respectively, compared to 28%, 16%, 16%, and 3% for pre-Tx samples (Supplementary Table 5 and Figures 4 and 5). Of note, the observed overall expansion of *Enterococcus* during leukopenia was driven by samples from patients assigned to pre-Tx Cluster 1; patients assigned to the *Enterococcus*-associated pre-Tx Cluster 3, by contrast, generally showed a reduction in median *Enterococcus* abundance (66% pre-Tx, 15% leukopenia). Interestingly, changes at the level of individual genera similar to bacteria were not detected for fungi and DNA viruses; that is, almost all fungal and viral genera detected at >5% validated relative abundance in leukopenic samples were also detected at >5% validated relative abundance in pre-Tx samples (Supplementary Table 6, Supplementary Table 7).

**Figure 5:**
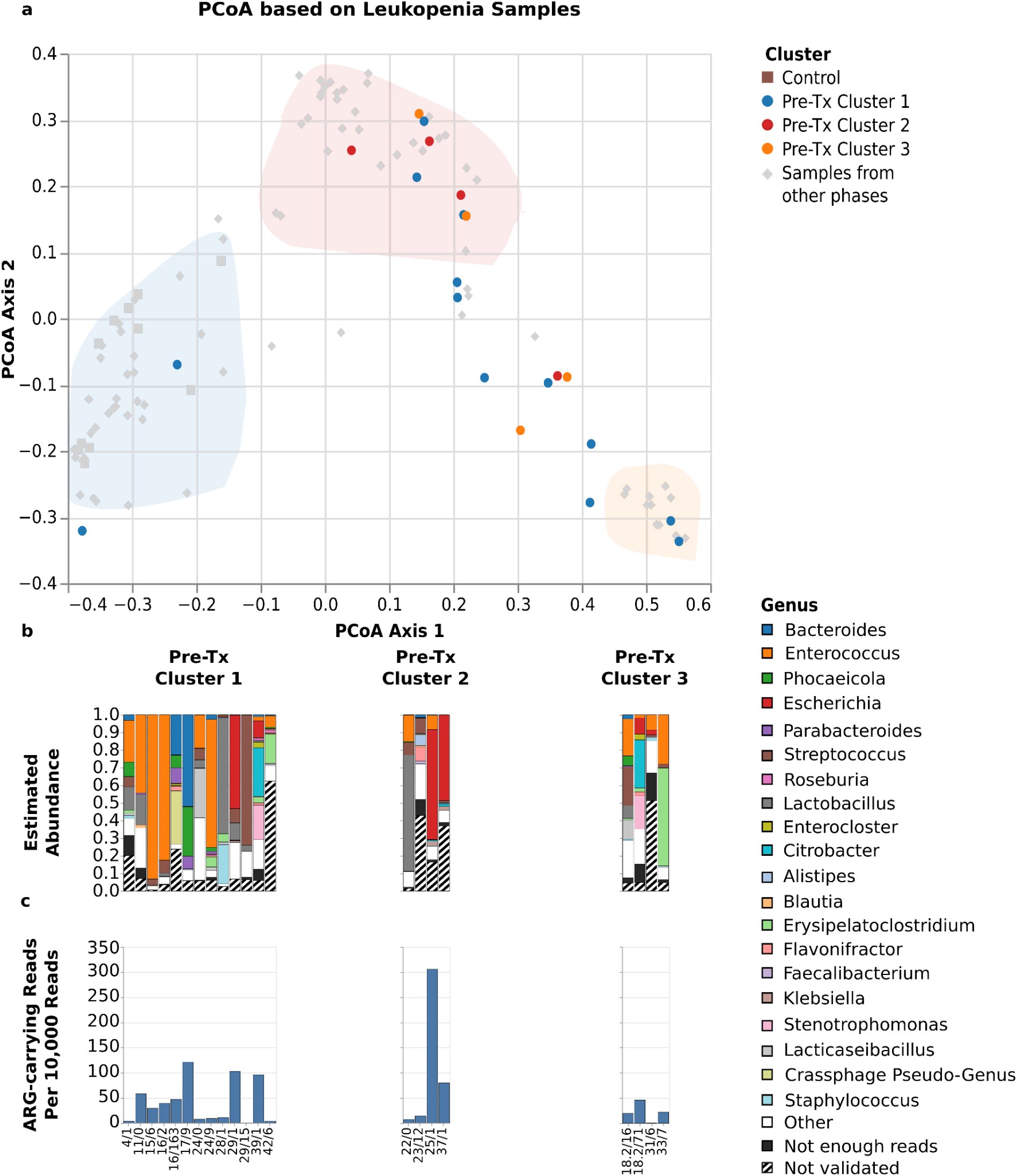
Microbiome structures and compositions during leukopenia. The Figure shows, for alloHSCT samples collected during the leukopenic period, **a)** sample positions in the joint PCoA space of all samples, **b)** bar plots visualizing sample microbiome compositions, and **c)** rates of reads carrying an ARG element. All visualizations are stratified by pre-Tx cluster membership of the corresponding patients; in the top panel, pre-Tx cluster membership is indicated by dot color. The bar plots show the 20 genera that attained the highest mean frequency across all time points, and, of these, within each sample, only the genera that 1.) were assigned at least 20 reads and 2.) passed the mapping-based taxon validation step of MetaGut were depicted. The combined abundance of genera with fewer than 20 reads is shown in the category “not enough reads”; the category “not validated” shows the combined abundance of genera with more than 20 Kraken2-assigned reads that did not pass the mapping-based validation step. Sample labels below the bar plots specify patient ID, followed by the day of sampling relative to the stem cell transplantation time point.

### The reconstitution microbiomes were characterized by increased similarity to the pre-Tx microbiomes

Compared to samples collected during leukopenia, samples during the reconstitution period collectively showed similarity to pre-Tx microbiome samples. During the leukopenic period, approximately 52% (11/21) of collected samples were found to be within one of the 3 clusters defined based on pre-Tx samples; during reconstitution, this fraction increased to 83% (40/48; Figure 6). In addition, reconstitution microbiomes also resembled pre-Tx microbiomes at the level of observed overall abundances of bacteria, fungi, and DNA viruses (Figure 3); at the level of bacterial genera detected at ≥ 5% abundance (Supplementary Table 5); and at the level of the fungal and DNA viral genera that exhibited the highest rates of validated detection at > 5% relative abundance, i.e. *Saccharomyces* and *Candida* for fungi, as well as as *Skunavirus* and *crAssphages* for DNA viruses (Supplementary Table 6, Supplementary Table 7, Figure 7).

**Figure 6:**
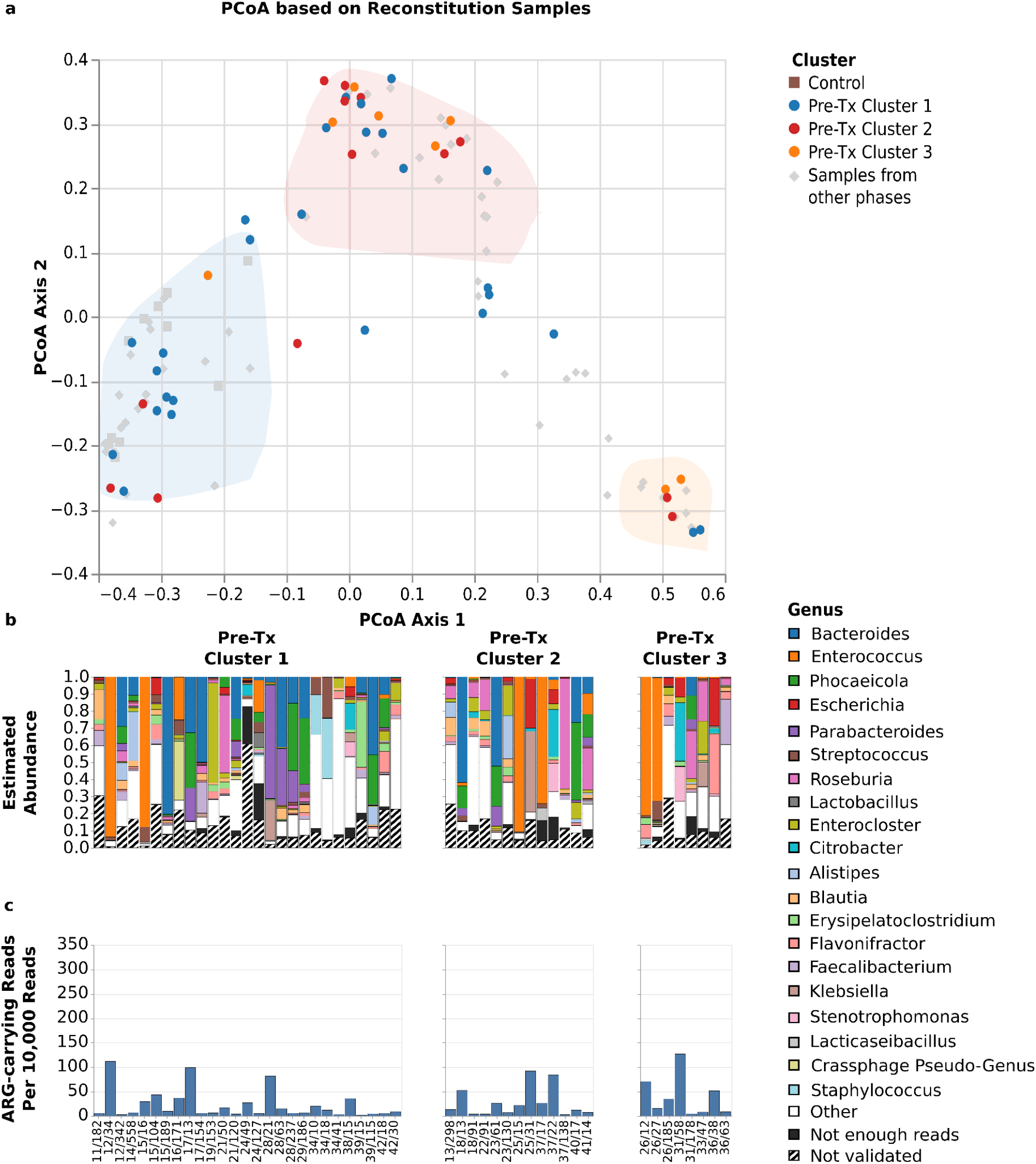
Microbiome structures and composition during reconstitution. The figure shows, for alloHSCT samples collected during the reconstitution period, **a)** sample positions in the joint PCoA space of all samples, **b)** bar plots visualizing sample microbiome compositions, and **c)** rates of reads carrying an ARG element. All visualizations are stratified by pre-Tx cluster membership of the corresponding patients; in the top panel, pre-Tx cluster membership is indicated by dot color. The bar plots show the 20 genera that attained the highest mean frequency across all time points, and, of these, within each sample, only the genera that 1.) were assigned at least 20 reads and 2.) passed the mapping-based taxon validation step of MetaGut were depicted. The combined abundance of genera with fewer than 20 reads is shown in the category “not enough reads”; the category “not validated” shows the combined abundance of genera with more than 20 Kraken2-assigned reads that did not pass the mapping-based validation step. Sample labels below the bar plots specify patient ID, followed by the day of sampling relative to the stem cell transplantation time point.

**Figure 7:**
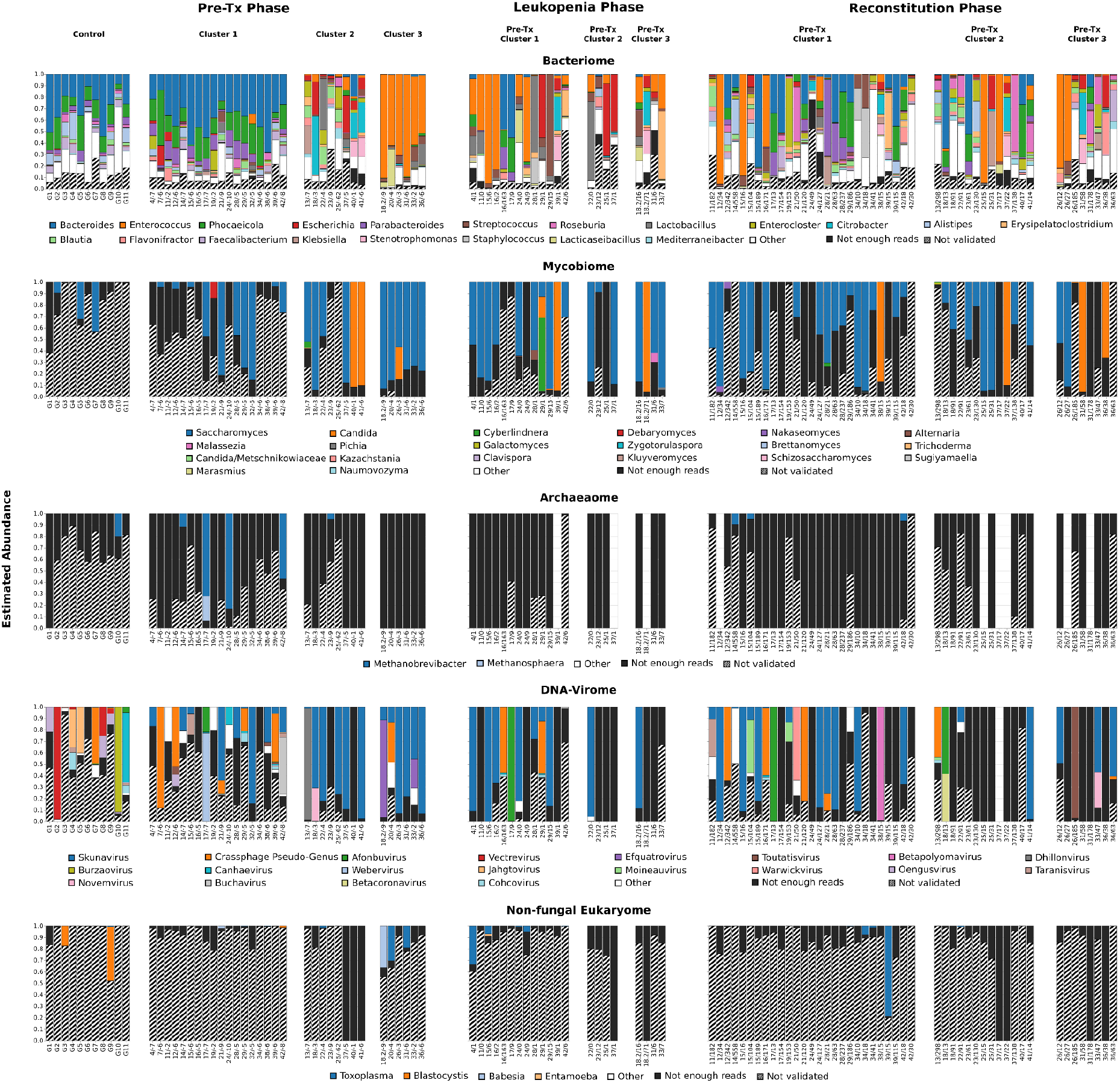
Integrated microbiome analysis of all characterized samples. Shown are, for each sample, bar plots that visualize the respective compositions of the bacterial, fungal, archaeal, DNA-viral, and non-fungal eukaryotic components of the sample microbiome. Normalization was applied independently to each bar plot. The bar plots only show the 20 (for bacteria, fungi and DNA viruses), 4 (for archaea), and 2 (for non-fungal eukaryotes) genera that attained the highest mean relative frequency within the considered taxonomic category across all timepoints, and, of these, within each sample, only the genera that **a)** were assigned at least 20 reads by Kraken2, and **b)** passed the mapping-based taxon validation step of MetaGut. The combined abundance of genera within each taxonomic category with fewer than 20 reads is shown in the category “not enough reads”; the category “not validated” shows the combined abundance of genera with more than 20 reads that did not pass the mapping-based validation step. Sample labels below the bar plots specify healthy control ID or the patient ID, followed by the day of sampling relative to the transplantation event.

Bacterial genera exhibiting notable differences compared to the pre-Tx period included *Roseburia*, which was detected in approximately twice as many samples at abundances ≥ 5% during reconstitution than during pre-Tx (19% compared to 9%), and which accounted for ≥ 50% of overall microbial abundance in individual reconstitution samples; *Parabacteroides*, which exhibited dominance (≥ 50% abundance) in one reconstitution sample, but overall decreased detection at ≥ 5% abundance (17% during reconstitution samples compared to 31% for pre-Tx samples); and *Phocaeicola* and *Bacteroides*, which also exhibited decreased detection rates at the ≥ 5% abundance threshold (29% and 35% for reconstitution samples compared to 47% and 59% for pre-Tx samples, respectively; Supplementary Table 5). While similar differences were not detected for fungal genera Supplementary Table 6), multiple viral genera exhibited exclusive detection at > 5% validated relative abundance in reconstitution samples compared to pre-Tx samples (Supplementary Table 7); while these detections were typically limited to individual samples, in some instances the underlying viral genera dominated the viral microbiome component (observed e.g., in the case of *Betapolyomavirus*; Figure 7).

### Pre-Tx clusters were not associated with longitudinal microbiome trajectories or clinical outcomes

Pre-Tx cluster assignment did not predict the cluster membership of samples from the same patient over the course of alloHSCT, neither during the leukopenic nor during the reconstitution period (Figure 5, Figure 6); that is, pre-Tx cluster membership was not associated with the microbiome trajectory at the level of individual patients. Similarly, the pre-Tx detection of *Candida* or crAssphage was not associated with the detection of these genera during leukocytopenia or reconstitution (Figure 7).

Furthermore, while the proportion of patients with adverse outcomes (relapse and death) or GvHD varied between the 3 clusters (44%, 25% and 67% for adverse outcomes in Cluster 1, 2 and 3, respectively; 11%, 12%, and 17% for GvHD), these differences were not statistically significant (p = 0.297 for serious outcomes; p = 0.938 for GvHD; Pearsons’ chi-squared test). Further, we did not observe an association between cluster membership and the type of hematological disease patients were treated for (Supplementary Table 8).

### MetaGut enabled analyses of antibiotic resistance, viral strain diversity and the tracking of individual marker genera

We further investigated whether the whole-genome microbiome sequencing component of MetaGut could enable inferences about the presence of antibiotic resistance genes (ARGs), viral strain diversity, and the abundances of individual bacterial, fungal, or DNA-viral marker species which were described in previous publications (Supplementary Table 9. Literature List Marker Genera) or which were of particular interest. First, we investigated the detection of ARGs by quantifying the proportion of sequencing reads that aligned against known antibiotic resistance genes; the observed rates of ARG-carrying reads (Figure 4) were relatively similar for pre-Tx and leukopenia alloHSCT samples (median = 20.39 ARG-carrying reads per 10.000 reads for pre-transplantation samples; 22.25 for leukopenia samples), but substantially lower in the reconstitution (median = 12.22) and control (median = 6.95) samples. Of note, we observed lower rates of ARG-carrying reads in pre-Tx Cluster 1 samples than in Cluster 2 and Cluster 3 samples (Figure 4; p = 0.00196); consistent with the observation that pre-Tx Clusters 2 and 3 contained microbiomes which were more divergent from the characterized control samples, possibly due to prior exposure to anti-infective medications. Next, prompted by the observation that crAssphages accounted for substantial proportions of total microbiome content in individual samples and by the availability of a large number of resolved crAssphage strain genomes from a recent publication (Gulyaeva et al. 2022), we investigated the detectability and relative abundances of different crAssphage strains using a mapping-based approach. This analysis showed that (i) mapping against a database containing only crAssphage genomes resulted in substantially higher estimated crAssphage abundances in many samples than classifying against the comprehensive MetaGut database (Supplementary Figure 4), (ii) read mapping enabled sensitive detection and differentiation between different crAssphage strains (Supplementary Figure 5), and (iii) many reads classified as crAssphage in the mapping-based analysis were assigned to other taxa by the Kraken 2-based MetaGut read assignment process (Supplementary Figure 6). Of note, while these results confirmed the applicability of the generated long-read data to crAssphage-focused analyses, they also suggested that substantial crAssphage strain diversity is currently not represented in the comprehensive MetaGut database. Third, we investigated the applicability of MetaGut to tracking longitudinal changes of the abundances of predefined marker genera. To this end, we assembled a list of bacterial genera reported to be associated with alloHSCT outcomes from the literature (Methods); and, to extend this analysis to non-bacterial genera, we added selected fungal and viral genera accounting for substantial proportions of microbiome content (Methods, Supplementary Table 9). We then carried out a longitudinal analysis of the presence and abundance of the selected marker genera over time (Supplementary Figure 7), and found that the MetaGut-generated abundance estimates were informative for changes over time, based on largely non-overlapping abundance confidence intervals between time points, demonstrating, for example, fluctuations in the abundances of *Akkermansia* and *Blautia*.

### Investigation of bacterial strain dynamics showed replacement of strains over the course of alloHSCT in some individuals

Finally, we investigated whether the significant impact of myeloablation, antiinfective therapy, and alloHSCT on the gut microbiome was associated with bacterial strain replacement, potentially indicating re-colonization of ecological niches in the microbiome. We developed a sub-module of MetaGut (Methods) to approximate species-level core genome average identities across samples based on short-read sequencing data, generated Illumina data for a subset of our samples with sufficient DNA yield (Methods), and applied the developed method for the detection of strain replacement events. Based on the generated data, we could characterize longitudinal strain dynamics in 31 cases, representing 6 bacterial species and 14 patients (Figure 8). Of 44 considered intervals, 12 were classified as representing a strain-switching event; of these, 8 spanned an alloHSCT and 2 a relapse event; compared to 19 of the 29 alloHSCT or relapse intervals not representing a strain-switching event. While the rate of strain-switching events was thus increased for intervals spanning an alloHSCT or relapse event, the observed difference was not found to be statistically significant (p = 0.17; Fisher’s exact test). This investigation demonstrates that individual strains can be replaced by other strains of the same species during the course of alloHSCT.

**Figure 8:**
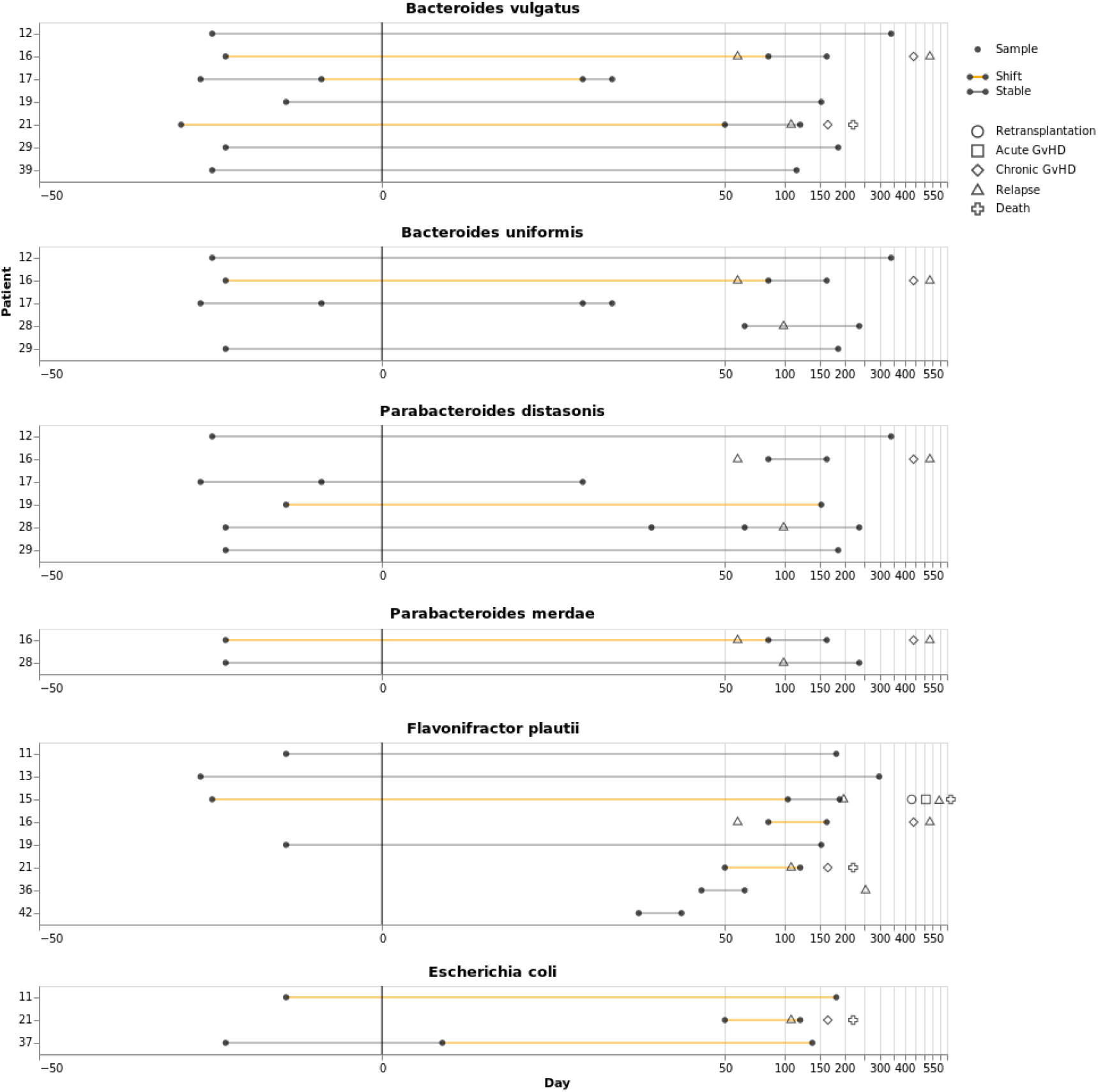
Strain replacement analysis. The Figure visualizes strain dynamics for combinations of bacterial species and sampling timepoints for which sufficient Illumina short-read data was available. Each species is represented by a main panel; main panels are labeled with patient IDs on the y-axis and sampling times on the x-axis. Each dot represents one sampling event, and each horizontal line represents the period between two sampling events. Horizontal lines are colored according to whether a strain replacement event was determined to have taken place between the two sampling events (orange) enclosing the line. Additional symbols indicate adverse events.

## Discussion

Associations between the microbiome and alloHSCT outcomes are incompletely understood, in particular with respect to the role of non-bacterial domains of microbial life; studying these, however, is complicated by limitations of established technologies for microbiome characterization, such as 16S or 18S rDNA sequencing. We thus developed MetaGut, a method based on long-read shotgun metagenomics enabling interrogation of alloHSCT patient microbiomes across almost all domains of microbial life (bacteriome, archaeome, mycobiome, non-human/non-fungal eukaryome, and DNA-virome). MetaGut comprises robust protocols for sample preparation and DNA extraction as well as a specifically developed mapping-based bioinformatics approach to reduce the false-positive taxon detection rate associated with k-mer-based read classification.

We used MetaGut to interrogate patient microbiomes in an explorative clinical study, and found that pre-Tx patient microbiomes fell into 3 clusters (see Figure 4), which can be categorized as “*Bacteroides*- and *Phocaeicola*-dominated” (Cluster 1), “heterogeneous” (Cluster 2), and “*Enterococcus*-dominated” (Cluster 3). The included control samples clustered with the pre-Tx alloHSCT samples of Cluster 1. Interestingly, Cluster 1 exhibited high abundances of *Bacteroides*, often considered an important component of an “intact gut microbiome” and comprising many species known to be commensals and beneficial for human gut health (Zafar and Saier 2021). Cluster 1 could thus be interpreted as least affected by alloHSCT-related microbiome dysbiosis; Clusters 2 and 3, by contrast, may be interpreted as more dysbiotic microbiome states (Ilett et al. 2020). Suggestive differences between the clusters were also detected at the level of viral and fungal microbiome components; for example, *Candida* was detected more often in Cluster 2 and 3 samples, and crAssphage more often in Cluster 1 samples. Consistent with the interpretation of Cluster 1 as least dysbiotic, Cluster 1 also exhibited the lowest rates of antibiotic resistance genes, and patients in Cluster 1 had a lower median number of pre-alloHSCT treatment cycles (median = 2 treatment cycles) compared to patients in Clusters 2 and 3 (combined median = 4.5 treatment cycles; p = 0.01; Mann-Whitney U test). Future studies are required to investigate whether pre-Tx cluster membership is predictive of alloHSCT treatment success since the statistical power necessary to reliably detect such associations cannot be achieved with the 31 patients recruited for our study.

Further applying MetaGut to longitudinally collected samples, we found that patient microbiomes developed dynamically over the course of alloHSCT. Consistent with previous studies (Peled et al. 2017) (Taur et al. 2012), we observed a reduction in microbiome diversity during leukopenia and partial recovery during the reconstitution period. Of note, the leukopenic period was also characterized by a marked increase in fungal abundances (see Figure 3), and expansion of bacterial genera like *Enterococcus* (also observed in (Ilett et al. 2020)) and *Streptococcus.* During the reconstitution period, patient microbiomes collectively showed a return in similarity to pre-Tx states; at the level of individual patient trajectories, however, pre-Tx cluster membership was not predictive of microbiome composition during leukocytopenia (see Figure 5) or reconstitution (see Figure 6). The analyzed reconstitution samples exhibited significant heterogeneity with respect to the time point of sampling (Supplementary Table 3), and individual patients were also represented with multiple samples in this analysis (Supplementary Table 3). It may also reflect the heterogeneity of alloHSCT patient journeys and the significant effects of these on the microbiome (Dethlefsen and Relman 2011). Consistently, the the enriched incidence of bacterial strain replacement around stem cell transplantation and relapse events further emphasizes the potential microbiome disruption in treated patients.

Intriguingly, our study also confirmed the relevance of non-bacterial taxa in the context of alloHSCT. First, we demonstrated robust detection of non-bacterial microbiome components in alloHSCT microbiomes, including, in particular, fungal (*Saccharomyces and Candida*) and viral (crAssphage) taxa. Archaea and non-fungal eukaryotes were also detected, but at lower relative abundances (see Figure 3). In this context, the mapping-based validation component of MetaGut was instrumental in enabling the distinction between confident hits and likely false-positives. Second, while the absolute proportion of non-bacterial taxa was small in most sampled patient microbiomes, viruses and fungi accounted for more than 50% and 10% of total microbial reads in individual microbiomes, respectively. Third, varying overall proportions of non-bacterial abundances, differences between the investigated treatment phases with respect to the validated detection of species at > 5% abundance (in the case of viruses), as well as individual microbiome trajectories indicated that the non-bacterial components of the microbiome also exhibited high inter-patient and temporal plasticity. Interestingly, *Saccharomyces* and *Candida* were detected at high relative abundances throughout the course of alloHSCT, potentially indicating relative stability of these microbiome components over the course of alloHSCT. Of note, in contrast to earlier studies based on 16S or 18S rDNA sequencing, employing an unbiased metagenomic approach, MetaGut enabled the unbiased measurement of the relative quantitative abundances of different types of microbial life. To the best of our knowledge, our study is one of the first few studies to employ shotgun metagenomics in the context of alloHSCT (Ilett et al. 2020, Yan et al. 2022), and, of these, the first to explicitly interrogate the mycobiome and DNA virome. Further studies are required to better characterize the relative stability of the fungal and viral microbiome components and the interplay between these and the bacterial microbiome, for instance at the level of bacteriophage-host relationships (Dutilh et al. 2014).

With respect to MetaGut, potential directions for future work include the further improvement of sample handling and DNA extraction protocols, potentially focused on extracting high-molecular weight DNA while retaining robustness, as well as improvements to the bioinformatics analysis components. In the current implementation of MetaGut (v1.0), significant proportions of reads remained unclassified (Figure 2); furthermore, the observed mapping-based validation rates for archaea, non-fungal eukaryotes, and (although to a lesser extent) fungi suggested substantial rates of residual mis-classification within these taxonomic groupings (Supplementary Figure 3). Classification accuracy may benefit from incorporating gut-specific reference databases (Almeida et al. 2021, Leviatan et al. 2022). Moreover, the currently employed validation approach, based on using the proportion of reads with high-quality pairwise alignments as a proxy for distinguishing between likely true-and false-positive taxon detections, could be complemented with approaches based on horizontal genome coverage (which we explored with promising results for a likely false-positive eukaryotic hit, *Toxoplasma*; Supplementary Note 2), or analysis of k-mer distributions (Breitwieser et al. 2018). It is noteworthy that significant fractions of reads classified as human by Kraken2 did not validate in some samples (Supplementary Figure 3). While we did not attempt an in-depth investigation, an improved understanding of this phenomenon may contribute to further reducing the rate of false-positive read assignments. An important feature of the current version of MetaGut is that the functional analysis of microbiome samples covers the detection of ARGs; future versions of MetaGut could also incorporate analyses of metabolic pathways (Franzosa et al. 2018) and virulence factors (Yan et al. 2022). Now, after successfully having established MetaGut, we can attempt to map established microbiome biomarkers associated with alloHSCT outcomes - such as the quantitative definition of “dominance” by (Taur et al. 2012) as ≥ 30% abundance - onto the microbiome measurements produced by MetaGut in future studies with sufficient patient numbers. As a prerequisite, a future calibration study should address factors such as the utilized extraction protocol, DNA sequencing technologies, and downstream bioinformatics, which influence the accuracy of microbiome measurements (Sinha et al. 2015)). Importantly, in contrast to rDNA amplification-based approaches, MetaGut enables the simultaneous, unbiased and quantitative measurement of human-derived DNA and the exploration of its biomarker potential, e.g., with respect to potential associations between human-derived DNA and mucosal integrity during leukopenia. Conceivably, such analyses could also include cell-of-origin analyses, leveraging the methylation detection capability of the Oxford Nanopore technology (Katsman et al. 2022).

In summary, we have presented MetaGut, a method for characterizing microbiomes simultaneously for their bacteriome, archaeome, mycobiome, non-human/non-fungal eukaryome, and DNA virome composition. Applying MetaGut in an exploratory study to a cohort of alloHSCT patients, we carried out one of the first investigations of the microbiome in the context of alloHSCT based on shotgun metagenomics, confirmed the potential relevance of non-bacterial microbiome components, and identified a 3-cluster structure of pre-Tx patient microbiomes. Furthermore, we could demonstrate that microbial strains can be newly acquired or replaced during the course of alloHCST. These findings can serve as an important basis for future studies, aiming at a more comprehensive characterization of the role of the microbiome in alloHSCT-patients based on shotgun metagenomics.

## Materials and Methods

### Patient recruitment and sampling scheme

Patient recruitment for collection of clinical alloHSCT samples was carried out between June 2018 and December 2019. Participation in the study was offered to all patients undergoing alloHSCT at the Department of Hematology, Immunology, and Clinical Immunology at University Hospital Düsseldorf, according to the following inclusion criteria: persons of legal age that are about to receive an allogeneic stem cell transplantation at the UKD and are able to consent. A full description of the recruited cohort is given in Supplementary Note 1; patient characteristics and treatment histories are summarized in Supplementary Table 2 and Supplementary Table 8. Fecal samples from alloHSCT patients were collected prior to transplantation, during the leukopenic period (defined as white blood count of ≤ 1,000 / µL), and after reconstitution. Stool samples were collected in standard stool collection tubes (feces tubes 76 x 20 mm; Sarstedt), stored at 4°C if possible and transported for further processing to the Institute for Medical Microbiology and Hospital Hygiene of Heinrich Heine University Düsseldorf as fast as possible.

A convenience control cohort of 10 individuals not diagnosed with hematological malignancies was recruited from employees of Düsseldorf University Hospital between February 2021 and June 2021. Control samples were collected from each individual after inclusion into the study and processed using the same protocols also applied to the alloHSCT samples.

### Sample preparation and sequencing

Upon receipt, 1 - 4 aliquots from each stool sample of approximately 1 mL or 1 g of specimen each were placed into 1 - 4 labeled PowerBead Tubes containing 750 µL C1 buffer solution (Qiagen) each.

DNA extraction was carried out using a modified version of the HMP protocol (Anon (n.d.)). Briefly, specimens were homogenized 30 - 40 seconds by vortexing and placed into a heating block at 65°C and 95 °C for 10 minutes respectively. Samples were then stored at - 80°C until DNA extraction. Aliquots for DNA extraction were thawed at 4°C or room temperature (RT) and gently mixed by vortexing. 60 µL of C1 solution was added and briefly vortexed or inverted several times. Aliquots were vortexed at maximum speed for 10 minutes and centrifuged at 10,000 ×g for 30 seconds at RT. From this step onwards, the DNeasy Power Soil Kit was used as per protocol. The concentration of the extracted DNA was determined by fluorometry using the Invitrogen Qubit 4 Fluorometer (Termo Fisher Scientific) and aliquots were stored at −80°C until the Nanopore and Illumina sequencing was performed.

Long-read Nanopore sequencing of all samples was carried out on the PromethION and MinION devices, using FLO-MIN106 R9-Version and FLO-PRO002 R9.4-Version flow cells, the SQK-LSK109 ligation sequencing kit, and the EXP-NBD104 or EXP-NBD114 barcoding kits for barcoded sequencing of native DNA, according to protocol NBE_9065_v109_revAA_14Aug2019. Basecalling was carried out using Guppy.

83 genomic DNA samples, selected to cover at least one pre-post pair per patient, were used for short-read Illumina sequencing. The samples were quantified by fluorometric assay (Qubit DNA HS Assay, Thermo Fisher Scientific) and their quality measured by capillary electrophoresis using the Fragment Analyzer and the ‘DNF-488 HS Genomic DNA Kit’ (Agilent Technologies, Inc.). Although the initial concentration of some samples was low, with some of them not measurable at all, we processed all of them, since the library preparation kit is also suitable for small input quantities. Library preparation was performed using the ‘Nextera XT DNA Library Preparation Kit - Document # 15031942 v05 (Illumina, San Diego, USA.). Depending on whether the concentration of the sample could be determined, either 1 ng gDNA or 5 µL of the sample volume was used for the tagmentation step. Library amplification for final enrichment was performed with 13 cycles. Bead purified libraries were normalized and subsequently sequenced on the HiSeq3000 system (Illumina, San Diego, USA) with a read setup of 2×151 bp. The bcl2fastq2 tool was used to convert the bcl files to fastq files as well for adapter trimming and demultiplexing.

DNA extraction and sequencing data generation metrics are summarized in Supplementary Table 3.

### Analysis of long-read sequencing data

Long-read sequencing data were analyzed using a comprehensive reference database (“MetaGut v. 1.0 database”) comprising 292 giga base pairs of sequence and 1.5 million microbial genomes from all domains of microbial life (Supplementary Table 10). We constructed a custom Kraken2 database by using the Kraken2 build scripts for all domains, modified to download all assemblies marked as “representative genomes” (as opposed to just “Chromosome” or “Full Genome” assemblies) for fungi and protozoa. In addition, we included the *Mus musculus* reference genome (GCF_000001635.27_GRCm39). A full breakdown of database contents is shown in Supplementary File 1. Read classification and compositional analysis were carried out using a 2-step approach that combined an initial k-mer-based read assignment step with mapping-based validation. During the first step, all reads of a sample were classified using Kraken2 (Wood et al. 2019), yielding an initial estimate of sample composition. During the second step, the presence of genera detected during the first step was validated using minimap 2 (Li 2018).

Briefly, to validate the presence of a genus in a sample corresponding to node *n* in the reference database taxonomy, the following steps were carried out: (i) All sample reads assigned to node *n* and its descendants were collected; (ii) a combined reference genome for node *n* was created by linearly concatenating the reference genomes of all leaf-level descendants of node *n*; (iii) the collected reads for node *n* were mapped against the combined reference genome for node *n*; (iv) the proportion of mapping-validated reads was determined, where “mapping-validated” was defined as ≥ 70% of a read being covered by read-to-reference alignments with ≥ 70% identity; (v) finally, the genus corresponding to node *n* was defined as “validated” in a sample if ≥ 30% of the sample reads assigned to *n* achieved mapping-based validation. To increase computational efficiency, validation was carried out on a random subsample of 100,000 reads in each sample (or on the full sample if it contained fewer reads).

Of note, an initial analysis of Zymo Gut Microbiome Standard data (see next section) showed an increased false-positive rate at the species level (see Results); all MetaGut analyses were therefore restricted to the genus level, unless indicated otherwise. Furthermore, due to low per-genus validation rates of archaea and non-fungal eukaryotes (see Results), analyses of the occurrence and abundance of individual genera were limited to the bacterial, viral and fungal components of the microbiome.

The sequencing data - filtered for human DNA - is available at SRA under BioProject Identifier: PRJNA929328.

The bioinformatics component of MetaGut is available under an Open Source license (see “Availability of Code”).

### Determination of read validation thresholds

The utilized read validation thresholds were defined empirically and based on an analysis of the Zymo Gut Microbiome Standard^1^. Briefly, the community standard was long-read-sequenced, employing the same protocols used for fecal samples. The generated reads were classified against the MetaGut default database using Kraken 2, and the assignments of individual reads were validated, using the 70%/70% criterion defined in the previous section, by mapping against (a) a database constructed from the Zymo-provided reference genomes of the species in the Zymo standard, and (b) the MetaGut default reference database.

When using the the Zymo-provided reference genomes for read validation (i.e. representing the assumed case that the genomes of all taxa present in the sample are perfectly represented in the reference database; but see below), per-genus read validation rates varied between 82.4% and 99.3%, with the exception of *Candida, Salmonella*, *Clostridium* and *Enterococcus* (Supplementary Table 11). None of the 7 and 22 reads assigned to *Clostridium* and *Enterococcus*, respectively, could be validated, indicating that these represent false-positives; this is consistent with an expected absolute read count (relative theoretical abundance x number of generated reads) for these genera of 1 and 0, respectively. *Candida and Salmonella* exhibited read validation rates of 49.7% and 73.4%; we hypothesized that these could be explained by a mismatch between the *Candida* and *Salmonella* genome present in the sequenced Zymo sample and the Zymo-provided reference genome. We tested and confirmed this hypothesis by *de novo* assembly of the Zymo sequencing data, followed by mapping of the assembled contigs against the database constructed from the Zymo-provided reference genomes using FastANI (Jain et al. 2018), weighted by assembled contig length; in this analysis, *Candida* and *Salmonella* appeared as clear outliers (FastANI estimates of c. 96%; Supplementary Table 12), indicating genetic divergence.

When using our comprehensive reference database for read validation (i.e. representing the case of potential divergence between the genomes in the sample and the next-closest database genomes) per-genus read validation rates varied between 38% and 98% for genera truly present in the Zymo standard (excluding Enterococcus and Clostridium due to low abundance)), and between 0% and 100% for false-positive genera (Supplementary Table 11). We hypothesized that per-taxon read validation rates would vary with the genetic distance between the sequenced and next-closest database genomes and confirmed this hypothesis by carrying out a FastANI analysis (Pearson’s r = 0.374 for the correlation between read validation rate and genetic distance; p =0.170).

Based on these results, we concluded that pragmatically employing a per-taxon read validation rate threshold of 30% would conservatively enable us to filter out false-positive classification results even in the presence of genetic divergence between the genomes present in the sequenced samples and the genomes present in our database.

### Zymo and LifeLines-DEEP compositional analysis

Zymo community standard long-read sequencing data (see previous section) were mapped to Zymo-provided reference genome using minimap2 and classified using the MetaGut default pipeline and database. Results were analyzed visually and by correlation analysis against the Zymo-provided expected abundance distribution. Whole-genome short-read sequencing data were obtained for 20 randomly selected from the Lifelines-DEEP cohort study^2^, and classified using Kraken 2 against the MetaGut default database.

### Microbiome cluster analysis

To investigate high-level microbiome structures, a PCoA analysis of all samples (alloHSCT and control cohorts), based on Bray-Curtis distance and Kraken 2-derived compositional estimates, was carried out. For the PCoA analysis, no results from the read validation step were taken into account. Microbiome clusters were identified by visual assessment.

### Diversity analysis

Reported numbers of genera were measured after downsampling each sample to the smallest number of reads observed in any sample (n =1029 reads). Reads mapping to any taxon within Chordata or Viridiplantae were ignored. For the diversity analysis, no results from the read validation step were taken into account.

### ARG gene detection

Long-read sequencing data were mapped using minimap2 against the CARD database (Alcock et al. 2020), comprising 2614 antibiotic resistance elements. A read was counted as carrying a specific ARG if a pairwise alignment with length > 500 and < 50 undefined “N” characters was detected between the read and the ARG sequence. The ARG gene detection pipeline was applied to all reads from a sample, independent of other pipeline components.

### crAssphage analysis

Long-read sequencing data were mapped using minimap2 against resolved crAssphage sequences (Gulyaeva et al. 2022) A read was counted as emanating from a crAssphage genome if a pairwise alignment with ≥ 70% of query cover at 70% identity was detected between the read and a crAssphage reference sequence; the fraction of primary alignments divided by the number of query reads was used as an estimate for crAssphage abundance. To identify reads that could be confidently assigned to individual strains, filtering based on mapping quality (MQ ≥ 5) was carried out, enabling assignment of 58% of reads. The crAssphage analysis component was applied to all reads from a sample, independent of other pipeline components.

### Tracking of marker genera

A comprehensive list of bacterial marker genera reported to be associated with alloHSCT outcomes was assembled from the literature (Supplementary Table 9), and complemented with selected fungal and bacterial marker genera detected in this study. The final list of genera is summarized in Supplementary Table 9. Longitudinal abundance dynamics were assessed visually, with confidence intervals in the plots representing binomial fraction confidence intervals.

### Detection of strain replacement events

Analysis of strain dynamics was based on (i) a novel measure (“aANI-lowFreq”) robustly approximating average nucleotide identities between the majority bacterial strain of a species present in a sample *a* and bacterial strains of the same species present in another sample *b* from short-read sequencing data and robust even in the presence of low-frequency strains; (ii) empirical calibration of expected aANI-lowFreq values for samples carrying different dominant bacterial strains using short-read data from different patients, based on the assumptions that (a) bacterial species will vary in their aANI-lowFreq distributions and (b) that different patients will typically be colonized by different bacterial strains; (iii) visual assessment of aANI-lowFreq distances for within-patient longitudinal samples from the same patients; a strain replacement event was assumed to have taken place if the observed within-patient aANI-lowFreq distance falls within the observed between-patient distribution. We note that our approach was optimized for reducing the rate of false-positive strain replacement events, potentially trading off sensitivity against precision. We also note that we experimented with MetaSNV 2 (Van Rossum et al. 2021); however, the subpopulation module failed due to insufficient substructure detection on each species.

aANI-lowFreq is based on counting, over shared regions of species-specific core genomes, the number of bacterial SNVs exclusive to sample *a*, i.e. not also found in sample *b*. As a strain replacement event is most clearly indicated by the presence of a novel majority bacterial strain in *a* that is not present in *b*, *a*-exclusive SNVs were defined as positions that carried an allele with ≥ 50% frequency supported by a minimum of 10 sequencing reads in sample *a* and ≥ 10% frequency in sample *b*; the 10% threshold was chosen to allow for the presence of sequencing errors and mis-aligned reads, and, at the utilized threshold on read coverage (see below), often translated to rejecting alleles that were observed on a single sequencing read on sample *b*. A set of species-specific core genome sequences was obtained from the proGenomes2 representative species level reference database (Mende et al. 2020); a position in one of these core genome sequences was defined as “shared” if both the individual position as well the surrounding core genome reference genome contig across ≥ 80% of its positions achieved ≥ 10x coverage in *a* and *b*. The aANI-lowFreq distance between *a* and *b* was defined as the count of *a*-exclusive SNVs divided by the length of the genomic regions classified as “shared” between *a* and *b*.

aANI-lowFreq was validated using simulation by obtaining 100 randomly selected NCBI RefSeq genomes for *Enterococcus faecium*, *Phocaeicola vulgatus*, *Escherichia coli*, and *Bacteroides uniformis*, as well as 72 genomes for *Lactobacillus gasseri*. For each genome, paired-end 2 x 100bp short-read sequencing data at different coverage levels (5, 10, 20, 50, 200, and 1000) were simulated using dwgsim 0.1.14 (Homer 2022). For each species and combination of coverage levels, we generated 50 random pairs of different genomes as well as 15 random pairs of identical genomes; for each pair, the aANI-lowFreq distance was calculated based on the simulated short reads, and a reference ANI (based on the NCBI RefSeq genome sequences) was obtained using FastANI (Jain et al. 2018). The simulations demonstrated that from genome-wide coverages of ≥ 20x, substantial core genome proportions were classified as “shared” (Supplementary Figure 8), and good approximation of FastANI by aANI-lowFreq, at slight levels of ANI underestimation (demonstrating conservativeness of aANI-lowFreq; Supplementary Figure 9).

For detection of strain replacement events within the alloHSCT cohort, only species and sample pairs with at least 50% of the corresponding core genome were considered, and only species for which at least 10 between-patient aANI-lowFreq distances were available for calibration. For the visual assessment step (Supplementary Figure 10), within-patient and between-patient aANI-lowFreq distances were plotted alongside each other, together with aANI-lowFreq distances from the RefSeq-based simulations, when available.

## Supplementary Information

**Supplementary Figure 1. Composition estimates for the Zymo Gut Microbiome Standard.** Abundance estimation was carried out using Kraken2 and the MetaGut v. 1.0 database, for the Minimap2 estimates we used either the fraction of reads or the fraction of bases while using only primary alignments to the Zymo provided reference sequences. The theoretical composition as reported by Zymo is plotted in the first column.

**Supplementary Figure 2. Composition Estimates for Lifelines Deep Samples.** Abundance estimation was carried out using Kraken2 with the MetaGut v. 1.0 database. The boxplot shows the fraction of *Bacteroides* and *Phocaeicola* for the Lifelines samples and for the control group samples of this study.

**Supplementary Figure 3. Validation rates stratified by sampling time point.** Groups (control, pre-Tx, leukopenia, reconstitution) are the same as in Figure 2. Non-fungal eukaryota excludes chordata and plants. For “binary” we considered a taxon to be validated when more than 20% of the associated reads could be validated. In this case we counted all reads as validated. “For continuous” we report the actual (weighted) validation rates.

**Supplementary Figure 4. Comparison of crAssphage abundance estimations.** Reads were mapped against crAssphage references using minimap2 retaining only primary alignments that aligned at least 70 % of the query with an identity of above 70 %. We used the fraction of primary alignments divided by the number of query reads as a rough estimate for the abundance. For comparison the default abundances provided by Kraken2 and the MetaGut v. 1.0 database are shown.

**Supplementary Figure 5. Distribution of reads across specific crAssphage references.** Barplots show the overall fraction of reads that are mapped to crAssphage. The heatmap is limited to the top 100 crAssphage sequences based on the mean of the assigned fractions and shows only ambiguous assignments (normalized to 100%). Sample-reference pairs with ambiguous assignments are colored gray. Since mapping quality directly corresponds to the uniqueness of each mapping, we used a threshold of mapping quality > 5 to determine reads that can be assigned to a specific reference.

**Supplementary Figure 6. Read migration from Kraken2 to minimap2.** Fraction of Kraken2 taxon assignments for reads that were assigned to crAssphage using minimap2 and a custom database of curated crAssphage references. For visual clarity, only the top 20 taxa are shown.

**Supplementary Figure 7. Marker Genera.** Contains abundance plots per patient for selected marker genera using a binomial distribution to model confidence intervals based on alpha=0.05. Patients are shown with time (days) relative to alloHSCT on the x-axis and all control samples are collectively shown on the right side of the figure. Relative frequency and confidence intervals for α=0.05 of validated reads for the specified marker genera are depicted on the y-axis. Each investigated taxon is shown on a separate page.

**Supplementary Figure 8. Share of Overlap Simulated Genomes ANI.** Shows the share of overlapping sequence for pairs of simulated read sets for five bacterial taxa relative to different coverage values.

**Supplementary Figure 9. aANI-lowFreq distance quality relative to coverage.** The inner x-axis contains the FastANI genome distance and the inner y-axis the calculated aANI-lowFreq distance.

Only distances with a shared overlap of the sequences above 70% are shown which removes all distances with simulated coverage 5 or 10.

**Supplementary Figure 10. Overview of high confidence strains level distances.** Distances for sequential samples within patients as histograms (column a), aANI-lowFreq distances for sequential samples within patients relative to the elapsed time (column b, in days), aANI-lowFreq distances between pre-Tx samples of different patients (column c). In addition we plotted the respective distances from the simulation of *Bacteroides vulgatus* and *Escherichia coli* references (see Methods) and marked the highest observed aANI-lowFreq distance (right border) of the two preceding histograms in the row. Significant strain replacement events are marked with a star.

**Supplementary Table 1. Zymo Abundances.** Theoretical composition of the “Zymo Gut Microbiome Standard” as well as abundances based on Kraken2 mappings to our comprehensive database as well as Minimap2 based mapping to the references provided by Zymo (using one estimate based on read counts and a second estimate based on nucleotide counts).

**Supplementary Table 2. Patient Statistics.** Metadata for each patient identifier. This contains primarily information about clinical outcomes and is used as an input file for the Jupyter Notebook based downstream analysis.

**Supplementary Table 3. Sample Statistics.** Metadata for each sample including sequencing statistics, top-level abundance estimates and cluster association. This is used as an input file for the Jupyter Notebook based downstream analysis and provided for reproducibility.

**Supplementary Table 4. Results Mann Whitney U.** Mann-Whitney U tests analyzing significant differentiation between pre-Tx clusters based on sample statistics

**Supplementary Table 5. Presence_Above_0.05_Bacteria.** Tables listing the presence of validated genera detected at an abundance exceeding 5%, stratified by domain-like groups.

**Supplementary Table 6. Presence_Above_0.05_Fungi.** Tables listing the presence of validated genera detected at an abundance exceeding 5%, stratified by domain-like groups.

**Supplementary Table 7. Presence_Above_0.05_Viruses.** Tables listing the presence of validated genera detected at an abundance exceeding 5%, stratified by domain-like groups.

**Supplementary Table 8. Pre-therapies.** Information about treatments administered to the patient cohort pre-transplantation.

**Supplementary Table 9. Literature List Marker Genera** List of literature references that report findings (regarding HSCT outcomes and IBD) correlated to specific taxonomic entities.

**Supplementary Table 10. Database Overview** Overview of sequences and their length used for building the exhaustive metagenomics Kraken2 database, stratified by domain.

**Supplementary Table 11. Zymo Gut Microbiome Standard Validation** Overview of calibration experiment for the validation method. Contains Kraken2 classifications as well as Minimap2 classifications using the Zymo references and the MetaGut v. 1.0 database respectively.

**Supplementary Table 12. FastANI Assembly Top-Level** Assembled contigs were compared to the Zymo References using FastANI. Each contig was assigned to the reference with the highest ANI. A weighted ANI was calculated based on the contig length for each reference.

**Supplementary Note 1. Cohort Description.** Additional information about the cohort makeup.

**Supplementary Note 2. In-Depth Analysis of Toxoplasma Content** Additional method description and presentation of results for the analysis of Kraken2 based Toxoplasma classifications.

**Supplementary File 1. MetaGut v. 1.0 Database Contents** All contained reference sequences are listed in the Kraken2 report format (Columns in order: Fraction of minimizers at subtree, number of minimizers at subtree, number of minimizers at node, taxonomic level, taxonomic id, taxonomic name).

## Availability of Code

The bioinformatics component of MetaGut was made available as a reproducible Snakemake (Mölder et al. 2021) workflow with the DOI: https://doi.org/10.5281/zenodo.7602236. Each step of the pipeline is executed within its own Conda environment, enabling finely-grained assignment of memory and CPU resources. Downstream analysis was carried out using a set of Jupyter Notebooks (Perez and Granger 2007), also available at the DOI given above.

## Availability of Data

Raw sequencing data and patient metadata are available upon request.

## Ethics Declaration

This study was approved by the ethics committee of the Medical Faculty of Heinrich Heine University Düsseldorf (2019-509).

## Acknowledgments

The study was funded and supported by the Jürgen Manchot Foundation and was part of the project “Decision-making with the help of Artificial Intelligence”. We gratefully acknowledge the support of the medical and nursing staff of the hemato-oncological wards and units at the University Hospital Düsseldorf, without whom this study could not have been carried out. Support extracting and annotating clinical data provided by Sarah Schweier was appreciated. We thank Rebecca Fröhlich for development of the MedReactor tool for visualization of clinical data; Max Ried for computational infrastructure and support, Patrick Finzer, Colin MacKenzie and Tobias Wienemann for valuable discussions. For critical reading of the manuscript we want to acknowledge Sascha Dietrich and Jennifer Jaufmann. Sequencing as well as technical support regarding sequencing hardware were provided by the Genomics and Transcriptomics Laboratory (GTL) of the Biological and Medical Research Centre (BMFZ), Medical Faculty, Heinrich-Heine-University Düsseldorf, Düsseldorf, Germany. Computational support and infrastructure were provided by the “Centre for Information and Media Technology” (ZIM) at the University of Düsseldorf (Germany). We thank the participants and the staff of LifeLines-DEEP for their collaboration. Funding for the project was provided by the Top Institute Food and Nutrition Wageningen grant GH001. The sequencing was carried out in collaboration with the Broad Institute.

## Author Contributions

Project conceptualization: A.D., K.P., R.H.

Wetlab protocol adaptation + validation: B.H., S.S.

Acquisition, analysis, or interpretation of data: A.D., A.R., B.H., G.K., K.P., P.S., R.H., S.S.

Bioinformatics method development: A.D., P.S.

Software development and validation: P.S.

Sample acquisition coordination and sequencing: A.R., S.S.

Visualization of results: A.R., P.S., S.S.

Supervision: A.D., B.H., G.K., K.P., R.H.

Clinical coordination: P.J.

Initial draft of manuscript: A.D., A.R., K.P., P.S., S.S.

Draft revision: A.D., G.K., K.P., R.H.

Funding acquisition: A.D., G.K., K.P., R.H.

All authors read, edited and approved the final manuscript.

## Footnotes

1 https://www.zymoresearch.de/products/zymobiomics-gut-microbiome-standard

2 http://www.lifelines.nl

