## Supplementary Figure 7 for "MetaGut: Insights into gut microbiomes in stem cell transplantation by comprehensive shotgun long-read sequencing"

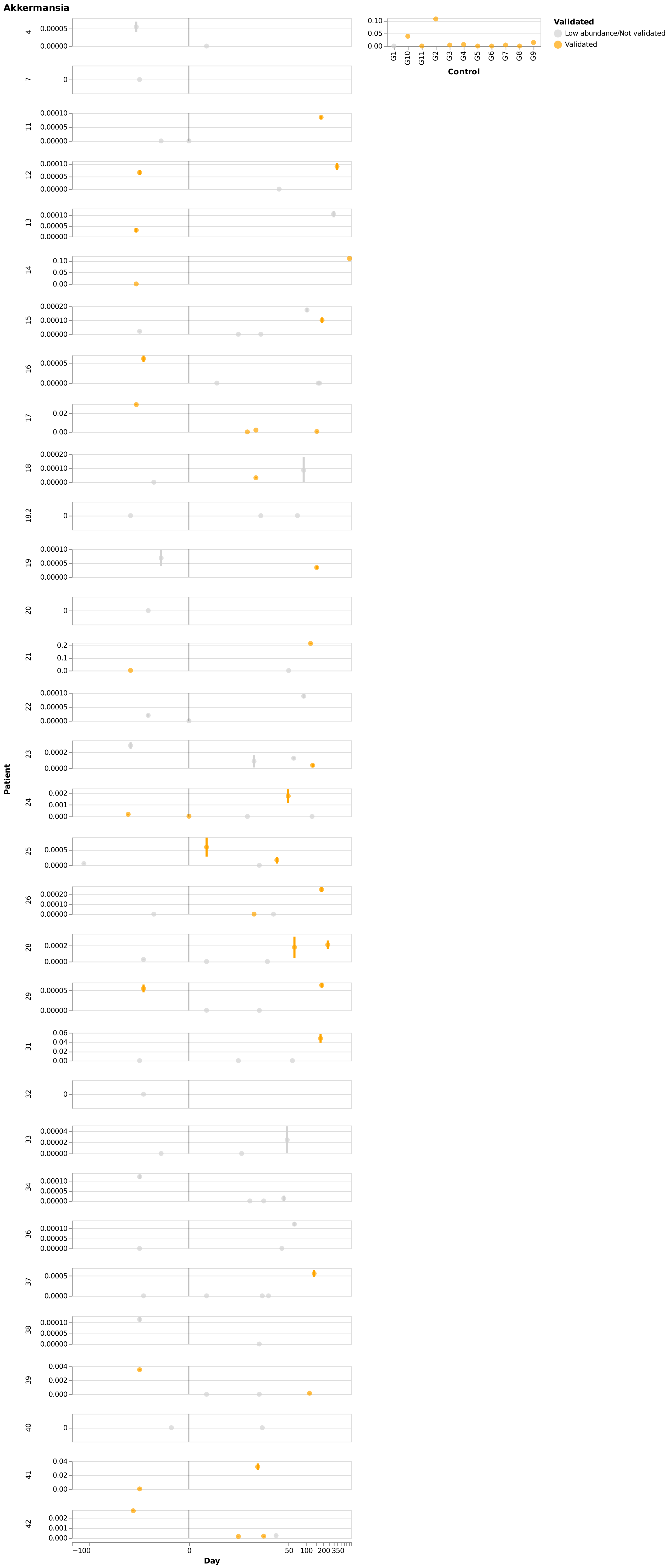
Akkermansia
Validated
Low abundance/Not validated
Validated
G1
G10
G11
G2
G3
G4
G5
G6
G7
G8
G9
Control
Patient
4
7
11
12
13
14
15
16
17
18
18.2
19
20
21
22
23
24
25
26
28
29
31
32
33
34
36
37
38
39
40
41
42
−100
0
50
100
200
350
Day

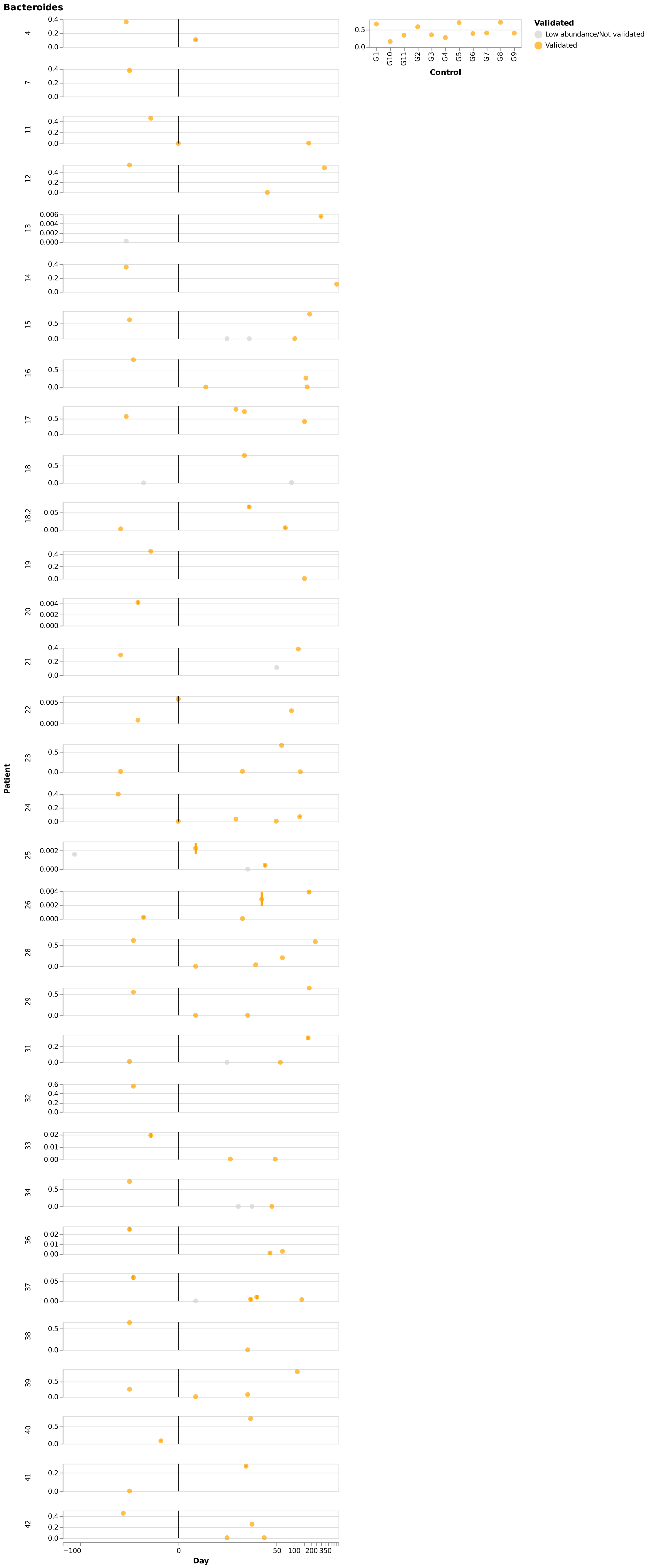

Bacteroides
Validated
Low abundance/Not validated
Validated
Control
G1
G10
G11
G2
G3
G4
G5
G6
G7
G8
G9
Patient
4
7
11
12
13
14
15
16
17
18
18.2
19
20
21
22
23
24
25
26
28
29
31
32
33
34
36
37
38
39
40
41
42
Day
−100
0
50
100
200
350

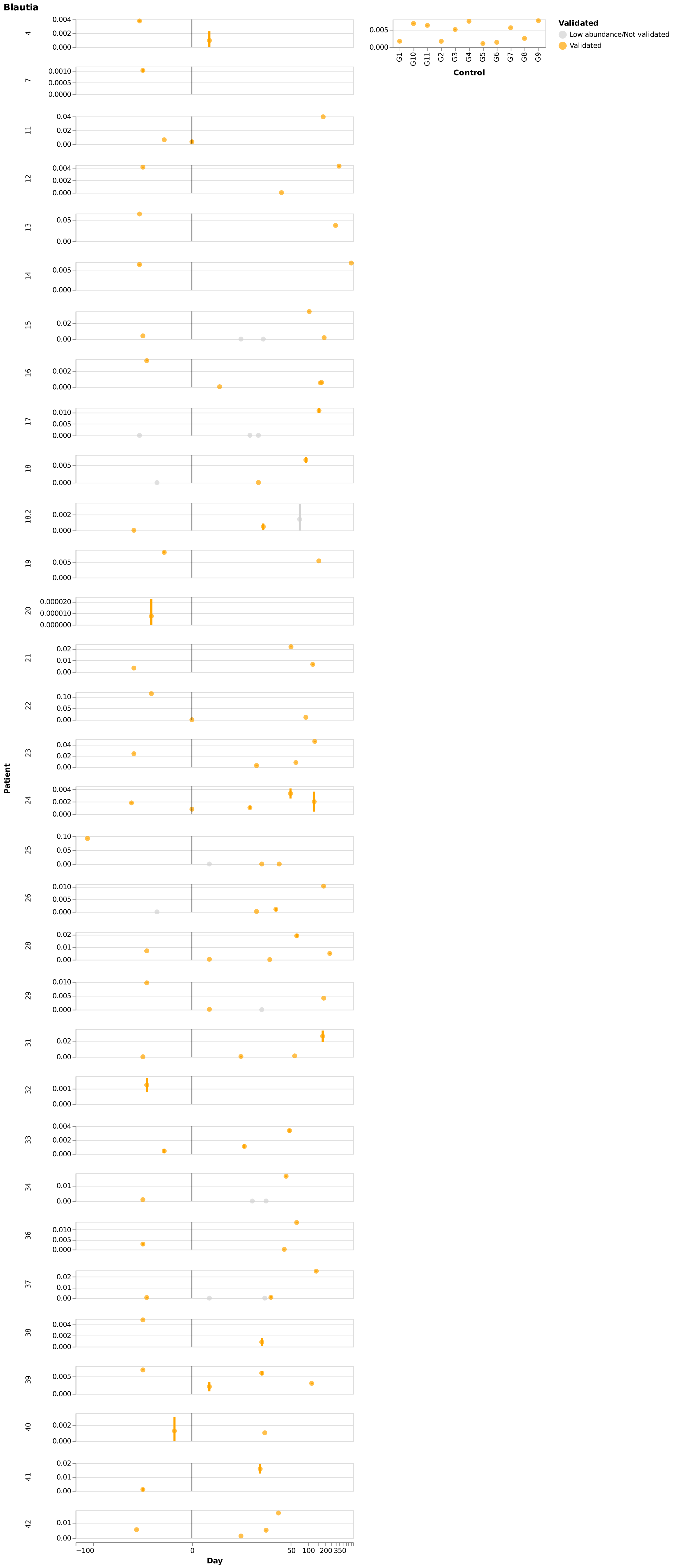

Blautia
Patient
Day
Control
Validated
Low abundance/Not validated
Validated
G1
G10
G11
G2
G3
G4
G5
G6
G7
G8
G9
4
7
11
12
13
14
15
16
17
18
18.2
19
20
21
22
23
24
25
26
28
29
31
32
33
34
36
37
38
39
40
41
42
−100
0
50
100
200
350

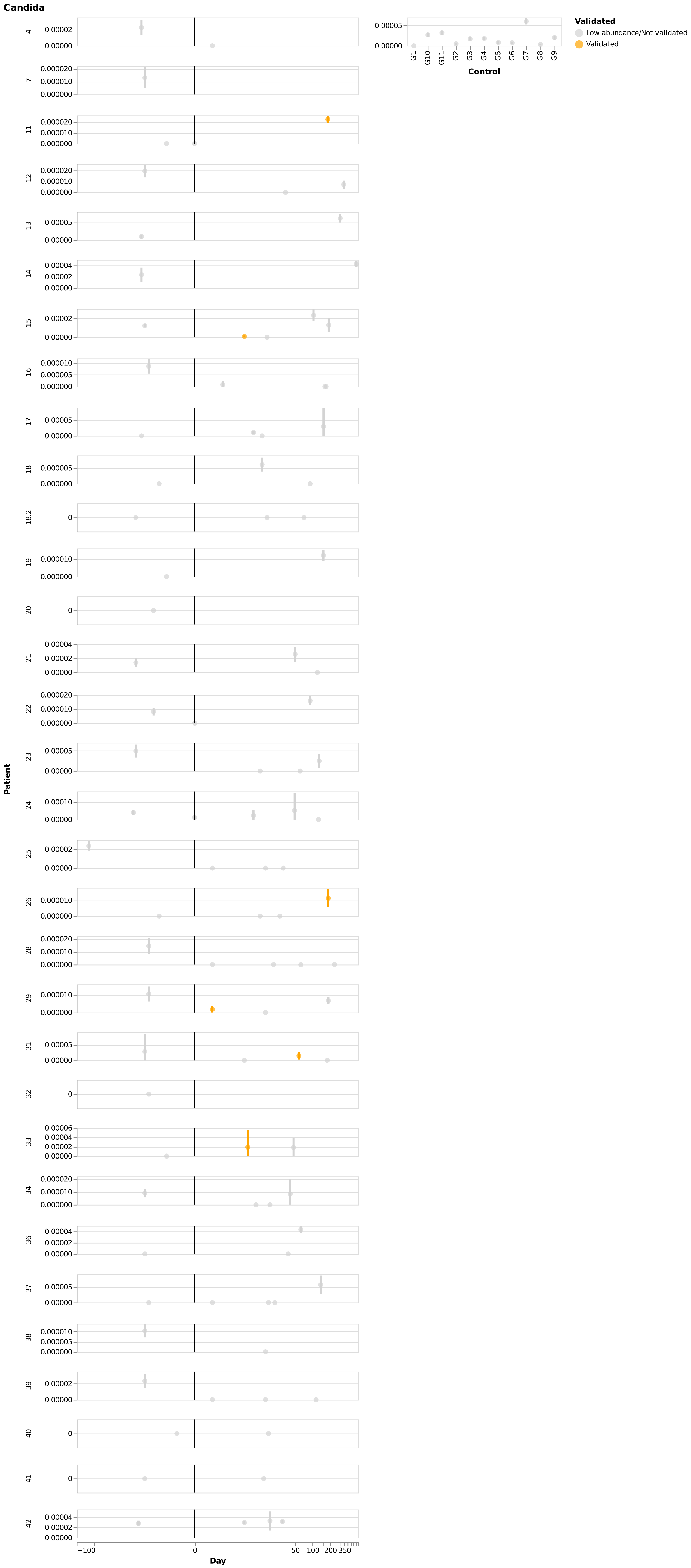

Candida
Validated
Low abundance/Not validated
Validated
Control
G1
G10
G11
G2
G3
G4
G5
G6
G7
G8
G9
Patient
Day
−100
0
50
100
200
350
4
7
11
12
13
14
15
16
17
18
18.2
19
20
21
22
23
24
25
26
28
29
31
32
33
34
36
37
38
39
40
41
42

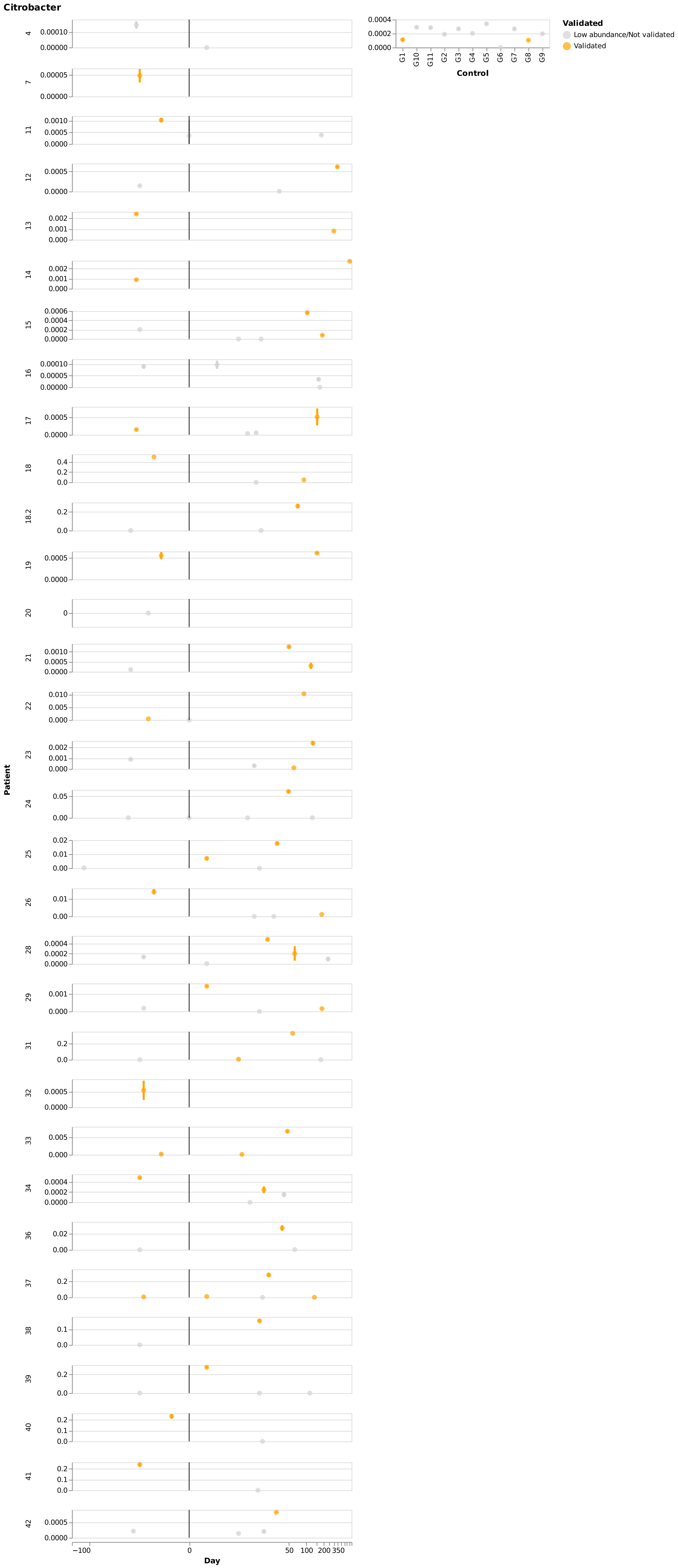

Citrobacter
Validated
Low abundance/Not validated
Validated
Control
Patient
Day

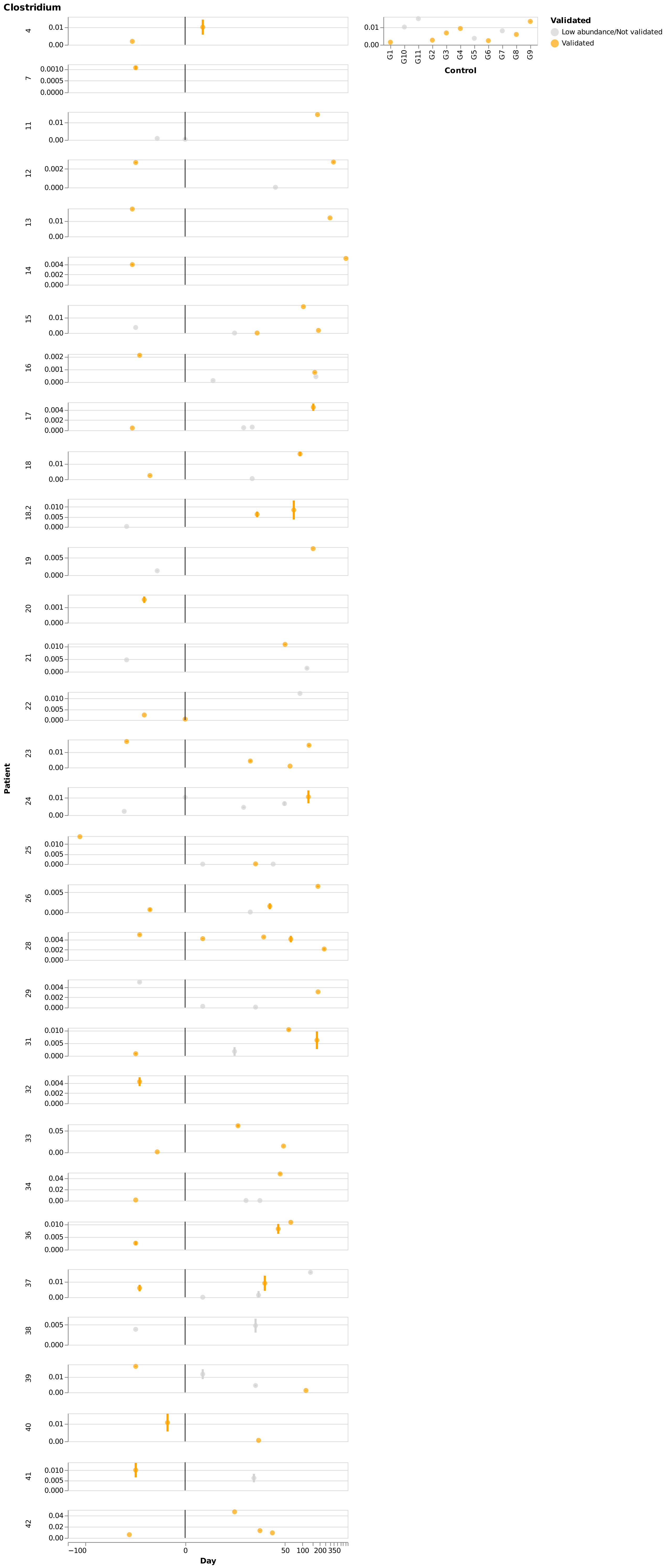

Clostridium
Validated
Low abundance/Not validated
Validated
Control
G1
G10
G11
G2
G3
G4
G5
G6
G7
G8
G9
Patient
4
7
11
12
13
14
15
16
17
18
18.2
19
20
21
22
23
24
25
26
28
29
31
32
33
34
36
37
38
39
40
41
42
Day
−100
0
50
100
200
350

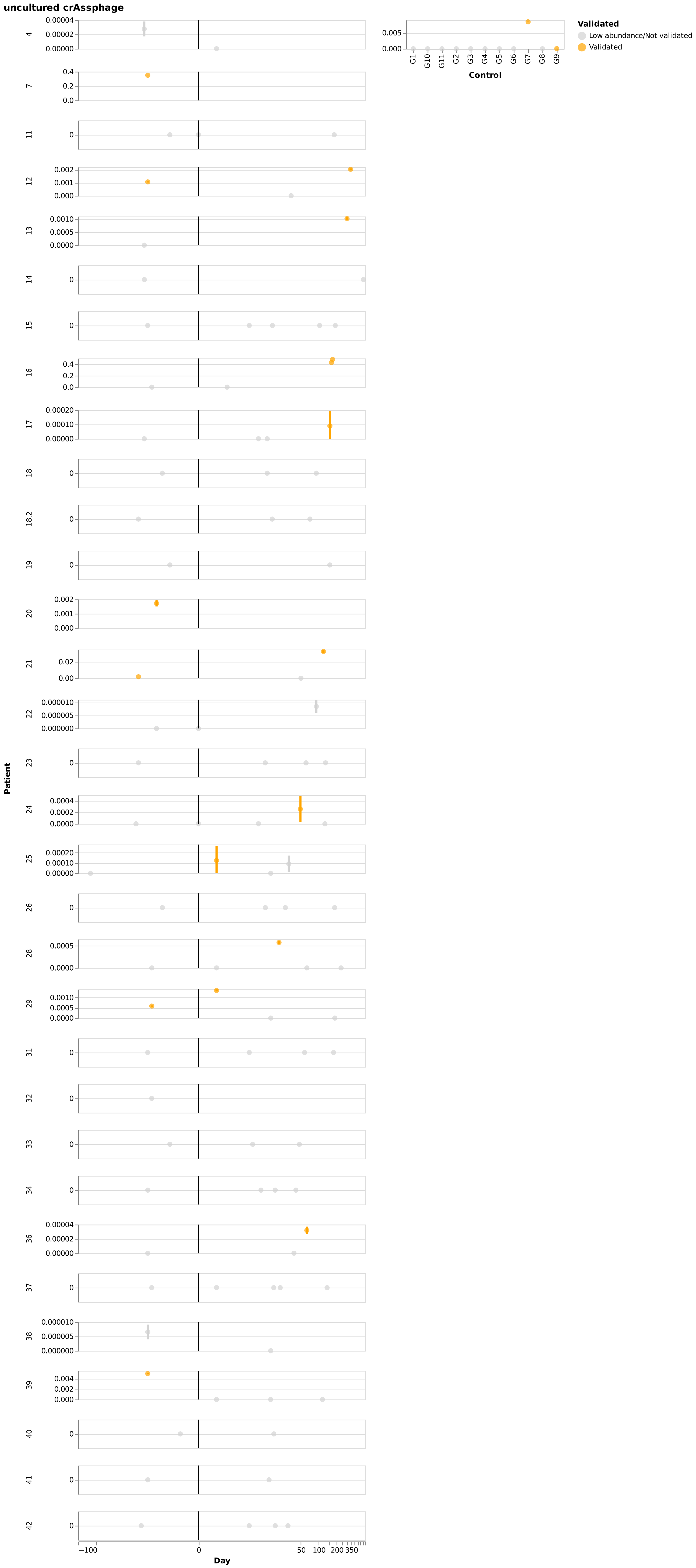

uncultured crAssphage
Validated
Low abundance/Not validated
Validated
Control
G1
G10
G11
G2
G3
G4
G5
G6
G7
G8
G9
Patient
4
7
11
12
13
14
15
16
17
18
18.2
19
20
21
22
23
24
25
26
28
29
31
32
33
34
36
37
38
39
40
41
42
Day
−100
0
50
100
200
350

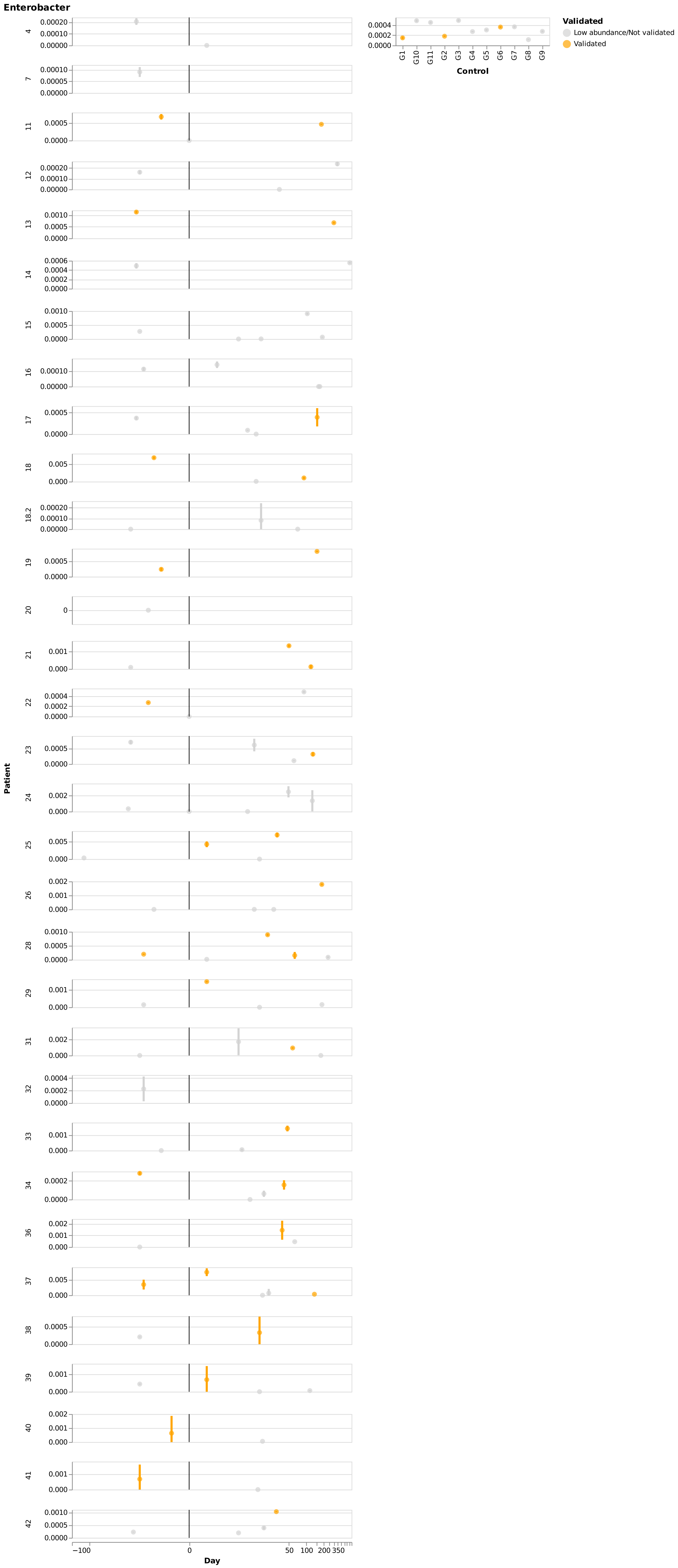

Enterobacter
Validated
Low abundance/Not validated
Validated
Control
G1 G10 G11 G2 G3 G4 G5 G6 G7 G8 G9
Patient
4
7
11
12
13
14
15
16
17
18
18.2
19
20
21
22
23
24
25
26
28
29
31
32
33
34
36
37
38
39
40
41
42
Day
−100 0 50 100 200 350

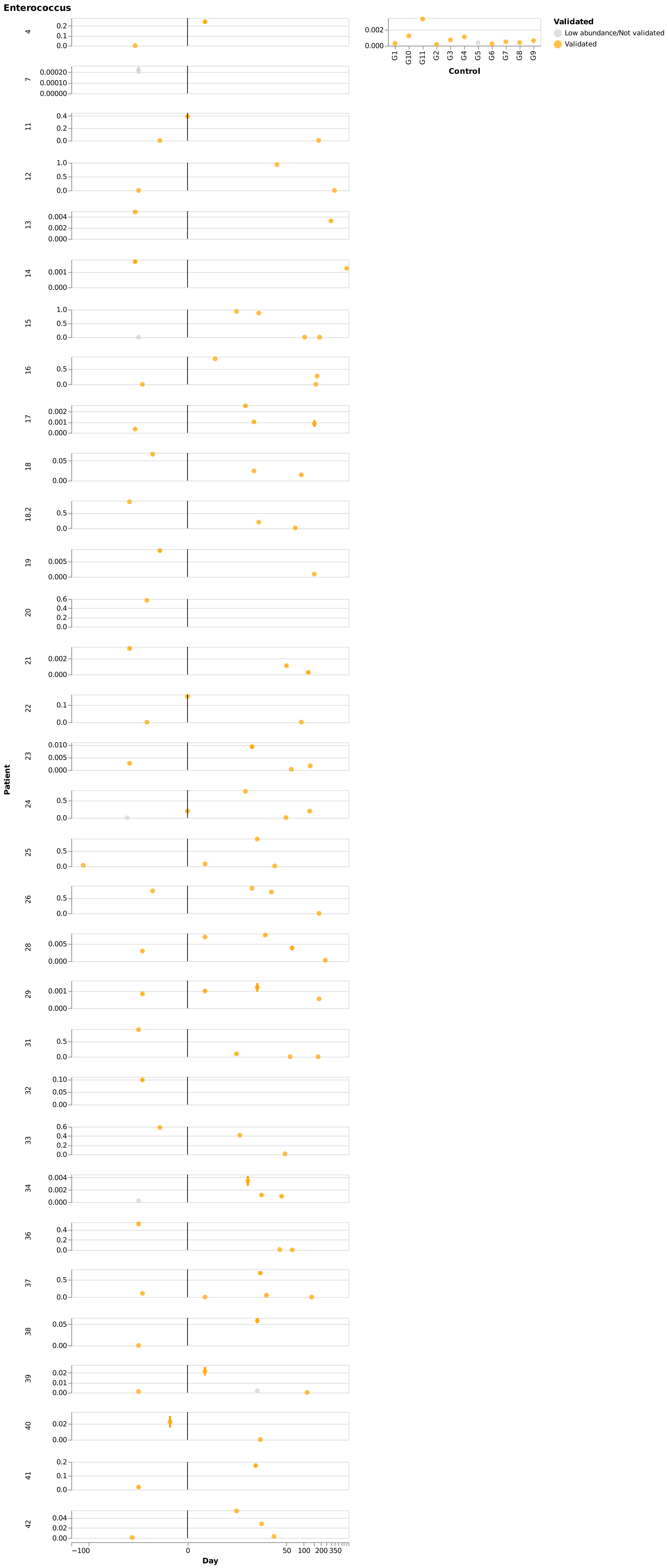

Enterococcus
Patient
Day
Control
Validated
Low abundance/Not validated
Validated
G1
G10
G11
G2
G3
G4
G5
G6
G7
G8
G9
4
7
11
12
13
14
15
16
17
18
18.2
19
20
21
22
23
24
25
26
28
29
31
32
33
34
36
37
38
39
40
41
42
−100
0
50
100
200
350

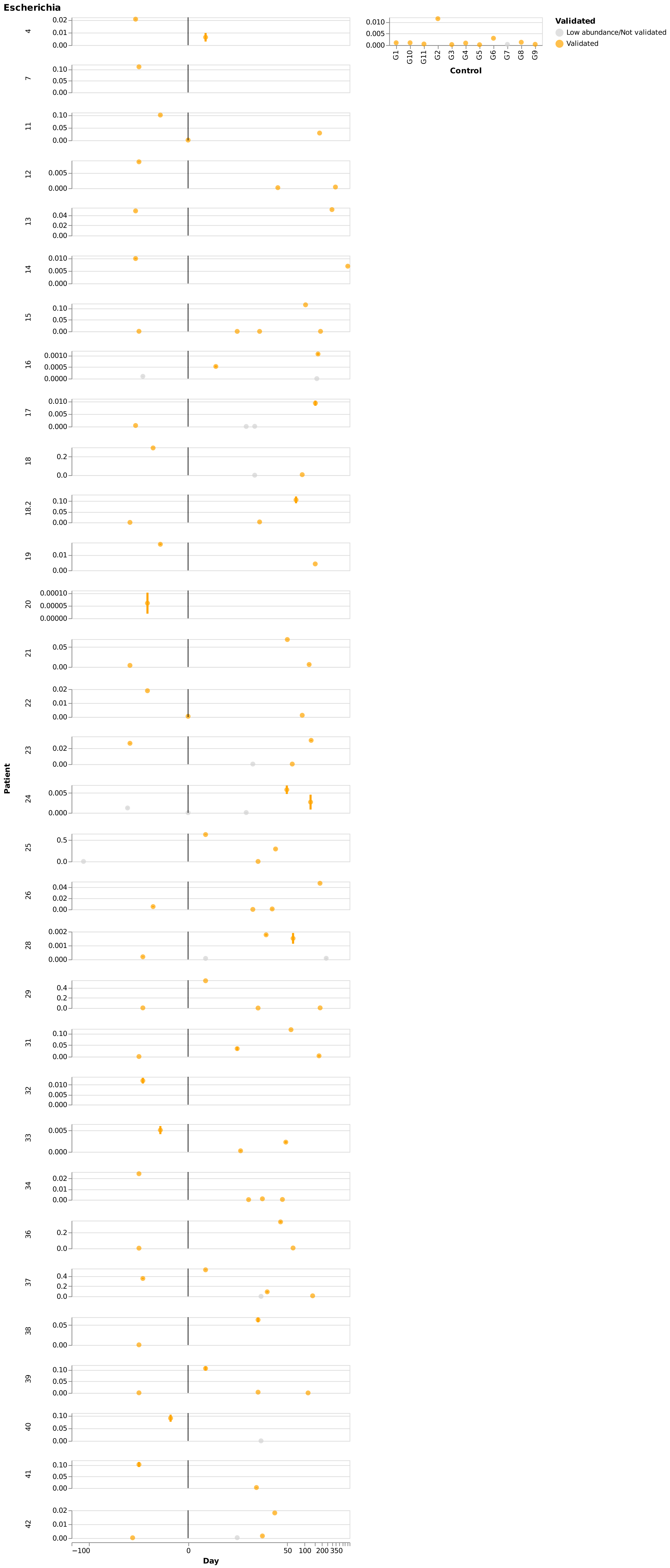

Escherichia
Validated
Low abundance/Not validated
Validated
Control
G1
G10
G11
G2
G3
G4
G5
G6
G7
G8
G9
Patient
4
7
11
12
13
14
15
16
17
18
18.2
19
20
21
22
23
24
25
26
28
29
31
32
33
34
36
37
38
39
40
41
42
Day
−100
0
50
100
200
350

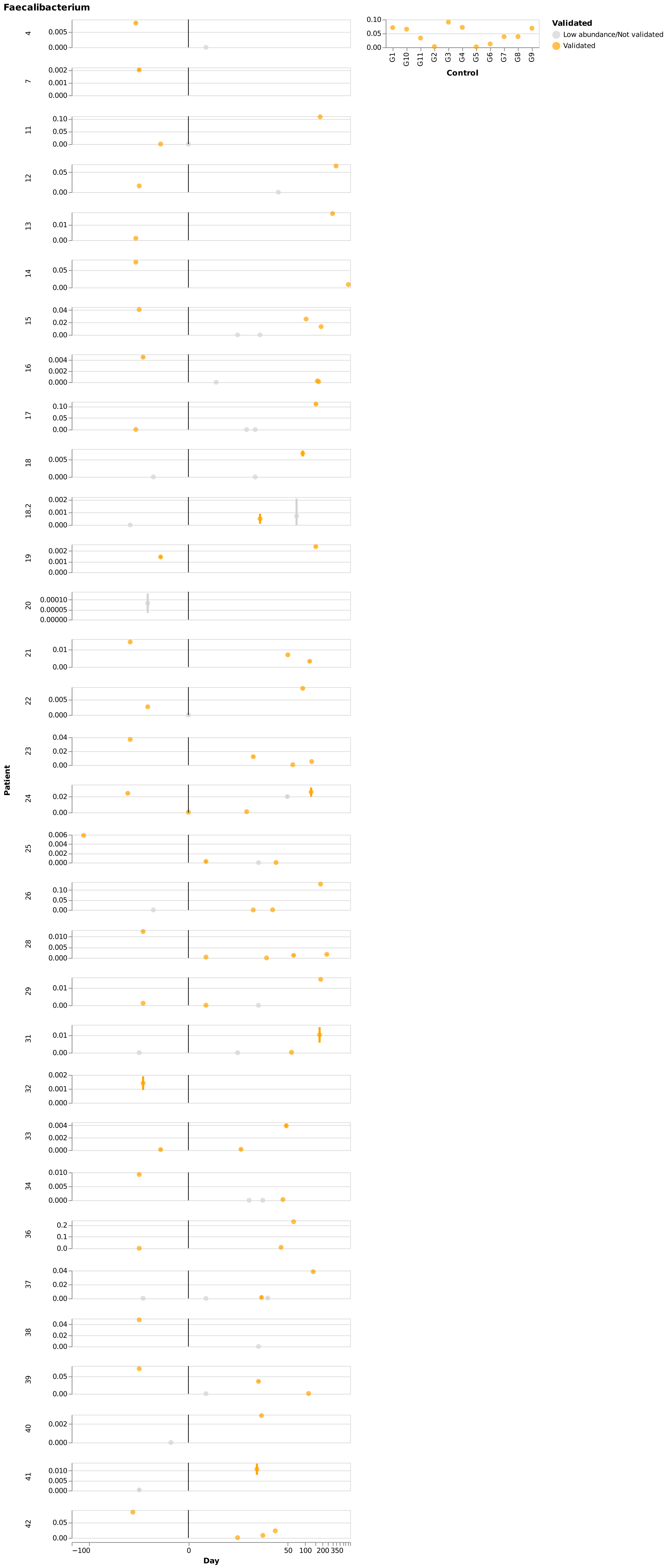
Faecalibacterium
Validated
Low abundance/Not validated
Validated
Control
G1 G10 G11 G2 G3 G4 G5 G6 G7 G8 G9
Patient
4 7 11 12 13 14 15 16 17 18 18.2 19 20 21 22 23 24 25 26 28 29 31 32 33 34 36 37 38 39 40 41 42
Day
−100 0 50 100 200 350

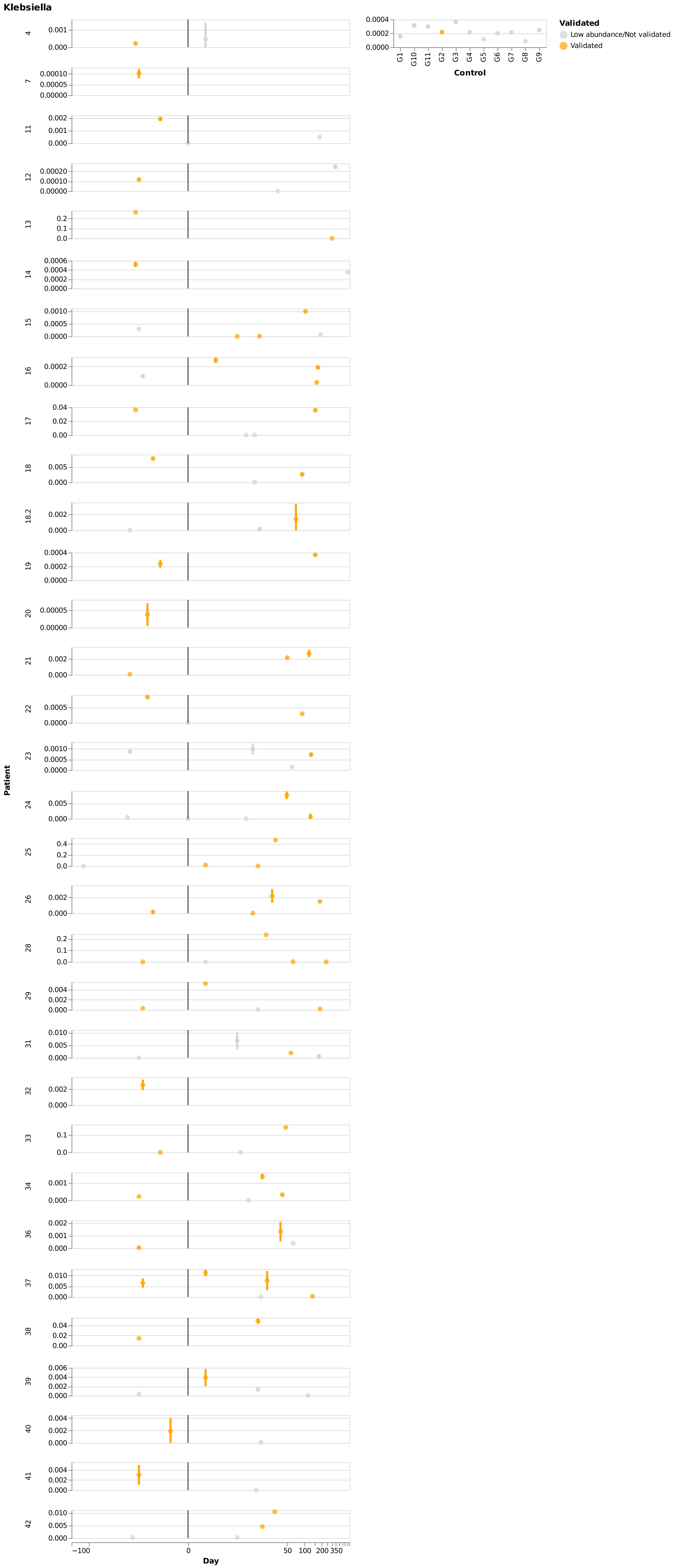

Klebsiella
Validated
Low abundance/Not validated
Validated
Control
G1
G10
G11
G2
G3
G4
G5
G6
G7
G8
G9
Patient
4
7
11
12
13
14
15
16
17
18
18.2
19
20
21
22
23
24
25
26
28
29
31
32
33
34
36
37
38
39
40
41
42
Day
−100
0
50
100
200
350

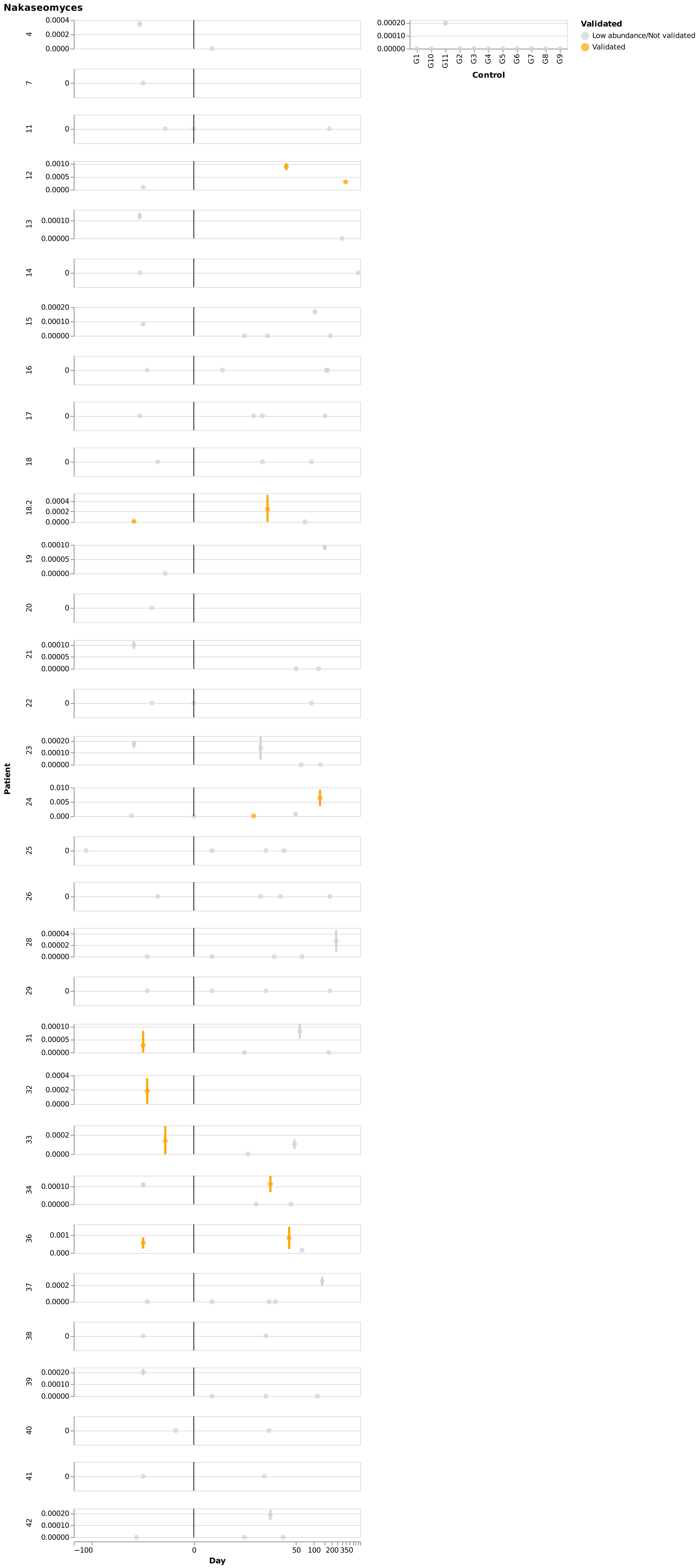
Nakaseomyces
Validated
Low abundance/Not validated
Validated
Control
G1 G10 G11 G2 G3 G4 G5 G6 G7 G8 G9
Patient
4 7 11 12 13 14 15 16 17 18 18.2 19 20 21 22 23 24 25 26 28 29 31 32 33 34 36 37 38 39 40 41 42
Day
−100 0 50 100 200 350

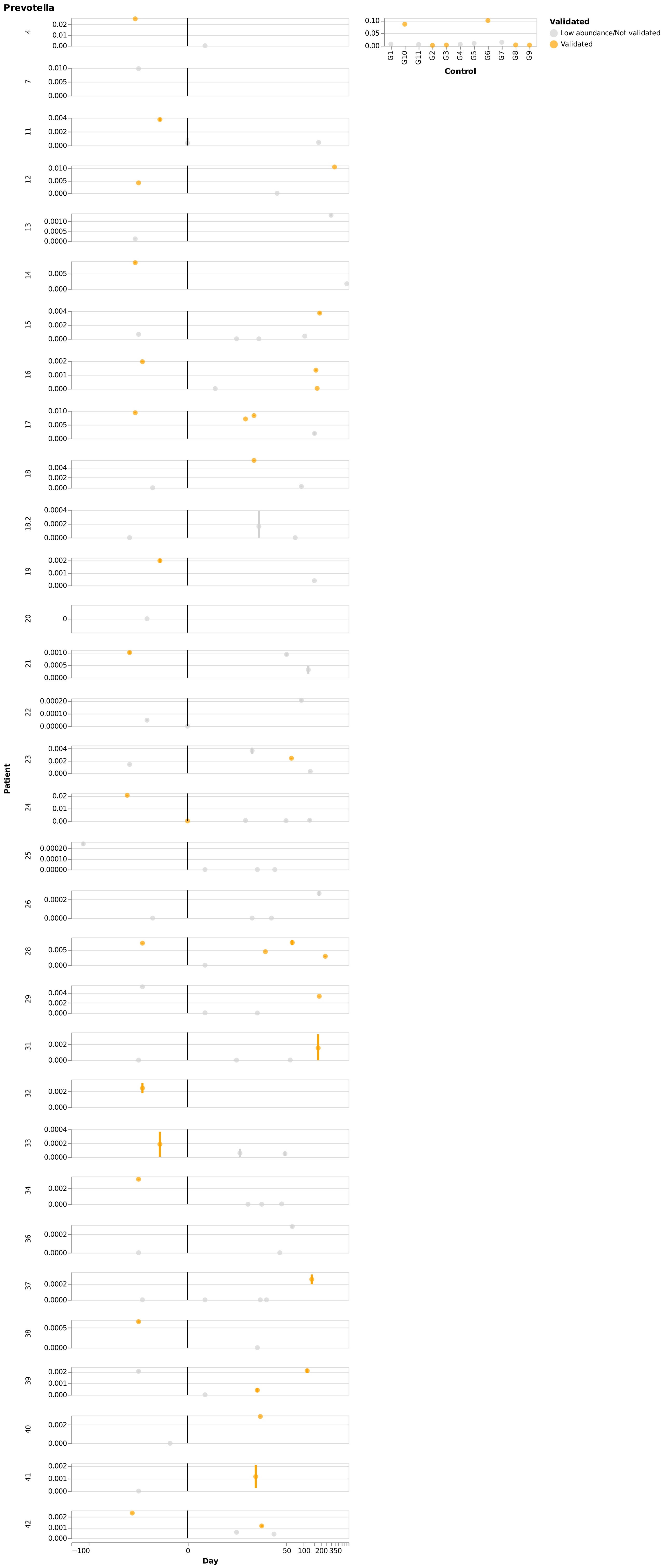

Prevotella
Validated
Low abundance/Not validated
Validated
G1
G10
G11
G2
G3
G4
G5
G6
G7
G8
G9
Control
Patient
4
7
11
12
13
14
15
16
17
18
18.2
19
20
21
22
23
24
25
26
28
29
31
32
33
34
36
37
38
39
40
41
42
−100
0
50
100
200
350
Day

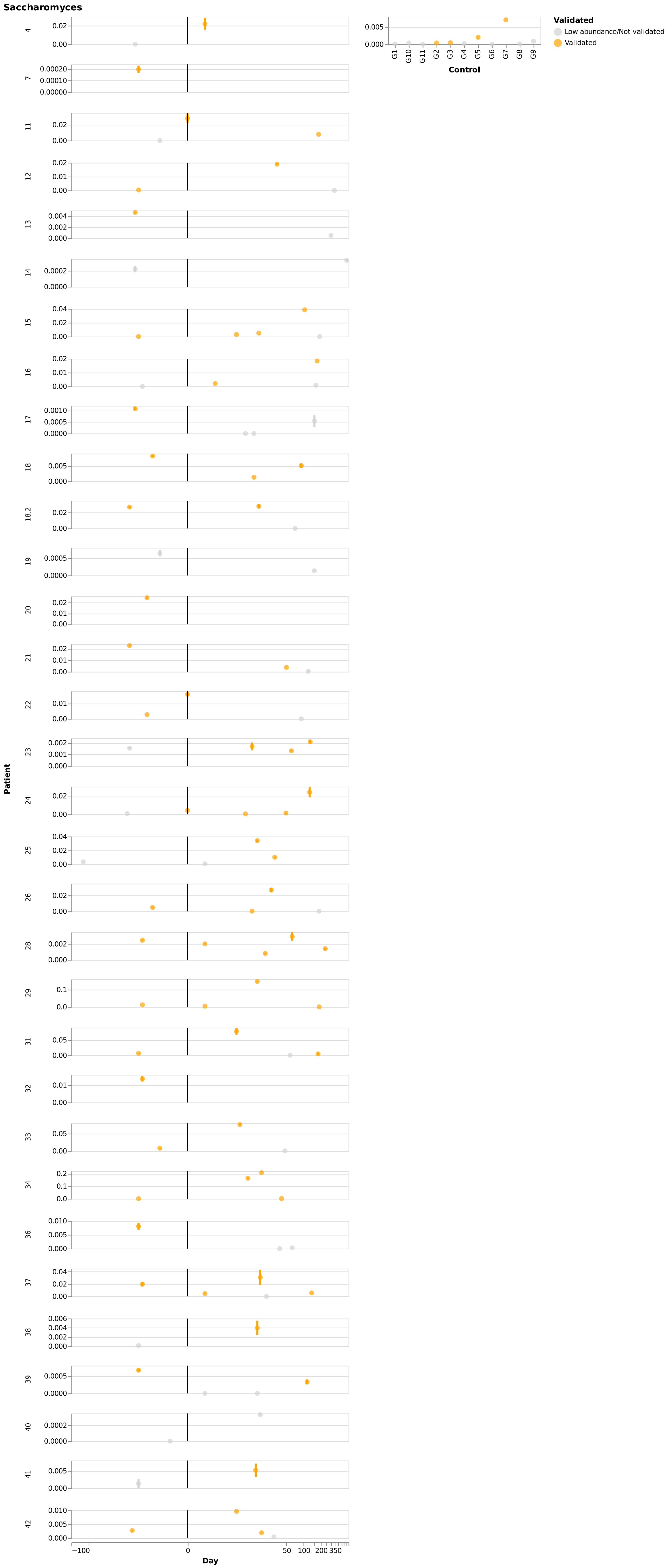
Saccharomyces
Validated
Low abundance/Not validated
Validated
Control
G1
G10
G11
G2
G3
G4
G5
G6
G7
G8
G9
Patient
4
7
11
12
13
14
15
16
17
18
18.2
19
20
21
22
23
24
25
26
28
29
31
32
33
34
36
37
38
39
40
41
42
Day
−100
0
50
100
200
350

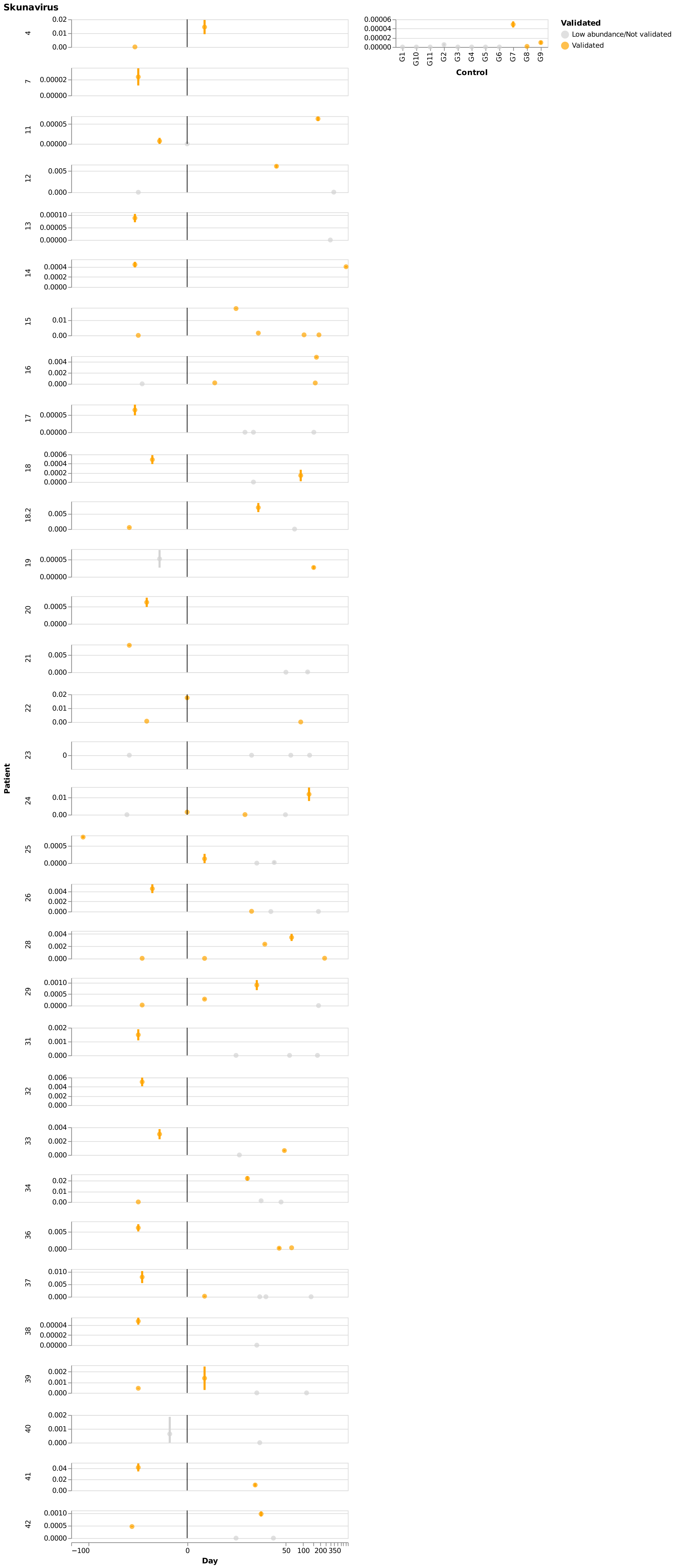
Skunavirus
Validated
Low abundance/Not validated
Validated
Control
G1
G10
G11
G2
G3
G4
G5
G6
G7
G8
G9
Patient
4
7
11
12
13
14
15
16
17
18
18.2
19
20
21
22
23
24
25
26
28
29
31
32
33
34
36
37
38
39
40
41
42
Day
−100
0
50
100
200
350

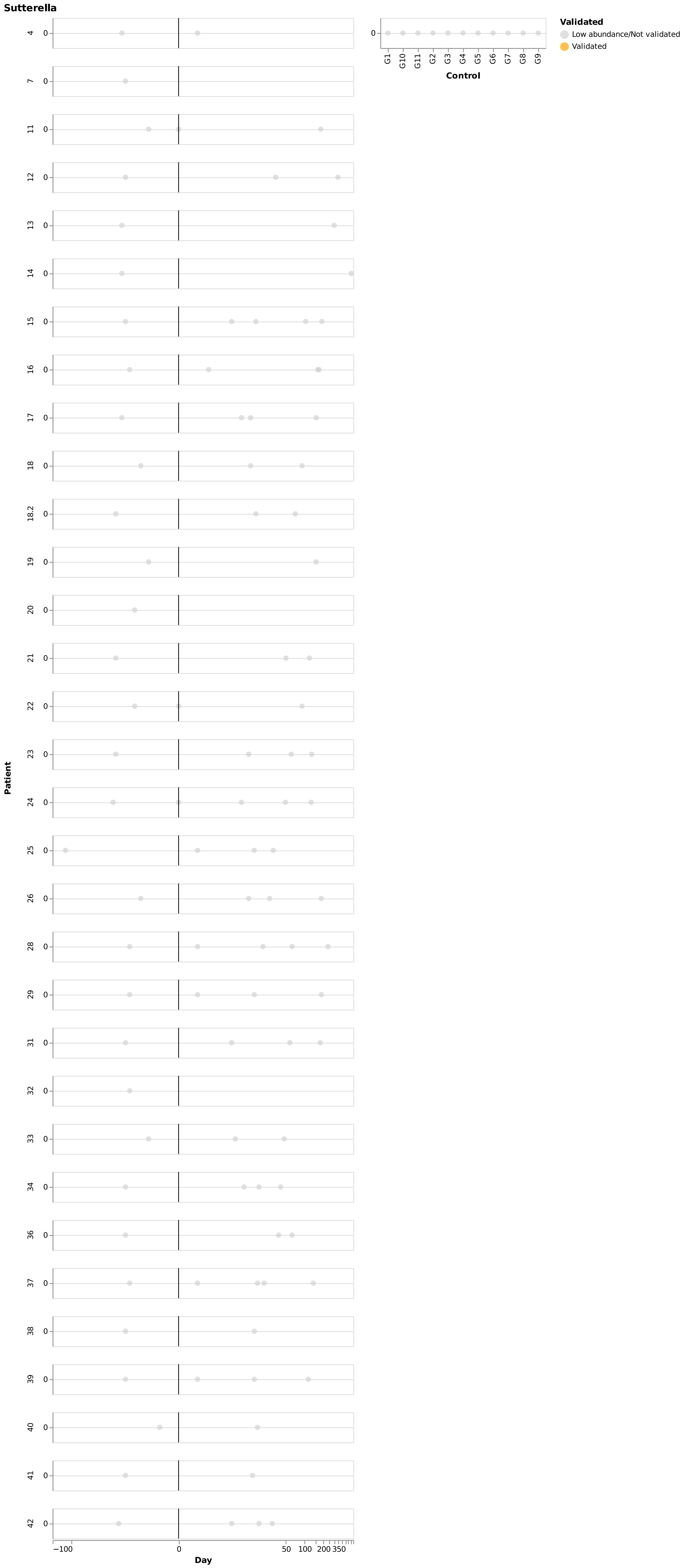
Sutterella
Validated
Low abundance/Not validated
Validated
Control
G1
G10
G11
G2
G3
G4
G5
G6
G7
G8
G9
Patient
4
7
11
12
13
14
15
16
17
18
18.2
19
20
21
22
23
24
25
26
28
29
31
32
33
34
36
37
38
39
40
41
42
Day
−100
0
50
100
200
350
