## Supplementary Note 1 for "MetaGut: Insights into gut microbiomes in stem cell transplantation by comprehensive shotgun long-read sequencing"

### Description of the recruited aHSCT cohort

Between June 2018 and December 2019, 31 patients (n = 12 female and 19 male) undergoing aHSCT at the Department of Hematology, Immunology, and Clinical Immunology of University Hospital Düsseldorf were enrolled into this study. Full patient characteristics, including diagnosis, therapy, demographic data, and HLA matching status are summarized in Supplementary Table 2. Patient Statistics.

The median age of the patients was 58,5 years (range: 19-73 years). Twenty-three patients had myeloid stem cell disease (acute myeloid leukemia (AML; n = 23) or high-risk (MDS; n = 4)), while the remaining 4 patients had lymphoid malignancies. Before initiation of high-dose myeloablative therapy, 12 and 2 patients were in complete and partial remission (CR/PR), respectively, while 17 patients had refractory or progressive disease. This reflects a proportion of 55% patients who presented with active disease at the time of conditioning therapy, partially explained by the fact that 7 patients had received the conditioning therapy without prior induction treatment. Details on the type of conditioning therapy, which depended on disease entity and remission status at the time of conditioning therapy, are given in Supplementary Table 2. Patient Statistics. With the exception of 5 patients, the donors were 10/10 HLA-identical. Of the donors, 5 were siblings and the remaining 26 unrelated donors. Anti-infective prophylaxis during conditioning therapy and after allogeneic transplantation consisted of administration of Ciprofloxacin, Posaconazol, and Amphotericin B irrigation solution. When body temperatures >38.5 °C occurred, empiric antibiotic therapy with Piperacillin or Meropenem in case of Penicillin allergy was started. In case of persistent fever, escalation was performed by switching to Meropenem after initial administration of Piperacillin as well as addition of Vancomycin or Teicoplanin or Tigecycline. Systemic therapy with Caspofungin was started in case of persistence and/or radiological signs of fungal pneumonia. For prophylaxis of graft-versus-host disease patients received the following regimens: Platelet concentrates were administered when the platelet concentration was below 20,000/µL, and erythrocyte concentrates were transfused when the hemoglobin concentration was below 8.0 g/dL.

The median follow-up of the included patients was 531 days (range: 27 - 911; Supplementary Table 2. Patient Statistics). For the main scientific purpose of our study, 29 patients (12 female / 19 male) were fully evaluable. Two patients died of transplantation-associated complications on day 22 and 32 post-aHSCT. Hematologic reconstitution with a white blood cell count (WBC) of > 1,000 /µL was observed after 11 days (median / range: 9 - 20 days). 12 patients suffered from relapse during the study period, from which 6 patients responded well to relapse therapy and went into remission again.
