## Supplementary Note 2 for "MetaGut: Insights into gut microbiomes in stem cell transplantation by comprehensive shotgun long-read sequencing"

### In-Depth Analysis of Toxoplasma Content

Applying our validation scheme we found some samples with validated Toxoplasma content but without clinical indication for Toxoplasmosis. We therefore assessed the alignments in mode detail and found that the references for Toxoplasma Gondii consisted of one large reference genome and multiple smaller contigs of length < 1000 bp. The majority of reads in samples that passed the validation were assigned to those references leading to an implausible coverage distribution with the large reference having almost no reads assigned to it. This has implications for future reference database preprocessing as well as classification approaches. We believe that the reads are unlikely to be real Toxoplasma DNA and identify this as a problem that needs to be resolved for future work.

| Reference Name | Reference Length | Aligned Reads | Aligned Bases |
|---|---|---|---|
| kraken:taxid\|508771\|NW_017384238.1 | 599 | 564 | 829608 |
| kraken:taxid\|508771\|NW_017384541.1 | 424 | 410 | 225024 |
| kraken:taxid\|508771\|NW_017384091.1 | 690 | 245 | 441049 |
| kraken:taxid\|508771\|NW_017384622.1 | 511 | 216 | 130475 |
| kraken:taxid\|508771\|NW_017383938.1 | 395 | 130 | 21992 |
| kraken:taxid\|508771\|NW_017384910.1 | 411 | 108 | 22319 |
| kraken:taxid\|508771\|NW_017384750.1 | 442 | 84 | 37128 |
| kraken:taxid\|508771\|NW_017384921.1 | 567 | 56 | 16477 |
| kraken:taxid\|508771\|NW_017385017.1 | 498 | 41 | 8750 |
| no reference | 0 | 40 | 0 |
| kraken:taxid\|508771\|NW_017384151.1 | 417 | 33 | 12005 |
| kraken:taxid\|508771\|NW_017384026.1 | 499 | 25 | 12014 |
| kraken:taxid\|508771\|NW_017385066.1 | 442 | 22 | 9099 |
| kraken:taxid\|508771\|NW_017384310.1 | 1438 | 17 | 8716 |
| kraken:taxid\|508771\|NW_017384196.1 | 502 | 14 | 3726 |
| kraken:taxid\|508771\|NW_017384311.1 | 809 | 11 | 8603 |
| kraken:taxid\|508771\|NW_017384568.1 | 520 | 8 | 2245 |
| kraken:taxid\|508771\|NW_017384545.1 | 754 | 7 | 3851 |
| kraken:taxid\|508771\|NW_017384808.1 | 359 | 6 | 798 |
| kraken:taxid\|508771\|NW_017384912.1 | 944 | 6 | 1090 |
| kraken:taxid\|508771\|NW_017384082.1 | 581 | 4 | 1328 |
| kraken:taxid\|508771\|NW_017385060.1 | 940 | 4 | 2468 |
| kraken:taxid\|508771\|NW_017385061.1 | 646 | 4 | 1798 |
| *kraken:taxid\|508771\|NC_031476.1* | *6970285* | *2* | *1116* |
| kraken:taxid\|508771\|NW_017384849.1 | 424 | 1 | 149 |
| kraken:taxid\|508771\|NW_017384305.1 | 740 | 1 | 254 |
| kraken:taxid\|508771\|NW_017384721.1 | 645 | 1 | 196 |
| kraken:taxid\|508771\|NW_017384086.1 | 880 | 1 | 790 |

Table: **Alignment distribution for reads classified as Toxoplasma in samples where they were labeled as validated** “Aligned Bases” reflects the number of query bases aligned for a given alignment. The table shows all reads in validated samples even if the individual reads did not pass the validation criteria.
