## Supplementary Table 1 for "MetaGut: Insights into gut microbiomes in stem cell transplantation by comprehensive shotgun long-read sequencing"

| | Zymo Theoretical Composition | Minimap2 Read-Based Estimate | Minimap2 Base-Based Estimate | Kraken2 Abundance |
|---|---|---|---|---|
| **Species** | | | | |
| Akkermansia muciniphila | 0.015000 | 0.018652 | 0.022194 | 0.019664 |
| Bacteroides fragilis | 0.140000 | 0.183928 | 0.213012 | 0.183693 |
| Bifidobacterium adolescentis | 0.060000 | 0.002497 | 0.002162 | 0.002509 |
| Candida albicans | 0.015000 | 0.020451 | 0.006930 | 0.004898 |
| Clostridioides difficile | 0.015000 | 0.020101 | 0.018865 | 0.019984 |
| Clostridium perfringens | 0.000001 | 0.000002 | 0.000000 | 0.000019 |
| Enterococcus faecalis | 0.000010 | 0.000006 | 0.000010 | 0.000019 |
| Escherichia coli | 0.140000 | 0.182881 | 0.228607 | 0.170464 |
| Faecalibacterium prausnitzii | 0.140000 | 0.211021 | 0.139893 | 0.196027 |
| Fusobacterium nucleatum | 0.060000 | 0.054483 | 0.043380 | 0.052388 |
| Lactobacillus fermentum | 0.060000 | 0.037100 | 0.007752 | 0.000000 |
| Methanobrevibacter smithii | 0.001000 | 0.000383 | 0.000383 | 0.000389 |
| Prevotella corporis | 0.060000 | 0.077604 | 0.092569 | 0.000000 |
| Roseburia hominis | 0.140000 | 0.021540 | 0.017006 | 0.018999 |
| Saccharomyces cerevisiae | 0.014000 | 0.006562 | 0.007754 | 0.005940 |
| Salmonella enterica | 0.000100 | 0.000188 | 0.000217 | 0.000711 |
| Veillonella rogosae | 0.140000 | 0.160218 | 0.198888 | 0.000000 |
| Other | | | | 0.324297 |
| Unmapped | | 0.002382 | 0.000379 | |
| **Genus** | | | | |
| Akkermansia | 0.015000 | 0.018652 | 0.022194 | 0.019666 |
| Bacteroides | 0.140000 | 0.183928 | 0.213012 | 0.191026 |
| Bifidobacterium | 0.060000 | 0.002497 | 0.002162 | 0.002568 |
| Candida | 0.015000 | 0.020451 | 0.006930 | 0.004953 |
| Clostridioides | 0.015000 | 0.020101 | 0.018865 | 0.020034 |
| Clostridium | 0.000001 | 0.000002 | 0.000000 | 0.000241 |
| Enterococcus | 0.000010 | 0.000006 | 0.000010 | 0.000107 |
| Escherichia | 0.140000 | 0.182881 | 0.228607 | 0.175942 |
| Faecalibacterium | 0.140000 | 0.211021 | 0.139893 | 0.212330 |
| Fusobacterium | 0.060000 | 0.054483 | 0.043380 | 0.053204 |
| Lactobacillus | 0.060000 | 0.037100 | 0.007752 | 0.000000 |
| Methanobrevibacter | 0.001000 | 0.000383 | 0.000383 | 0.000399 |
| Prevotella | 0.060000 | 0.077604 | 0.092569 | 0.047521 |
| Roseburia | 0.140000 | 0.021540 | 0.017006 | 0.019518 |
| Saccharomyces | 0.014000 | 0.006562 | 0.007754 | 0.005975 |
| Salmonella | 0.000100 | 0.000188 | 0.000217 | 0.000841 |
| Veillonella | 0.140000 | 0.160218 | 0.198888 | 0.160689 |
| Not validated | | | | 0.029441 |
| Other | | | | 0.055546 |
| Unmapped | | 0.002382 | 0.000379 | |
