## Supplementary Table 2 for "MetaGut: Insights into gut microbiomes in stem cell transplantation by comprehensive shotgun long-read sequencing"

| Pat ID | Start Cluster | Outcomes (non | Replase_1 | Day relative to | Replase_2 | Day relative to | 2nd HSCT | Day relative to | aGVHD Grade 1 | Day relative to | aGvHD Grade 3 | Day relative to | moderate cGVH | Day relative to | severe cGvHD | Day relative to | Death | Day relative to |
|---|---|---|---|---|---|---|---|---|---|---|---|---|---|---|---|---|---|---|
| 4 | 1 | 1 | 1 | 91 | | | | | | | | | | | | | 1 | 276 |
| 7 | 1 | 1 | 0 | | | | | | | | | | 1 | 190 | | | 0 | |
| 11 | 1 | 0 | 0 | | | | | | | | | | | | | | 0 | |
| 12 | 1 | 0 | 0 | | | | | | | | | | | | | | 0 | |
| 13 | 2 | 0 | 0 | | | | | | | | | | | | | | 0 | |
| 14 | 1 | 1 | 1 | 117 | | | | | | | | | 1 | 507 | | | 0 | |
| 15 | 1 | 1 | 1 | 198 | 1 | 579 | 1 | 481 | | | 1 | 510 | | | | | 1 | 612 |
| 16 | 1 | 1 | 1 | 58 | 1 | 534 | | | | | | | | | 1 | 443 | 0 | |
| 17 | 1 | 0 | 0 | | | | | | | | | | | | | | 0 | |
| 18 | 2 | 1 | 1 | 82 | | | | | | | | | | | | | 0 | |
| 19 | 1 | 1 | 0 | | | | | | 1 ? | | | | | | | | 0 | |
| 20 | 3 | 1 | 0 | | | | | | | | | | | | | | 1 | 32 |
| 21 | 1 | 1 | 1 | 108 | | | | | | | | | 1 | 165 | | | 1 | 221 |
| 22 | 2 | 0 | 0 | | | | | | | | | | | | | | 0 | |
| 23 | 2 | 0 | 0 | | | | | | | | | | | | | | 0 | |
| 24 | 1 | 0 | 0 | | | | | | | | | | | | | | 0 | |
| 25 | 2 | 1 | 0 | | | | | | 1 ? | | | | | | | | 0 | |
| 26 | 3 | 1 | 0 | 27 | | | | | 1 ? | | | | | | | | 0 | |
| 28 | 1 | 1 | 1 | 99 | | | | | | | | | | | | | 0 | |
| 29 | 1 | 0 | 0 | | | | | | | | | | | | | | 0 | |
| 31 | 3 | 0 | 0 | | | | | | | | | | | | | | 0 | |
| 32 | 1 | 1 | 0 | | | | | | | | | | | | | | 1 | 22 |
| 33 | 3 | 1 | 1 | 447 | | | | | | | | | | | | | 0 | |
| 34 | 1 | 1 | 1 | | | | | | | | | | | | | | 0 | |
| 36 | 3 | 1 | 1 | 255 | | | | | | | | | | | | | 0 | |
| 37 | 2 | 0 | 0 | | | | | | | | | | | | | | 0 | |
| 38 | 1 | 1 | 1 | 100 | | | | | | | | | | | | | 0 | |
| 39 | 1 | 0 | 0 | | | | | | | | | | | | | | 0 | |
| 40 | 2 | 0 | 0 | | | | | | | | | | | | | | 0 | |
| 41 | 2 | 1 | 1 | 100 | | | | | | | | | 1 | 229 | | | 0 | |
| 42 | 1 | 0 | 0 | | | | | | | | | | | | | | 0 | |
| 18.2 | 3 | 1 | 1 | 29 | | | 1 | 138 | | | | | | | | | 1 | 150 |
