## Supplementary Table 3 for "MetaGut: Insights into gut microbiomes in stem cell transplantation by comprehensive shotgun long-read sequencing"

| Patient ID | Day | Weight of the extracted stool (g) | Total amount of extract DNA (ng) | DNA µg / g Stool | Reads Total | Reads / ng DNA Library | Gbb Total | Gbp / ng DNA Library | Read Lengths Median | Read Lengths Mean | Read Lengths Standard Deviation | Read Lengths Min | Read Lengths Max | Total Reads | | Classified reads | | | Classified reads w/o human and plants | | | | | ARG Reads | Phyla | | Genera | | Species | |
|---|---|---|---|---|---|---|---|---|---|---|---|---|---|---|---|---|---|---|---|---|---|---|---|---|---|---|---|---|---|---|
| | | | | | | | | | | | | | | Classified (%) | Unclassified (%) | Chordata (%) | Plantae (%) | Microbial (%) | Bacteria (%) | Fungi (%) | Viruses (%) | Archae (%) | Other eukaryota (%) | per analysed Reads *10000 | not normalised | normalised w/o human and plants | not normalised | normalised w/o human and plants | not normalised | normalised w |
| 4 | -7 | 1 | 4128 | 4.13 | 1.38E+06 | 1.21E+04 | 4.51E+06 | 3.95E+04 | 1156 | 3278.28 | 4416.4 | 39 | 58102 | 81.23 | 18.77 | 1.31 | 4.74 | 75.17 | 98.49 | 0.37 | 0.04 | 0.05 | 1.06 | 18 | 70 | 9 | 2121 | 64 | 6946 | 120 |
| | 1 | 1 | 9.7 | 0.01 | 1.15E+05 | 1.98E+04 | 2.12E+05 | 3.64E+04 | 1173 | 1840.93 | 1995.48 | 5 | 52275 | 97.44 | 2.56 | 95.53 | 0.21 | 1.7 | 85.5 | 3.37 | 1.69 | 0.05 | 9.4 | 4 | 20 | 3 | 159 | 6 | 223 | 11 |
| 7 | -6 | 1 | 244.8 | 0.24 | 9.15E+05 | 1.21E+04 | 3.83E+06 | 5.05E+04 | 2592 | 4187.01 | 4899.22 | 2 | 106341 | 89.73 | 10.27 | 0.3 | 1.21 | 88.22 | 66.58 | 0.16 | 33.03 | 0.02 | 0.22 | 11 | 58 | 8 | 1705 | 58 | 4545 | 84 |
| 11 | -2 | 1 | 643.2 | 0.64 | 5.72E+05 | 3.96E+03 | 1.83E+06 | 1.27E+04 | 1766 | 3205.57 | 3761.19 | 30 | 77681 | 94.19 | 5.81 | 2.04 | 2.87 | 89.29 | 99.02 | 0.17 | 0.02 | 0.01 | 0.77 | 26 | 53 | 6 | 1159 | 41 | 2670 | 69 |
| | 0 | 1 | 367.72 | 0.12 | 3.62E+03 | 7.24E+01 | 1.60E+04 | 3.19E+02 | 2149 | 4409.51 | 5391.14 | 34 | 34204 | 92.54 | 7.46 | 14.11 | 8.98 | 69.46 | 93.76 | 3.34 | 0.04 | 0.04 | 2.82 | 59 | 13 | 7 | 79 | 33 | 125 | 68 |
| | 182 | 1 | 5568 | 5.57 | 2.08E+07 | 2.92E+04 | 2.27E+07 | 3.17E+04 | 483 | 1087.64 | 1818.3 | 30 | 55599 | 74.79 | 25.21 | 0.59 | 1.69 | 72.5 | 99.04 | 0.58 | 0.03 | 0.04 | 0.31 | 6 | 79 | 6 | 2566 | 71 | 9263 | 107 |
| 12 | -6 | 1 | 5760 | 5.76 | 3.83E+06 | 2.88E+03 | 1.27E+07 | 9.55E+03 | 976 | 3319.34 | 6041.7 | 20 | 137446 | 86.43 | 13.57 | 0.85 | 0.89 | 84.69 | 99.6 | 0.12 | 0.1 | 0.01 | 0.17 | 35 | 74 | 6 | 2154 | 47 | 7138 | 84 |
| | 34 | 1 | 676.8 | 0.23 | 1.96E+05 | 3.73E+03 | 1.25E+06 | 2.37E+04 | 4516.5 | 6356.61 | 6510.87 | 30 | 127783 | 98.83 | 1.17 | 2.88 | 0.51 | 95.45 | 97.61 | 1.71 | 0.62 | 0 | 0.06 | 112 | 19 | 5 | 182 | 10 | 407 | 20 |
| | 342 | 1 | 3753.6 | 3.75 | 5.09E+06 | 3.64E+04 | 4.34E+06 | 3.10E+04 | 407 | 853.29 | 1937.75 | 6 | 439541 | 68.3 | 31.7 | 9.19 | 0.93 | 58.18 | 99.17 | 0.27 | 0.15 | 0.04 | 0.36 | 3 | 74 | 7 | 2284 | 74 | 8019 | 123 |
| 13 | -7 | 1 | 1660.8 | 1.66 | 1.83E+06 | 1.73E+04 | 7.22E+06 | 6.82E+04 | 1033 | 3947.82 | 5523.15 | 4 | 82075 | 76.96 | 23.04 | 1.83 | 2 | 73.13 | 98.16 | 0.65 | 0.67 | 0.03 | 0.49 | 45 | 71 | 13 | 2006 | 69 | 6265 | 116 |
| | 298 | 1 | 1497.6 | 1.5 | 4.59E+06 | 3.28E+04 | 8.75E+06 | 6.25E+04 | 620 | 1906.36 | 3542.05 | 11 | 92459 | 52.82 | 47.18 | 1.57 | 2.34 | 48.91 | 98.11 | 0.75 | 0.13 | 0.11 | 0.89 | 13 | 80 | 12 | 2414 | 138 | 8814 | 190 |
| 14 | -7 | 1 | 864 | 0.86 | 1.04E+06 | 1.70E+04 | 2.76E+06 | 4.48E+04 | 1089 | 2641.83 | 3737.08 | 11 | 76480 | 72.46 | 27.54 | 0.75 | 1.78 | 69.93 | 98.81 | 0.45 | 0.16 | 0.09 | 0.48 | 12 | 73 | 10 | 2191 | 109 | 7488 | 179 |
| | 558 | 1 | 8208 | 0.04 | 1.31E+07 | 5.33E+04 | 2.19E+07 | 8.95E+04 | 634 | 1678.6 | 2877.55 | 2 | 702383 | 67.05 | 32.95 | 0.61 | 2.26 | 64.17 | 95.36 | 0.44 | 3.45 | 0.09 | 0.66 | 7 | 82 | 11 | 2627 | 114 | 9491 | 161 |
| 15 | -6 | 1 | 14016 | 14.02 | 1.44E+07 | 2.06E+04 | 1.01E+08 | 1.44E+05 | 4387 | 6958.21 | 6760.92 | 15 | 109429 | 95.44 | 4.56 | 0.52 | 1.44 | 93.48 | 99.53 | 0.15 | 0.04 | 0.02 | 0.26 | 31 | 77 | 6 | 2406 | 63 | 8755 | 99 |
| | 6 | 4 | 134.4 | 0.03 | 2.59E+06 | 3.82E+04 | 6.17E+06 | 9.10E+04 | 899 | 2384.31 | 3695.59 | 3 | 71237 | 95.3 | 4.7 | 3.45 | 0.52 | 91.33 | 97.89 | 0.27 | 1.77 | 0 | 0.07 | 30 | 54 | 5 | 876 | 10 | 1911 | 14 |
| | 16 | 4 | 268.8 | 0.07 | 5.18E+05 | 3.19E+03 | 1.22E+06 | 7.54E+03 | 1027 | 2361.78 | 3928.6 | 4 | 101699 | 95.94 | 4.06 | 1.72 | 1.3 | 92.92 | 99.25 | 0.49 | 0.17 | 0 | 0.08 | 29 | 30 | 4 | 528 | 10 | 1048 | 21 |
| | 104 | 1 | 1593.6 | 1.59 | 4.22E+06 | 1.81E+04 | 1.55E+07 | 6.64E+04 | 1110 | 3665.69 | 5005.53 | 21 | 67259 | 68.55 | 31.45 | 1.74 | 3.84 | 62.97 | 95.2 | 3.51 | 0.29 | 0.09 | 0.92 | 43 | 77 | 10 | 2362 | 111 | 8664 | 153 |
| | 189 | 1 | 4992 | 4.99 | 1.60E+06 | 3.09E+03 | 1.98E+06 | 3.82E+03 | 607 | 1237.42 | 2036.53 | 2 | 76715 | 80.81 | 19.19 | 0.62 | 0.79 | 79.39 | 99.69 | 0.11 | 0.03 | 0.01 | 0.15 | 10 | 66 | 8 | 1845 | 39 | 5220 | 71 |
| 16 | -5 | 1 | 2390.4 | 2.39 | 5.75E+06 | 1.15E+04 | 1.00E+07 | 2.01E+04 | 599 | 1738.76 | 3162.22 | 8 | 63486 | 83.69 | 16.31 | 0.37 | 0.97 | 82.35 | 99.66 | 0.12 | 0.01 | 0.02 | 0.19 | 11 | 75 | 6 | 2280 | 39 | 8009 | 74 |
| | 2 | 1 | 835.2 | 0.84 | 1.37E+06 | 7.77E+03 | 3.70E+06 | 2.10E+04 | 959 | 2701.86 | 3916.66 | 16 | 73364 | 90.23 | 9.77 | 0.5 | 0.69 | 89.05 | 99.6 | 0.24 | 0.02 | 0.01 | 0.12 | 40 | 57 | 5 | 1638 | 26 | 4649 | 47 |
| | 163 | 1 | 5280 | 5.28 | 2.80E+06 | 1.20E+04 | 2.25E+07 | 9.65E+04 | 4363 | 8038.2 | 9355.38 | 14 | 116663 | 96.3 | 3.7 | 0.86 | 0.51 | 94.93 | 47.21 | 0.09 | 52.59 | 0 | 0.1 | 47 | 57 | 4 | 1460 | 26 | 3686 | 38 |
| | 171 | 4 | 3792 | 0.95 | 2.48E+06 | 1.77E+04 | 8.98E+06 | 6.41E+04 | 1516 | 3624.2 | 4587.52 | 5 | 83128 | 97.94 | 2.06 | 1.58 | 0.46 | 95.9 | 39.68 | 1.53 | 58.71 | 0 | 0.08 | 37 | 52 | 6 | 1076 | 25 | 2542 | 36 |
| 17 | -7 | 1 | 3676.8 | 3.68 | 1.44E+06 | 3.71E+03 | 4.22E+06 | 1.08E+04 | 1042 | 2923.77 | 4930.82 | 4 | 102599 | 93.6 | 6.4 | 0.21 | 0.62 | 92.76 | 88.84 | 0.11 | 10.86 | 0.1 | 0.09 | 33 | 57 | 7 | 1334 | 23 | 3257 | 69 |
| | 9 | 1 | 2476.8 | 2.48 | 3.07E+06 | 1.32E+04 | 1.85E+07 | 7.94E+04 | 3415 | 6026.4 | 6948.79 | 83 | 124123 | 96.4 | 3.6 | 0.82 | 1.08 | 94.51 | 94.46 | 0.32 | 4.97 | 0.01 | 0.25 | 121 | 67 | 6 | 1615 | 19 | 4299 | 41 |
| | 13 | 1 | 2611.2 | 2.61 | 2.46E+06 | 1.05E+04 | 1.39E+07 | 5.98E+04 | 2754 | 5668.94 | 7113.58 | 95 | 109921 | 96.93 | 3.07 | 0.2 | 0.54 | 96.2 | 95.76 | 0.13 | 4 | 0.01 | 0.11 | 99 | 65 | 8 | 1656 | 22 | 4693 | 47 |
| | 154 | 1 | 5568 | 5.57 | 8.25E+04 | 7.05E+02 | 1.90E+05 | 1.62E+03 | 525 | 2303.72 | 3827.73 | 54 | 55314 | 65.14 | 34.86 | 0.64 | 1.91 | 62.59 | 99.09 | 0.32 | 0.03 | 0.03 | 0.54 | 6 | 45 | 8 | 648 | 55 | 1217 | 102 |
| 18 | -3 | 5 | 326.21 | 0.07 | 2.05E+05 | 2.89E+03 | 6.74E+05 | 9.48E+03 | 1058 | 3283.44 | 5233.92 | 23 | 70716 | 98.91 | 1.09 | 0.25 | 0.22 | 98.44 | 99.17 | 0.71 | 0.07 | 0 | 0.05 | 146 | 21 | 3 | 277 | 21 | 717 | 51 |
| | 13 | 1 | 1152 | 1.15 | 6.51E+06 | 2.79E+04 | 1.74E+07 | 7.47E+04 | 1179 | 2675.82 | 4126.01 | 87 | 101250 | 95.75 | 4.25 | 4.27 | 1.27 | 90.21 | 95.65 | 0.41 | 3.74 | 0.01 | 0.19 | 52 | 72 | 8 | 1817 | 20 | 5257 | 49 |
| | 91 | 3 | 437.76 | 0.15 | 9.42E+04 | 4.04E+02 | 8.67E+04 | 3.72E+02 | 390 | 920.4 | 2085.81 | 56 | 46420 | 59.44 | 40.56 | 7.07 | 1.92 | 50.45 | 98.62 | 0.67 | 0.03 | 0.05 | 0.62 | 4 | 45 | 7 | 793 | 77 | 1552 | 103 |
| 18.2 | -9 | 4 | 84.96 | 0.02 | 2.33E+05 | 5.10E+03 | 7.42E+05 | 1.63E+04 | 1072 | 3188.13 | 4635.88 | 16 | 154747 | 95.86 | 4.14 | 22.55 | 0.89 | 72.41 | 96.8 | 2.35 | 0.46 | 0 | 0.38 | 79 | 28 | 7 | 354 | 13 | 686 | 15 |
| | 16 | 4 | 112.32 | 0.03 | 1.81E+04 | 3.63E+02 | 3.11E+04 | 6.24E+02 | 525 | 1718.35 | 3376.84 | 42 | 108936 | 89.21 | 10.79 | 22.46 | 1.77 | 64.98 | 95.44 | 2.63 | 0.82 | 0.01 | 1.1 | 19 | 18 | 9 | 174 | 44 | 334 | 65 |
| | 71 | 4 | 0.48 | 0 | 1.98E+03 | 1.33E+03 | 6.47E+03 | 4.33E+03 | 996 | 3267.11 | 4917.61 | 3 | 38320 | 75.11 | 24.89 | 1.87 | 1.26 | 71.98 | 94.53 | 4.84 | 0.07 | 0 | 0.56 | 46 | 9 | 6 | 69 | 52 | 110 | 67 |
| 19 | -2 | 1 | 781.44 | 0.39 | 4.74E+05 | 1.36E+03 | 9.98E+05 | 2.87E+03 | 654 | 2105.74 | 3935.75 | 9 | 72156 | 89.57 | 10.43 | 1.28 | 0.51 | 87.78 | 99.74 | 0.12 | 0.03 | 0.01 | 0.09 | 13 | 47 | 6 | 979 | 35 | 2047 | 73 |
| | 153 | 1 | 2448 | 2.45 | 8.26E+06 | 5.90E+04 | 1.31E+07 | 9.33E+04 | 590 | 1582.28 | 3070.97 | 1 | 298458 | 69.55 | 30.45 | 0.42 | 1.94 | 67.19 | 99.07 | 0.33 | 0.02 | 0.06 | 0.52 | 6 | 81 | 9 | 2483 | 85 | 8921 | 123 |
| 20 | -4 | 4 | 564 | 0.14 | 1.31E+06 | 5.61E+03 | 2.19E+06 | 9.39E+03 | 1104 | 1672.61 | 2070.17 | 32 | 55121 | 98.25 | 1.75 | 87.95 | 0.21 | 10.08 | 95.94 | 2.24 | 0.45 | 0.01 | 1.36 | 23 | 34 | 3 | 481 | 7 | 918 | 9 |
| 21 | -9 | 4 | 600.1 | 0.15 | 2.36E+06 | 1.01E+04 | 3.71E+06 | 1.59E+04 | 514 | 1570.43 | 2870.02 | 23 | 107709 | 75.59 | 24.41 | 4.45 | 1.4 | 69.74 | 96.85 | 1.63 | 1.05 | 0.03 | 0.44 | 7 | 72 | 10 | 2192 | 65 | 7315 | 120 |
| | 50 | 1 | 2995.2 | 3 | 2.30E+06 | 9.85E+03 | 6.54E+06 | 2.80E+04 | 576 | 2846.31 | 5408.81 | 32 | 84416 | 72.22 | 27.78 | 2.31 | 2.72 | 67.19 | 98.53 | 0.57 | 0.07 | 0.06 | 0.78 | 17 | 77 | 11 | 2277 | 89 | 8044 | 113 |
| | 120 | 1 | 11904 | 11.9 | 8.29E+04 | 7.08E+02 | 1.81E+05 | 1.55E+03 | 633 | 2181 | 3624.22 | 71 | 46887 | 66.55 | 33.45 | 0.51 | 1.73 | 64.31 | 96.15 | 0.32 | 3 | 0.04 | 0.5 | 4 | 39 | 11 | 766 | 70 | 1449 | 99 |
| 22 | -4 | 4 | 777.6 | 0.19 | 6.50E+06 | 2.79E+04 | 8.73E+06 | 3.75E+04 | 512 | 1344 | 2487.09 | 28 | 66056 | 76.18 | 23.82 | 1.82 | 1.01 | 73.34 | 99.33 | 0.33 | 0.07 | 0.02 | 0.25 | 5 | 77 | 6 | 2312 | 55 | 8062 | 78 |
| | 0 | 4 | 63.13 | 0.02 | 6.48E+04 | 1.24E+03 | 1.70E+05 | 3.25E+03 | 815 | 2625.65 | 4229.5 | 15 | 58135 | 95.14 | 4.86 | 9.81 | 1.42 | 83.91 | 96.59 | 1.32 | 1.83 | 0.01 | 0.25 | 8 | 23 | 6 | 223 | 20 | 473 | 43 |
| | 91 | 1 | 2956.8 | 2.96 | 1.15E+07 | 8.20E+04 | 1.53E+07 | 1.09E+05 | 406 | 1332.91 | 3151.75 | 9 | 340155 | 61.75 | 38.25 | 2.05 | 2.28 | 57.42 | 98.72 | 0.43 | 0.02 | 0.07 | 0.75 | 4 | 80 | 11 | 2545 | 90 | 9211 | 116 |
| 23 | -9 | 1 | 4435.2 | 4.44 | 2.03E+06 | 8.70E+03 | 5.91E+06 | 2.54E+04 | 731 | 2914.64 | 5070.67 | 7 | 68977 | 45.35 | 54.65 | 0.65 | 3.07 | 41.63 | 97.39 | 1.11 | 0.05 | 0.2 | 1.25 | 16 | 76 | 15 | 2333 | 185 | 8397 | 247 |
| | 12 | 2 | 130.85 | 0.07 | 1.86E+05 | 3.71E+03 | 4.98E+05 | 9.94E+03 | 465 | 2681.75 | 5313.49 | 7 | 74243 | 35.63 | 64.37 | 0.88 | 2.11 | 32.64 | 97.31 | 1.24 | 0.03 | 0.2 | 1.22 | 14 | 58 | 14 | 1591 | 194 | 4529 | 272 |
| | 61 | 1 | 3264 | 3.26 | 2.66E+06 | 1.90E+04 | 1.14E+07 | 8.13E+04 | 974 | 4270.11 | 6889.38 | 11 | 455583 | 91.27 | 8.73 | 0.74 | 0.48 | 90.05 | 99.73 | 0.15 | 0.01 | 0.01 | 0.1 | 26 | 63 | 5 | 1849 | 32 | 5339 | 60 |
| | 130 | 1 | 4339.2 | 4.34 | 7.10E+05 | 9.87E+03 | 9.62E+05 | 1.34E+04 | 434 | 1354.59 | 3382.28 | 2 | 84898 | 56.37 | 43.63 | 1.6 | 2.45 | 52.31 | 98.46 | 0.6 | 0.02 | 0.08 | 0.84 | 7 | 67 | 11 | 1960 | 83 | 5978 | 120 |
| 24 | -10 | 1 | 3974.4 | 3.97 | 1.29E+06 | 5.52E+03 | 5.00E+06 | 2.14E+04 | 1124 | 3887.5 | 6059.25 | 4 | 85249 | 70.49 | 29.51 | 0.63 | 3.12 | 66.74 | 98.03 | 0.47 | 0.32 | 0.28 | 0.9 | 20 | 72 | 10 | 2164 | 88 | 7201 | 149 |
| | 0 | 4 | 635.9 | 0.16 | 1.23E+06 | 5.27E+03 | 3.38E+06 | 1.45E+04 | 673 | 2758.01 | 4876.98 | 31 | 83885 | 84.04 | 15.96 | 2.61 | 1.99 | 79.45 | 98.88 | 0.51 | 0.18 | 0.02 | 0.41 | 8 | 62 | 8 | 1385 | 43 | 3690 | 68 |
| | 9 | 4 | 751.1 | 0.19 | 1.62E+05 | 6.95E+02 | 1.79E+05 | 7.68E+02 | 357 | 1105 | 2681.97 | 29 | 55984 | 73.43 | 26.57 | 13.79 | 1.73 | 57.91 | 99.22 | 0.23 | 0.01 | 0.03 | 0.5 | 9 | 41 | 8 | 635 | 46 | 1369 | 56 |
| | 49 | 1 | 5568 | 5.57 | 4.02E+04 | 3.44E+02 | 2.24E+05 | 1.92E+03 | 1995 | 5573.7 | 6841.58 | 45 | 48589 | 47.64 | 52.36 | 1.51 | 7.55 | 38.58 | 92.74 | 2.35 | 0.12 | 0.39 | 4.41 | 27 | 51 | 21 | 1318 | 231 | 3218 | 294 |
| | 127 | 4 | 147.84 | 0.04 | 9.72E+03 | 9.88E+01 | 1.42E+04 | 1.45E+02 | 526 | 1464.64 | 2638.06 | 50 | 45938 | 43.41 | 56.59 | 9.61 | 3.25 | 30.55 | 92.76 | 3.54 | 1.35 | 0.1 | 2.26 | 5 | 31 | 19 | 350 | 119 | 505 | 162 |
| 25 | -62 | 1 | 1516.8 | 1.52 | 9.54E+06 | 4.09E+04 | 1.77E+07 | 7.60E+04 | 511 | 1856.38 | 3653.57 | 13 | 85973 | 53.92 | 46.08 | 1.39 | 2.21 | 50.32 | 98.37 | 0.72 | 0.08 | 0.07 | 0.77 | 8 | 82 | 10 | 2508 | 87 | 9018 | 120 |
| | 1 | 4 | 1372.8 | 0.34 | 2.87E+04 | 2.45E+02 | 1.20E+05 | 1.02E+03 | 1223 | 4167.48 | 5674.24 | 86 | 60133 | 90.77 | 9.23 | 6.67 | 0.48 | 83.63 | 99.75 | 0.1 | 0.03 | 0 | 0.11 | 306 | 15 | 5 | 144 | 26 | 298 | 39 |
| | 15 | 4 | 39.46 | 0.01 | 5.35E+04 | 4.67E+03 | 5.77E+04 | 5.04E+03 | 317 | 1078.57 | 2817.76 | 7 | 62036 | 91.42 | 8.58 | 9.39 | 3.78 | 78.25 | 96.47 | 3.03 | 0 | 0 | 0.49 | 22 | 21 | 5 | 170 | 10 | 278 | 20 |
| | 31 | 1 | 792 | 0.79 | 6.52E+04 | 5.57E+02 | 1.14E+05 | 9.74E+02 | 482 | 1747.33 | 3678.59 | 29 | 70521 | 86.74 | 13.26 | 1.7 | 0.46 | 84.59 | 98.96 | 0.94 | 0.01 | 0 | 0.08 | 92 | 20 | 4 | 218 | 24 | 445 | 51 |
| 26 | -3 | 4 | 33.5 | 0.01 | 3.16E+04 | 2.54E+03 | 8.57E+04 | 6.90E+03 | 432 | 2715.75 | 5215.24 | 27 | 53681 | 82.63 | 17.37 | 4.68 | 1.01 | 76.94 | 98.78 | 0.67 | 0.46 | 0 | 0.09 | 52 | 19 | 6 | 196 | 32 | 405 | 56 |
| | 12 | 1 | 1075.2 | 1.08 | 2.85E+06 | 1.22E+04 | 9.21E+06 | 3.95E+04 | 892 | 3227.06 | 5502.52 | 29 | 104940 | 96.15 | 3.85 | 4.73 | 0.19 | 91.23 | 99.91 | 0.05 | 0.01 | 0 | 0.03 | 71 | 53 | 3 | 1071 | 12 | 2563 | 24 |
| | 27 | 4 | 617.47 | 0.15 | 1.41E+04 | 7.68E+02 | 2.07E+04 | 1.13E+03 | 381 | 1469.55 | 3499.42 | 62 | 48500 | 87.57 | 12.43 | 11.38 | 1.32 | 74.87 | 96.88 | 2.24 | 0.01 | 0 | 0.87 | 16 | 15 | 7 | 123 | 34 | 216 | 48 |
| | 185 | 1 | 1046.4 | 1.05 | 1.86E+06 | 1.33E+04 | 9.70E+06 | 6.93E+04 | 3497 | 5214.7 | 5530.43 | 18 | 538317 | 81.18 | 18.82 | 1.06 | 1.89 | 78.22 | 97.92 | 0.53 | 0.89 | 0.12 | 0.53 | 35 | 74 | 13 | 2308 | 138 | 8299 | 192 |
| 28 | -5 | 1 | 3398.4 | 3.4 | 2.55E+06 | 1.10E+04 | 8.10E+06 | 3.48E+04 | 737 | 3171.25 | 5358.36 | 28 | 88078 | 81.65 | 18.35 | 0.44 | 0.9 | 80.31 | 99.54 | 0.24 | 0.01 | 0.01 | 0.2 | 18 | 71 | 8 | 1933 | 47 | 5744 | 81 |
| | 1 | 4 | 889.34 | 0.22 | 4.06E+06 | 1.74E+04 | 6.94E+06 | 2.98E+04 | 602 | 1709.44 | 3344.92 | 37 | 75900 | 92.46 | 7.54 | 7.12 | 4.19 | 81.14 | 99.56 | 0.24 | 0.01 | 0.01 | 0.18 | 11 | 71 | 6 | 1597 | 29 | 4356 | 41 |
| | 21 | 4 | 1593.6 | 0.4 | 2.08E+06 | 1.48E+04 | 1.02E+07 | 7.31E+04 | 2218 | 4926.3 | 5861.58 | 5 | 85867 | 98.24 | 1.76 | 0.18 | 0.37 | 97.69 | 99.54 | 0.09 | 0.3 | 0 | 0.07 | 82 | 47 | 5 | 977 | 20 | 2410 | 42 |
| | 63 | 1 | 4608 | 4.61 | 8.04E+04 | 6.87E+02 | 1.84E+05 | 1.58E+03 | 584 | 2290.95 | 3858.13 | 50 | 40479 | 74.62 | 25.38 | 0.74 | 1.32 | 72.56 | 99.03 | 0.35 | 0.25 | 0.02 | 0.36 | 15 | 33 | 7 | 483 | 46 | 894 | 76 |
| | 237 | 1 | 8208 | 8.21 | 5.05E+05 | 4.29E+03 | 6.25E+05 | 5.32E+03 | 447 | 1239.08 | 2846.06 | 15 | 84015 | 77.15 | 22.85 | 0.25 | 0.97 | 75.93 | 99.54 | 0.23 | 0.01 | 0.01 | 0.2 | 6 | 58 | 8 | 1191 | 48 | 2763 | 86 |
| 29 | -5 | 1 | 3945.6 | 3.95 | 3.54E+06 | 2.53E+04 | 1.14E+07 | 8.16E+04 | 1111 | 3228.04 | 4479.67 | 1 | 76057 | 83.81 | 16.19 | 0.68 | 1.75 | 81.38 | 98.52 | 0.92 | 0.15 | 0.02 | 0.39 | 21 | 70 | 9 | 2101 | 52 | 6582 | 94 |
| | 1 | 4 | 695.81 | 0.17 | 2.39E+06 | 1.71E+04 | 6.35E+06 | 4.54E+04 | 1092 | 2657.74 | 3803.61 | 19 | 67557 | 94.99 | 5.01 | 1.09 | 0.77 | 93.13 | 96.81 | 2.8 | 0.23 | 0.01 | 0.15 | 103 | 59 | 8 | 1401 | 31 | 3946 | 69 |
| | 15 | 4 | 8.83 | 0 | 1.28E+05 | 3.00E+04 | 7.98E+04 | 1.87E+04 | 345 | 622.11 | 1017.86 | 3 | 29180 | 89.38 | 10.62 | 31.49 | 7.69 | 50.19 | 84.58 | 13.17 | 0.11 | 0.02 | 2.11 | 0 | 38 | 8 | 394 | 26 | 700 | 30 |
| | 186 | 4 | 2083.2 | 0.52 | 9.22E+06 | 7.86E+04 | 1.14E+07 | 9.74E+04 | 626 | 1240.18 | 1927.89 | 1 | 570652 | 87.92 | 12.08 | 0.36 | 1.15 | 86.41 | 99.58 | 0.13 | 0.01 | 0.01 | 0.27 | 7 | 73 | 7 | 2143 | 42 | 7004 | 77 |
| 31 | -6 | 4 | 182.4 | 0.05 | 6.43E+04 | 1.13E+03 | 1.13E+05 | 1.99E+03 | 450 | 1759.97 | 3458.39 | 16 | 52676 | 92.34 | 7.66 | 35.76 | 0.65 | 55.93 | 98.42 | 0.7 | 0.19 | 0.02 | 0.67 | 33 | 20 | 6 | 214 | 23 | 404 | 31 |
| | 6 | 4 | 6.34 | 0 | 9.50E+04 | 2.92E+04 | 7.10E+04 | 2.19E+04 | 447 | 747.48 | 978.54 | 46 | 27764 | 93.67 | 6.33 | 91.04 | 0.47 | 2.16 | 71.27 | 11.42 | 0.39 | 0.19 | 16.72 | 0 | 27 | 5 | 215 | 14 | 313 | 13 |
| | 58 | 1 | 619.2 | 0.62 | 4.34E+05 | 1.26E+03 | 2.17E+06 | 6.30E+03 | 2590 | 5005.08 | 5734.21 | 30 | 107960 | 93.07 | 6.93 | 0.4 | 1.86 | 90.81 | 94.28 | 5.28 | 0.01 | 0.01 | 0.42 | 127 | 48 | 7 | 1130 | 47 | 2834 | 72 |
| | 178 | 1 | 5.09 | 0.01 | 4.45E+03 | 4.45E+03 | 4.27E+03 | 4.27E+03 | 499 | 959 | 1934.62 | 28 | 36089 | 73.79 | 26.21 | 3.71 | 2.81 | 67.28 | 98.93 | 0.47 | 0.07 | 0 | 0.53 | 5 | 14 | 8 | 136 | 76 | 220 | 107 |
| 32 | -5 | 4 | 108.48 | 0.03 | 4.28E+04 | 1.72E+03 | 3.75E+04 | 1.51E+03 | 262 | 877.31 | 2164.93 | 11 | 54277 | 73.16 | 26.84 | 10.56 | 1.03 | 61.56 | 98.12 | 1.02 | 0.51 | 0.01 | 0.35 | 6 | 26 | 7 | 314 | 45 | 566 | 70 |
| 33 | -2 | 4 | 44.06 | 0.01 | 4.81E+04 | 2.40E+03 | 1.06E+05 | 5.29E+03 | 571 | 2203.09 | 4223.14 | 6 | 61312 | 92.67 | 7.33 | 46.7 | 0.8 | 45.16 | 97.87 | 0.8 | 0.63 | 0.01 | 0.68 | 54 | 21 | 8 | 221 | 29 | 421 | 46 |
| | 7 | 4 | 93.12 | 0.02 | 1.09E+05 | 4.37E+03 | 1.78E+05 | 7.14E+03 | 408 | 1633.9 | 3807.58 | 10 | 132453 | 88.66 | 11.34 | 14.73 | 2.4 | 71.52 | 94.92 | 4.51 | 0.04 | 0.01 | 0.53 | 22 | 31 | 6 | 561 | 29 | 1137 | 41 |
| | 47 | 1 | 1286.4 | 1.29 | 2.51E+05 | 6.48E+03 | 3.91E+05 | 1.01E+04 | 647 | 1558.17 | 2893.74 | 45 | 227144 | 74.81 | 25.19 | 0.39 | 2.14 | 72.29 | 99.05 | 0.28 | 0.1 | 0.04 | 0.53 | 8 | 51 | 7 | 1242 | 58 | 2930 | 94 |
| 34 | -6 | 1 | 7728 | 7.73 | 6.39E+06 | 4.57E+04 | 1.06E+07 | 7.58E+04 | 554 | 1659.25 | 3622.98 | 5 | 830851 | 83.16 | 16.84 | 0.62 | 1.87 | 80.68 | 99.27 | 0.25 | 0.01 | 0.02 | 0.45 | 20 | 71 | 6 | 2084 | 31 | 6495 | 61 |
| | 10 | 1 | 4.99 | 0 | 3.41E+04 | 1.11E+04 | 4.22E+04 | 1.37E+04 | 423 | 1239.01 | 2447.76 | 43 | 42388 | 87.68 | 12.32 | 22.62 | 3.53 | 61.52 | 85.8 | 10.81 | 2.3 | 0.01 | 1.07 | 21 | 24 | 7 | 266 | 37 | 543 | 80 |
| | 18 | 4 | 48 | 0.01 | 4.15E+05 | 2.05E+04 | 2.74E+05 | 1.36E+04 | 369 | 660.97 | 1172.33 | 2 | 57123 | 91.3 | 8.7 | 37.85 | 1.72 | 51.74 | 82.51 | 15.71 | 0.31 | 0.01 | 1.45 | 12 | 37 | 8 | 734 | 19 | 1625 | 49 |
| | 41 | 1 | 1756.8 | 1.76 | 3.49E+05 | 9.21E+03 | 7.63E+05 | 2.02E+04 | 963 | 2188.9 | 3488.08 | 41 | 67835 | 80.8 | 19.2 | 6.84 | 2.66 | 71.3 | 98.83 | 0.44 | 0.01 | 0.04 | 0.68 | 3 | 53 | 8 | 1302 | 45 | 3122 | 76 |
| 36 | -6 | 4 | 77.76 | 0.02 | 2.86E+04 | 3.77E+02 | 1.05E+05 | 1.39E+03 | 1411 | 3685.91 | 5048.76 | 6 | 50610 | 93.65 | 6.35 | 14.78 | 1.13 | 77.74 | 98.02 | 0.75 | 0.63 | 0.01 | 0.59 | 46 | 18 | 7 | 181 | 31 | 372 | 52 |
| | 38 | 4 | 734.4 | 0.18 | 1.12E+04 | 4.80E+01 | 2.42E+04 | 1.04E+02 | 768 | 2166.66 | 3454.45 | 27 | 39034 | 85.63 | 14.37 | 0.89 | 1.2 | 83.54 | 99.15 | 0.54 | 0.03 | 0.03 | 0.25 | 51 | 20 | 7 | 166 | 48 | 267 | 64 |
| | 63 | 1 | 1132.8 | 1.13 | 8.91E+06 | 6.36E+04 | 1.82E+07 | 1.30E+05 | 788 | 2046.44 | 2866.62 | 17 | 171700 | 66.19 | 33.81 | 0.48 | 2.01 | 63.7 | 98.82 | 0.46 | 0.05 | 0.07 | 0.6 | 9 | 82 | 11 | 2511 | 95 | 9088 | 144 |
| 37 | -5 | 4 | 28.13 | 0.01 | 7.40E+03 | 5.80E+02 | 1.07E+04 | 8.35E+02 | 536.5 | 1439.91 | 2746.43 | 1 | 40594 | 91.06 | 8.94 | 18.65 | 1.34 | 71.07 | 96.66 | 1.9 | 0.89 | 0.02 | 0.53 | 56 | 19 | 7 | 131 | 39 | 253 | 71 |
| | 1 | 4 | 46.08 | 0.01 | 2.00E+04 | 2.95E+03 | 1.59E+04 | 2.33E+03 | 360 | 791.24 | 1665.11 | 35 | 42532 | 92.8 | 7.2 | 2.22 | 0.53 | 90.04 | 99.41 | 0.46 | 0.04 | 0 | 0.09 | 80 | 15 | 5 | 146 | 35 | 309 | 60 |
| | 17 | 4 | 33.5 | 0.01 | 1.19E+03 | 1.12E+02 | 3.15E+03 | 2.97E+02 | 817 | 2644.7 | 4131.64 | 22 | 27146 | 86.38 | 13.62 | 9 | 0.76 | 76.62 | 97.8 | 1.87 | 0 | 0 | 0.33 | 27 | 7 | 7 | 45 | 33 | 68 | 55 |
| | 22 | 6 | 18.14 | 0 | 3.86E+03 | 1.00E+03 | 9.28E+03 | 2.40E+03 | 887 | 2403.4 | 3856.57 | 17 | 40955 | 89.97 | 10.03 | 50.39 | 0.98 | 38.6 | 94.9 | 4.03 | 0.07 | 0 | 1.01 | 84 | 12 | 8 | 76 | 36 | 147 | 57 |
| | 138 | 1 | 1382.4 | 1.38 | 8.82E+05 | 7.39E+03 | 8.39E+05 | 7.03E+03 | 525 | 951.57 | 1773.21 | 29 | 55638 | 70.31 | 29.69 | 3.31 | 2.76 | 64.24 | 98.68 | 0.56 | 0.02 | 0.04 | 0.71 | 3 | 70 | 9 | 1694 | 58 | 4568 | 80 |
| 38 | -6 | 1 | 4550.4 | 4.55 | 5.37E+06 | 3.83E+04 | 1.74E+07 | 1.24E+05 | 1239 | 3238.34 | 4807.68 | 6 | 291076 | 85.94 | 14.06 | 0.5 | 1.66 | 83.79 | 99.37 | 0.22 | 0.01 | 0.03 | 0.37 | 7 | 77 | 7 | 2335 | 65 | 8426 | 92 |
| | 15 | 4 | 64.8 | 0.02 | 9.13E+03 | 8.36E+02 | 2.25E+04 | 2.06E+03 | 651.5 | 2462.19 | 4243.34 | 42 | 41847 | 72.81 | 27.19 | 2.85 | 1.8 | 68.16 | 89.18 | 4.07 | 6.36 | 0.03 | 0.35 | 36 | 14 | 6 | 195 | 58 | 358 | 88 |
| 39 | -6 | 1 | 998.4 | 1 | 2.51E+06 | 1.79E+04 | 3.10E+06 | 2.21E+04 | 789 | 1234.17 | 1431.77 | 10 | 157801 | 61.46 | 38.54 | 0.76 | 4.04 | 56.66 | 97.12 | 0.71 | 0.6 | 0.15 | 1.42 | 4 | 76 | 18 | 2398 | 134 | 8596 | 213 |
| | 1 | 6 | 13.54 | 0 | 5.23E+03 | 4.65E+02 | 2.11E+04 | 1.88E+03 | 1668 | 4035.76 | 5182.7 | 12 | 36290 | 89.45 | 10.55 | 3.21 | 1.84 | 84.4 | 95.4 | 3.85 | 0.25 | 0 | 0.5 | 96 | 14 | 7 | 136 | 61 | 272 | 92 |
| | 15 | 1 | 892.8 | 0.89 | 4.56E+06 | 3.25E+04 | 2.22E+07 | 1.59E+05 | 3179 | 4878.22 | 5266.06 | 19 | 117617 | 99.44 | 0.56 | 98.09 | 0.1 | 1.25 | 82.96 | 0.92 | 0.11 | 0.07 | 15.93 | 2 | 47 | 3 | 939 | 5 | 1956 | 3 |
| | 115 | 1 | 12960 | 12.96 | 3.50E+05 | 2.73E+03 | 5.89E+05 | 4.60E+03 | 476 | 1680.73 | 4261.37 | 10 | 180296 | 89.19 | 10.81 | 0.37 | 0.36 | 88.47 | 99.85 | 0.07 | 0 | 0.01 | 0.07 | 5 | 42 | 6 | 845 | 33 | 1597 | 63 |
| 40 | -1 | 4 | 609.6 | 0.15 | 2.25E+03 | 2.00E+01 | 7.48E+03 | 6.70E+01 | 1054.5 | 3330.65 | 4887.17 | 28 | 38041 | 82.37 | 17.63 | 1.02 | 1.16 | 80.19 | 96.45 | 3.28 | 0.11 | 0.06 | 0.11 | 63 | 13 | 9 | 98 | 59 | 172 | 103 |
| | 17 | 1 | 1084.8 | 1.08 | 5.85E+07 | 4.87E+05 | 7.09E+07 | 5.89E+05 | 607 | 1211.22 | 1842.67 | 1 | 549526 | 90.9 | 9.1 | 0.11 | 0.57 | 90.22 | 99.79 | 0.08 | 0.01 | 0.01 | 0.11 | 12 | 82 | 5 | 2417 | 27 | 8524 | 50 |
| 41 | -6 | 4 | 13.25 | 0 | 3.89E+03 | 3.89E+02 | 1.35E+04 | 1.35E+03 | 1113 | 3463.88 | 5013.11 | 30 | 57555 | 85.41 | 14.59 | 6.63 | 2.03 | 76.76 | 91.07 | 4.08 | 4.18 | 0.07 | 0.6 | 82 | 12 | 8 | 113 | 65 | 248 | 113 |
| | 14 | 4 | 41.09 | 0.01 | 1.05E+04 | 5.58E+02 | 1.16E+04 | 6.21E+02 | 509 | 1111.45 | 2141.69 | 33 | 38812 | 76.15 | 23.85 | 2.5 | 2.1 | 71.54 | 98.21 | 0.53 | 0.7 | 0.01 | 0.55 | 8 | 20 | 9 | 190 | 65 | 324 | 110 |
| 42 | -8 | 4 | 1862.4 | 0.47 | 1.00E+07 | 8.46E+04 | 1.23E+07 | 1.04E+05 | 576 | 1232.31 | 2133.99 | 1 | 485868 | 67.59 | 32.41 | 0.48 | 2.68 | 64.43 | 98.45 | 0.52 | 0.14 | 0.12 | 0.76 | 5 | 80 | 10 | 2440 | 94 | 8706 | 157 |
| | 6 | 4 | 638.4 | 0.16 | 2.02E+07 | 1.70E+05 | 3.81E+07 | 3.21E+05 | 988 | 1888.71 | 2623.35 | 1 | 888591 | 56.09 | 43.91 | 7.2 | 14.68 | 34.22 | 88.03 | 3.33 | 0.09 | 0.41 | 8.14 | 4 | 82 | 20 | 2677 | 178 | 9457 | 222 |
| | 18 | 4 | 816 | 0.2 | 8.42E+05 | 7.13E+03 | 1.25E+06 | 1.06E+04 | 535 | 1489.91 | 2675.87 | 1 | 358254 | 58.64 | 41.36 | 1.62 | 3.92 | 53.1 | 97.44 | 0.85 | 0.11 | 0.12 | 1.48 | 6 | 65 | 12 | 1932 | 124 | 5917 | 171 |
| | 30 | 4 | 2870.4 | 0.72 | 2.33E+07 | 1.90E+05 | 5.12E+07 | 4.18E+05 | 761 | 2193.78 | 3491.03 | 1 | 515440 | 59.84 | 40.16 | 0.75 | 2.66 | 56.43 | 98.31 | 0.65 | 0.02 | 0.11 | 0.9 | 9 | 81 | 9 | 2620 | 111 | 9370 | 155 |
| G1 | | 4 | 2371.2 | 0.79 | 1.38E+06 | 1.49E+04 | 1.15E+06 | 1.25E+04 | 511 | 836.17 | 1127.72 | 33 | 127252 | 84.98 | 15.02 | 0.08 | 0.58 | 84.32 | 99.75 | 0.1 | 0.01 | 0.02 | 0.11 | 2 | 65 | 11 | 1849 | 52 | 5355 | 91 |
| G2 | | 1 | 5280 | 5.28 | 5.70E+06 | 3.81E+04 | 1.07E+07 | 7.18E+04 | 666 | 1883.58 | 3475.69 | 10 | 273342 | 83.43 | 16.57 | 0.22 | 0.94 | 82.26 | 94.6 | 0.18 | 5.01 | 0.02 | 0.19 | 7 | 75 | 10 | 2268 | 58 | 8020 | 96 |
| G3 | | 1 | 12096 | 12.1 | 8.09E+06 | 5.68E+04 | 1.59E+07 | 1.11E+05 | 516 | 1959.31 | 4346.77 | 1 | 139428 | 64.53 | 35.47 | 0.57 | 1.75 | 62.2 | 98.4 | 0.45 | 0.46 | 0.08 | 0.61 | 7 | 80 | 13 | 2485 | 109 | 9074 | 176 |
| G4 | | 1 | 4896 | 4.9 | 1.50E+07 | 1.06E+05 | 1.13E+07 | 8.00E+04 | 422 | 755.4 | 1300.16 | 1 | 68018 | 61.82 | 38.18 | 0.33 | 1.61 | 59.88 | 98.95 | 0.36 | 0.15 | 0.07 | 0.48 | 2 | 81 | 10 | 2623 | 110 | 9383 | 161 |
| G5 | | 1 | 9168 | 9.17 | 8.51E+06 | 5.85E+04 | 1.49E+07 | 1.03E+05 | 558 | 1752.93 | 3198.55 | 19 | 291117 | 71.02 | 28.98 | 0.38 | 1.22 | 69.42 | 99.26 | 0.35 | 0.08 | 0.03 | 0.28 | 10 | 80 | 8 | 2324 | 62 | 7979 | 98 |
| G6 | | 1 | 5040 | 5.04 | 1.21E+07 | 8.45E+04 | 1.30E+07 | 9.08E+04 | 454 | 1074.8 | 2438.93 | 9 | 617076 | 78.35 | 21.65 | 0.32 | 0.76 | 77.27 | 99.68 | 0.13 | 0.01 | 0.01 | 0.16 | 4 | 80 | 6 | 2317 | 41 | 8165 | 76 |
| G7 | | 1 | 5184 | 5.18 | 1.43E+07 | 9.67E+04 | 1.06E+07 | 7.14E+04 | 389 | 738.08 | 1463.07 | 2 | 77266 | 45.33 | 54.67 | 0.57 | 1.98 | 42.78 | 97.28 | 0.94 | 1 | 0.07 | 0.72 | 4 | 80 | 13 | 2501 | 99 | 8961 | 162 |
| G8 | | 1 | 5952 | 5.95 | 1.33E+07 | 8.91E+04 | 1.30E+07 | 8.73E+04 | 415 | 979.24 | 2256.87 | 20 | 88473 | 85.24 | 14.76 | 0.35 | 0.58 | 84.32 | 99.77 | 0.09 | 0.02 | 0.01 | 0.11 | 2 | 79 | 5 | 2310 | 51 | 8164 | 94 |
| G9 | | 1 | 12864 | 12.86 | 4.88E+06 | 3.19E+04 | 1.10E+07 | 7.18E+04 | 658 | 2250.17 | 4460.02 | 26 | 127789 | 71.78 | 28.22 | 1.9 | 1.62 | 68.26 | 98.6 | 0.38 | 0.15 | 0.06 | 0.81 | 8 | 76 | 13 | 2379 | 80 | 8512 | 143 |
| G10 | | 1 | 7728 | 7.73 | 9.27E+06 | 6.61E+04 | 1.38E+07 | 9.85E+04 | 500 | 1488.9 | 3147.88 | 1 | 710776 | 57.85 | 42.15 | 0.49 | 2.04 | 55.32 | 98.24 | 0.49 | 0.57 | 0.07 | 0.62 | 7 | 81 | 13 | 2473 | 110 | 8927 | 168 |
| G11 | | 1 | 9984 | 9.98 | 7.14E+06 | 4.96E+04 | 1.47E+07 | 1.02E+05 | 700 | 2052.25 | 3878.55 | 1 | 81852 | 61.1 | 38.9 | 0.73 | 3.58 | 56.79 | 97.75 | 0.64 | 0.36 | 0.1 | 1.16 | 10 | 82 | 17 | 2550 | 117 | 9132 | 184 |
