## Supplementary Table 4 for "MetaGut: Insights into gut microbiomes in stem cell transplantation by comprehensive shotgun long-read sequencing"

| Category | Group 1 | Group 2 | p (Mann Whitney U) | |
|---|---|---|---|---|
| ARG-carrying Reads Per 10,000 Reads | **Pre TX** | **Healthy** | **0.00013** | ** |
| | Pre TX | Leukopenia | 0.87716 | N.s. |
| | Pre TX | Reconstitution | 0.10619 | N.s. |
| | Leukopenia | Healthy | 0.00548 | N.s. |
| | Leukopenia | Reconstitution | 0.22272 | N.s. |
| | Reconstitution | Healthy | 0.00992 | N.s. |
| DNA [µg] Per g Stool | **Pre TX** | **Healthy** | **0.00009** | ** |
| | Pre TX | Leukopenia | 0.00907 | N.s. |
| | Pre TX | Reconstitution | 0.67998 | N.s. |
| | **Leukopenia** | **Healthy** | **0.00002** | *** |
| | **Leukopenia** | **Reconstitution** | **0.00068** | * |
| | **Reconstitution** | **Healthy** | **0.00011** | ** |
| Human [% Reads] | Pre TX | Healthy | 0.00303 | N.s. |
| | Pre TX | Leukopenia | 0.01240 | N.s. |
| | Pre TX | Reconstitution | 0.87127 | N.s. |
| | **Leukopenia** | **Healthy** | **0.00003** | ** |
| | Leukopenia | Reconstitution | 0.00681 | N.s. |
| | **Reconstitution** | **Healthy** | **0.00111** | * |
| Median Readlength | Pre TX | Healthy | 0.00166 | N.s. |
| | Pre TX | Leukopenia | 0.66913 | N.s. |
| | Pre TX | Reconstitution | 0.08931 | N.s. |
| | Leukopenia | Healthy | 0.05188 | N.s. |
| | Leukopenia | Reconstitution | 0.46518 | N.s. |
| | Reconstitution | Healthy | 0.06879 | N.s. |
| No. Genera | Pre TX | Healthy | 0.02323 | N.s. |
| | Pre TX | Leukopenia | 0.01592 | N.s. |
| | Pre TX | Reconstitution | 0.83656 | N.s. |
| | **Leukopenia** | **Healthy** | **0.00091** | * |
| | Leukopenia | Reconstitution | 0.03347 | N.s. |
| | Reconstitution | Healthy | 0.05394 | N.s. |
| Total Reads | **Pre TX** | **Healthy** | **0.00011** | ** |
| | Pre TX | Leukopenia | 0.05181 | N.s. |
| | Pre TX | Reconstitution | 0.35633 | N.s. |
| | **Leukopenia** | **Healthy** | **0.00004** | ** |
| | Leukopenia | Reconstitution | 0.09499 | N.s. |
| | **Reconstitution** | **Healthy** | **0.00019** | ** |
| Unclassified [% Reads] | Pre TX | Healthy | 0.00723 | N.s. |
| | Pre TX | Leukopenia | 0.08571 | N.s. |
| | Pre TX | Reconstitution | 0.22895 | N.s. |
| | **Leukopenia** | **Healthy** | **0.00086** | * |
| | Leukopenia | Reconstitution | 0.02062 | N.s. |
| | Reconstitution | Healthy | 0.05523 | N.s. |

| * | ** | *** |
|---|---|---|
| 1.19E-03 | 2.38E-04 | 2.38E-05 |
| 0.00116279 | 0.00023256 | 2.32558E-05 |
