## Supplementary Table 5 for "MetaGut: Insights into gut microbiomes in stem cell transplantation by comprehensive shotgun long-read sequencing"

| | Fraction Healthy (n=11) | Fraction Reconstitution (n=48) | Fraction Pre TX (n=32) | Fraction Leukozytopenia (n=21) | Fraction Total (n=112) |
|---|---|---|---|---|---|
| Bacteroides | 1 | 0.354166666666667 | 0.59375 | 0.095238095238095 | 0.4375 |
| Phocaeicola | 0.909090909090909 | 0.291666666666667 | 0.5 | 0.190476190476191 | 0.392857142857143 |
| Enterococcus | | 0.229166666666667 | 0.28125 | 0.571428571428571 | 0.285714285714286 |
| Parabacteroides | 0.272727272727273 | 0.166666666666667 | 0.34375 | 0.095238095238095 | 0.214285714285714 |
| Streptococcus | | 0.145833333333333 | 0.1875 | 0.380952380952381 | 0.1875 |
| Escherichia | | 0.125 | 0.1875 | 0.285714285714286 | 0.160714285714286 |
| Alistipes | 0.545454545454545 | 0.104166666666667 | 0.125 | 0.047619047619048 | 0.142857142857143 |
| Enterocloster | | 0.229166666666667 | 0.15625 | | 0.142857142857143 |
| Faecalibacterium | 0.454545454545454 | 0.125 | 0.125 | | 0.133928571428571 |
| Blautia | | 0.166666666666667 | 0.1875 | | 0.125 |
| Roseburia | 0.090909090909091 | 0.1875 | 0.09375 | | 0.116071428571429 |
| Lactobacillus | | 0.020833333333333 | 0.15625 | 0.285714285714286 | 0.107142857142857 |
| Flavonifractor | | 0.125 | 0.125 | 0.095238095238095 | 0.107142857142857 |
| Citrobacter | | 0.083333333333333 | 0.09375 | 0.095238095238095 | 0.080357142857143 |
| Erysipelatoclostridium | | 0.083333333333333 | 0.03125 | 0.142857142857143 | 0.071428571428572 |
| Stenotrophomonas | | 0.0625 | 0.0625 | 0.095238095238095 | 0.0625 |
| Proteus | | 0.0625 | 0.03125 | 0.095238095238095 | 0.053571428571429 |
| Veillonella | | 0.104166666666667 | | | 0.044642857142857 |
| Dysosmobacter | | 0.083333333333333 | 0.03125 | | 0.044642857142857 |
| Klebsiella | | 0.083333333333333 | 0.03125 | | 0.044642857142857 |
| Mediterraneibacter | | 0.083333333333333 | 0.03125 | | 0.044642857142857 |
| Providencia | | 0.020833333333333 | 0.03125 | 0.095238095238095 | 0.035714285714286 |
| Lacticaseibacillus | | 0.020833333333333 | 0.03125 | 0.095238095238095 | 0.035714285714286 |
| Schaalia | | 0.0625 | 0.03125 | | 0.035714285714286 |
| Clostridium | | 0.0625 | | 0.047619047619048 | 0.035714285714286 |
| Pediococcus | | | 0.03125 | 0.095238095238095 | 0.026785714285714 |
| Staphylococcus | | 0.041666666666667 | | 0.047619047619048 | 0.026785714285714 |
| Lactococcus | | 0.020833333333333 | 0.03125 | 0.047619047619048 | 0.026785714285714 |
| Subdoligranulum | | 0.041666666666667 | 0.03125 | | 0.026785714285714 |
| Simiaoa | 0.090909090909091 | 0.041666666666667 | | | 0.026785714285714 |
| Ruminococcus | 0.090909090909091 | 0.041666666666667 | | | 0.026785714285714 |
| Rothia | | 0.041666666666667 | 0.03125 | | 0.026785714285714 |
| Akkermansia | 0.090909090909091 | 0.041666666666667 | | | 0.026785714285714 |
| Vescimonas | | 0.020833333333333 | 0.03125 | | 0.017857142857143 |
| Hungatella | | 0.041666666666667 | | | 0.017857142857143 |
| Ruthenibacterium | | 0.041666666666667 | | | 0.017857142857143 |
| Lachnoclostridium | | 0.020833333333333 | 0.03125 | | 0.017857142857143 |
| Ligilactobacillus | | | | 0.095238095238095 | 0.017857142857143 |
| Prevotella | 0.181818181818182 | | | | 0.017857142857143 |
| Intestinimonas | | | | 0.047619047619048 | 0.008928571428571 |
| Cutibacterium | | | | 0.047619047619048 | 0.008928571428571 |
| Catenibacterium | | | | 0.047619047619048 | 0.008928571428571 |
| Lachnospira | | 0.020833333333333 | | | 0.008928571428571 |
| Flintibacter | | | 0.03125 | | 0.008928571428571 |
| Bifidobacterium | | | 0.03125 | | 0.008928571428571 |
| Anaerostipes | | | 0.03125 | | 0.008928571428571 |
| Cellulosilyticum | | 0.020833333333333 | | | 0.008928571428571 |
| Wujia | 0.090909090909091 | | | | 0.008928571428571 |
| Pseudomonas | | | | 0.047619047619048 | 0.008928571428571 |
