## Supplementary Table 6 for "MetaGut: Insights into gut microbiomes in stem cell transplantation by comprehensive shotgun long-read sequencing"

| | Fraction Leukozytopenia (n=21) | Fraction Pre TX (n=32) | Fraction Reconstitution (n=48) | Fraction Healthy (n=11) | Fraction Total (n=112) | Fraction Cluster 1 (n=18) | Fraction Cluster 2 (n=8) | Fraction Cluster 3 (n=6) |
|---|---|---|---|---|---|---|---|---|
| Saccharomyces | 0.714285714285714 | 0.53125 | 0.541666666666667 | 0.272727272727273 | 0.544642857142857 | 0.388888888888889 | 0.5 | 1 |
| Candida | 0.142857142857143 | 0.09375 | 0.083333333333333 | | 0.089285714285714 | | 0.25 | 0.166666666666667 |
| Cyberlindnera | 0.047619047619048 | 0.03125 | | | 0.017857142857143 | | 0.125 | |
| Nakaseomyces | | | 0.041666666666667 | | 0.017857142857143 | | | |
| Alternaria | 0.047619047619048 | | | | 0.008928571428571 | | | |
| Malassezia | 0.047619047619048 | | | | 0.008928571428571 | | | |
| Debaryomyces | | 0.03125 | | | 0.008928571428571 | 0.055555555555556 | | |
