## Supplementary Table 7 for "MetaGut: Insights into gut microbiomes in stem cell transplantation by comprehensive shotgun long-read sequencing"

| | Fraction Healthy (n=11) | Fraction Pre TX (n=32) | Fraction Leukozytopenia (n=21) | Fraction Reconstitution (n=48) | Fraction Total (n=112) | Fraction Cluster 1 (n=18) | Fraction Cluster 2 (n=8) | Fraction Cluster 3 (n=6) |
|---|---|---|---|---|---|---|---|---|
| Skunavirus | | 0.625 | 0.428571428571429 | 0.333333333333333 | 0.401785714285714 | 0.5 | 0.625 | 1 |
| Crassphage Pseudo-Genus | 0.090909090909091 | 0.1875 | 0.095238095238095 | 0.104166666666667 | 0.125 | 0.277777777777778 | | 0.166666666666667 |
| Afonbuvirus | 0.090909090909091 | 0.03125 | 0.047619047619048 | 0.041666666666667 | 0.044642857142857 | 0.055555555555556 | | |
| Taranisvirus | 0.181818181818182 | 0.03125 | | 0.020833333333333 | 0.035714285714286 | 0.055555555555556 | | |
| Oengusvirus | 0.272727272727273 | 0.03125 | | | 0.035714285714286 | 0.055555555555556 | | |
| Vectrevirus | 0.272727272727273 | | | | 0.026785714285714 | | | |
| Cohcovirus | 0.181818181818182 | 0.03125 | | | 0.026785714285714 | 0.055555555555556 | | |
| Brigitvirus | 0.090909090909091 | 0.03125 | | 0.020833333333333 | 0.026785714285714 | 0.055555555555556 | | |
| Jahgtovirus | 0.181818181818182 | | | | 0.017857142857143 | | 0.125 | |
| Novemvirus | | 0.03125 | | 0.020833333333333 | 0.017857142857143 | | | 0.333333333333333 |
| Efquatrovirus | | 0.0625 | | | 0.017857142857143 | | | |
| Blohavirus | 0.090909090909091 | | | 0.020833333333333 | 0.017857142857143 | 0.055555555555556 | | |
| Toutatisvirus | | 0.03125 | | 0.020833333333333 | 0.017857142857143 | | | |
| Moineauvirus | | | | 0.041666666666667 | 0.017857142857143 | | | |
| Canhaevirus | 0.090909090909091 | 0.03125 | | | 0.017857142857143 | 0.055555555555556 | | |
| Ceduovirus | 0.090909090909091 | | | 0.020833333333333 | 0.017857142857143 | | | |
| Kahucivirus | | | | 0.020833333333333 | 0.008928571428571 | | | 0.166666666666667 |
| Felixounavirus | | | | 0.020833333333333 | 0.008928571428571 | | | |
| Lentivirus | | | | 0.020833333333333 | 0.008928571428571 | | | |
| Vedamuthuvirus | | 0.03125 | | | 0.008928571428571 | | | |
| Betapolyomavirus | | | | 0.020833333333333 | 0.008928571428571 | | | |
| Betacoronavirus | | | | 0.020833333333333 | 0.008928571428571 | | | |
| Webervirus | | 0.03125 | | | 0.008928571428571 | 0.055555555555556 | | |
| Birpovirus | | | | 0.020833333333333 | 0.008928571428571 | | | |
| Dhillonvirus | | 0.03125 | | | 0.008928571428571 | 0.055555555555556 | | |
| Kahnovirus | | 0.03125 | | | 0.008928571428571 | 0.055555555555556 | | |
| Junavirus | | 0.03125 | | | 0.008928571428571 | | | 0.166666666666667 |
| Drulisvirus | | 0.03125 | | | 0.008928571428571 | | 0.125 | |
| Culoivirus | | 0.03125 | | | 0.008928571428571 | 0.055555555555556 | | |
| Buchavirus | | 0.03125 | | | 0.008928571428571 | 0.055555555555556 | | |
| Aurodevirus | | 0.03125 | | | 0.008928571428571 | 0.055555555555556 | | |
| Ashduovirus | | 0.03125 | | | 0.008928571428571 | | | 0.166666666666667 |
| Burzaovirus | 0.090909090909091 | | | | 0.008928571428571 | | | |
| Warwickvirus | | | | 0.020833333333333 | 0.008928571428571 | | | |
