## Supplementary Table 8 for "MetaGut: Insights into gut microbiomes in stem cell transplantation by comprehensive shotgun long-read sequencing"

| Patient ID | Age at Tx | Sex | Pre-Tx Cluster | Diagnosis | Month between First Diagnosis and Tx | No. Chemotherapy Cycles | No. Non-Cytostatic Pre-Therapies | Remission state prior to Tx | Immuno-suppression | Conditioning Regime | HLA-Match | Donor |
|---|---|---|---|---|---|---|---|---|---|---|---|---|
| 4 | 65 | 1 | 1 | 1 | 3 | 3 | 0 | 3 | 1 | 0 | 0 | 1 |
| 7 | 51 | 1 | 1 | 1 | 4 | 0 | 0 | 3 | 1 | 0 | 0 | 1 |
| 11 | 50 | 0 | 1 | 2 | 20 | 14 | 1 | 1 | 1 | 0 | 0 | 1 |
| 12 | 56 | 0 | 1 | 1 | 5 | 2 | 0 | 1 | 1 | 0 | 0 | 1 |
| 13 | 58 | 1 | 2 | 1 | 3 | 3 | 0 | 1 | 1 | 0 | 0 | 1 |
| 14 | 66 | 0 | 1 | 1 | 8 | 4 | 1 | 3 | 1 | 0 | 0 | 1 |
| 15 | 55 | 0 | 1 | 1 | 3 | 2 | 0 | 1 | 1 | 0 | 0 | 1 |
| 16 | 68 | 1 | 1 | 1 | 9 | 2 | 0 | 3 | 1 | 0 | 0 | 1 |
| 17 | 59 | 0 | 1 | 1 | 4 | 1 | 0 | 3 | 1 | 0 | 0 | 1 |
| 18 | 60 | 0 | 2 | 1 | 19 | 5 | 0 | 3 | 1 | 0 | 0 | 1 |
| 18.2 | 60 | 0 | 3 | 1 | 19 | 6 | 0 | 3 | 1 | 0 | 0 | 1 |
| 19 | 57 | 0 | 1 | 2 | 8 | 4 | 0 | 1 | 2 | 1 | 1 | 1 |
| 20 | 35 | 0 | 3 | 2 | 13 | 3 | 2 | 2 | 2 | 1 | 1 | 0 |
| 21 | 71 | 0 | 3 | 1 | 2 | 1 | 0 | 3 | 1 | 0 | 0 | 1 |
| 22 | 68 | 1 | 2 | 3 | 10 | 0 | 1 | 3 | 2 | 1 | 0 | 0 |
| 23 | 54 | 1 | 2 | 1 | 8 | 5 | 1 | 3 | 1 | 0 | 0 | 1 |
| 24 | 53 | 0 | 1 | 1 | 24 | 0 | 1 | 3 | 1 | 0 | 0 | 1 |
| 25 | 45 | 1 | 2 | 1 | 13 | 6 | 0 | 1 | 1 | 0 | 0 | 1 |
| 26 | 35 | 1 | 3 | 1 | 4 | 3 | 0 | 1 | 1 | 0 | 0 | 1 |
| 28 | 52 | 1 | 1 | 1 | 5 | 2 | 0 | 2 | 1 | 0 | 0 | 1 |
| 29 | 48 | 0 | 1 | 3 | 4 | 11 | 0 | 3 | 2 | 1 | 1 | 0 |
| 31 | 19 | 0 | 3 | 1 | 28 | 0 | 0 | 3 | 1 | 0 | 0 | 1 |
| 32 | 69 | 0 | 1 | 1 | 5 | 0 | 3 | 3 | 1 | 0 | 0 | 1 |
| 33 | 60 | 0 | 3 | 3 | 3 | 14 | 1 | 1 | 1 | 0 | 0 | 1 |
| 34 | 48 | 0 | 1 | 1 | 78 | 0 | 0 | 3 | 1 | 0 | 0 | 1 |
| 36 | 65 | 1 | 3 | 1 | 18 | 4 | 0 | 1 | 2 | 1 | 1 | 0 |
| 37 | 44 | 0 | 3 | 2 | 39 | 6 | 1 | 1 | 1 | 1 | 1 | 0 |
| 38 | 60 | 1 | 1 | 1 | 5 | 2 | 0 | 1 | 1 | 0 | 0 | 1 |
| 39 | 65 | 0 | 1 | 1 | 8 | 0 | 0 | 3 | 1 | 0 | 0 | 1 |
| 40 | 73 | 0 | 2 | 1 | 9 | 4 | 2 | 1 | 1 | 0 | 0 | 1 |
| 41 | 60 | 1 | 2 | 1 | 25 | 10 | 6 | 3 | 1 | 0 | 0 | 1 |
| 42 | 67 | 0 | 1 | 3 | 19 | 0 | 0 | 3 | 1 | 0 | 0 | 1 |

| Pat ID | Age at transplantation | Sex | Pre-transplantation cluster | Colonisation (0 = none; 1 = 3-MRGN; 2 = VRE; 3 = 4-MRGN) | Diagnosis: AML (1), ALL (2), other myeloic neoplasia (3) | Time first diagnosis to HSCT [month] | Number chemotherapy cycles | Number non-cytostatic pre-therapies | Remission state prior to HSCT (1 = MRD-, 2 = hematological complete remission, MRD+, 3 = hematological active disease) | Immunosuppression (ATG = 1, PostCy = 2) | Type of conditioning regimen (TBI = 1, Chemotherapy = 0) | HLA-match (HLA-identical = 0; non-identical = 1) | donor (related = 0; non-related = 1) |
|---|---|---|---|---|---|---|---|---|---|---|---|---|---|
| 4 | 65 | 1 | 1 | 0 | 1 | 3 | 3 | 0 | 3 | 1 | 0 | 0 | 1 |
| 7 | 51 | 1 | 1 | 0 | 1 | 4 | 0 | 0 | 3 | 1 | 0 | 0 | 1 |
| 11 | 50 | 0 | 1 | 0 | 2 | 20 | 14 | 1 | 1 | 1 | 0 | 0 | 1 |
| 12 | 56 | 0 | 1 | 0 | 1 | 5 | 2 | 0 | 1 | 1 | 0 | 0 | 1 |
| 13 | 58 | 1 | 2 | 0 | 1 | 3 | 3 | 0 | 1 | 1 | 0 | 0 | 1 |
| 14 | 66 | 0 | 1 | 0 | 1 | 8 | 4 | 1 | 3 | 1 | 0 | 0 | 1 |
| 15 | 55 | 0 | 1 | 0 | 1 | 3 | 2 | 0 | 1 | 1 | 0 | 0 | 1 |
| 16 | 68 | 1 | 1 | 0 | 1 | 9 | 2 | 0 | 3 | 1 | 0 | 0 | 1 |
| 17 | 59 | 0 | 1 | 0 | 1 | 4 | 1 | 0 | 3 | 1 | 0 | 0 | 1 |
| 18 | 60 | 0 | 2 | 0 | 1 | 19 | 5 | 0 | 3 | 1 | 0 | 0 | 1 |
| 18.2 | 60 | 0 | 3 | 0 | 1 | 19 | 6 | 0 | 3 | 1 | 0 | 0 | 1 |
| 19 | 57 | 0 | 1 | 0 | 2 | 8 | 4 | 0 | 1 | 2 | 1 | 1 | 1 |
| 20 | 35 | 0 | 3 |  | 2 | 13 | 3 | 2 | 2 | 2 | 1 | 1 | 0 |
| 21 | 71 | 0 | 3 | 0 | 1 | 2 | 1 | 0 | 3 | 1 | 0 | 0 | 1 |
| 22 | 68 | 1 | 2 | 1+3 | 3 | 10 | 0 | 1 | 3 | 2 | 1 | 0 | 0 |
| 23 | 54 | 1 | 2 | 0 | 1 | 8 | 5 | 1 | 3 | 1 | 0 | 0 | 1 |
| 24 | 53 | 0 | 1 | 0 | 1 | 24 | 0 | 1 | 3 | 1 | 0 | 0 | 1 |
| 25 | 45 | 1 | 2 | 0 | 1 | 13 | 6 | 0 | 1 | 1 | 0 | 0 | 1 |
| 26 | 35 | 1 | 3 | 1 | 1 | 4 | 3 | 0 | 1 | 1 | 0 | 0 | 1 |
| 28 | 52 | 1 | 1 | 0 | 1 | 5 | 2 | 0 | 2 | 1 | 0 | 0 | 1 |
| 29 | 48 | 0 | 1 | 0 | 3 | 4 | 11 | 0 | 3 | 2 | 1 | 1 | 0 |
| 31 | 19 | 0 | 3 | 0 | 1 | 28 | 0 | 0 | 3 | 1 | 0 | 0 | 1 |
| 32 | 69 | 0 | 1 |  | 1 | 5 | 0 | 3 | 3 | 1 | 0 | 0 | 1 |
| 33 | 60 | 0 | 3 | 0 | 3 | 3 | 14 | 1 | 1 | 1 | 0 | 0 | 1 |
| 34 | 48 | 0 | 1 | 2 | 1 | 78 | 0 | 0 | 3 | 1 | 0 | 0 | 1 |
| 36 | 65 | 1 | 3 | 1 | 1 | 18 | 4 | 0 | 1 | 2 | 1 | 1 | 0 |
| 37 | 44 | 0 | 3 | 1 | 2 | 39 | 6 | 1 | 1 | 1 | 1 | 1 | 0 |
| 38 | 60 | 1 | 1 | 3 | 1 | 5 | 2 | 0 | 1 | 1 | 0 | 0 | 1 |
| 39 | 65 | 0 | 1 | 0 | 1 | 8 | 0 | 0 | 3 | 1 | 0 | 0 | 1 |
| 40 | 73 | 0 | 2 | 0 | 1 | 9 | 4 | 2 | 1 | 1 | 0 | 0 | 1 |
| 41 | 60 | 1 | 2 | 0 | 1 | 25 | 10 | 6 | 3 | 1 | 0 | 0 | 1 |
| 42 | 67 | 0 | 1 | 0 | 3 | 19 | 0 | 0 | 3 | 1 | 0 | 0 | 1 |
