## Supplementary Table 9 for "MetaGut: Insights into gut microbiomes in stem cell transplantation by comprehensive shotgun long-read sequencing"

| Reported Marker Family / Genus / Spezies | Literature | | | | |
|---|---|---|---|---|---|
| Akkermansia | Ilett et al., 2020 | Shono et al., 2016 | Ugrayová et al., 2022 | | |
| Bacteroides | Doki et al., 2017 | Golob et al., 2017 | Simms-Walrip et al., 2017 | Marandi et al., 2022 | |
| Bifidobacterium | Simms-Walrip et al., 2017 | Kusakabe et al., 2020 | Rajilic-Stojanovic et al., 2011 | | |
| Blautia | Jenq et al., 2015 | Golob et al., 2017 | Weber et al., 2017 | Ilett et al., 2020 | Rajilic-Stojanov[illegible] |
| Candida (parapsilosis) | Rolling et al., 2021 | Kumamoto et al., 2020 | | | |
| Clostridium | Weber et al., 2017 | Jenq et al., 2012 | Rajilic-Stojanovic et al., 2011 | | |
| crAssphage | Shkoporov et al., 2019 | Guerin et al., 2018 | | | |
| Dorea | Golob et al., 2017 | Rajilic-Stojanovic et al., 2011 | | | |
| Enterobacter | Ugrayová et al., 2022 | | | | |
| Enterobacteroaceaea (z.B. Citrobacter, Enterobacter, Klebsiella, Escherichia) | Simms-Walrip et al., 2017 | | | | |
| Enterococcus | Holler et al., 2014 | Simms-Walrip et al., 2017 | Ugrayová et al., 2022 | | |
| Escherichia | Lee at al., 2019 | | | | |
| Eubacterium | Doki et al., 2017 | Lee at al., 2019 | | | |
| Faecalibacterium | Doki et al., 2017 | Kusakabe et al., 2020 | Rajilic-Stojanovic et al., 2011 | | |
| Klebsiella | Ugrayová et al., 2022 | | | | |
| Prevotella | Marandi et al., 2022 | Willing et al., 2010 | | | |
| Ruminococcus | Lee at al., 2019 | Golob et al., 2017 | Ingham et al., 2019 | Rajilic-Stojanovic et al., 2011 | |
| Sutterella | Kusakabe et al., 2020 | | | | |
