## Supplementary Table 10 for "MetaGut: Insights into gut microbiomes in stem cell transplantation by comprehensive shotgun long-read sequencing"

| File | otal Length (Gbp) | Sequences |
|---|---|---|
| Fungi | 14.955 | 73,001 |
| Archaea | 1.400 | 894 |
| Human | 3.298 | 705 |
| Viral | 0.467 | 14,758 |
| Protozoa | 3.618 | 216,550 |
| Plant | 119.067 | 1,026,960 |
| Plasmid | 4.531 | 47,953 |
| Bacteria | 142.086 | 80,596 |
| adds/Cladosporium_Cladosporioides_GCA_002901145.1_ASM290114v1_geno | 0.033 | 67 |
| adds/Candida_Lusitaniae_GCF_000003835.1_ASM383v1_genomic.fna | 0.012 | 9 |
| adds/Galactomyces_GCA_025134825.1_LMA-1150_v1_genomic.fna | 0.025 | 2,700 |
| adds/Cryptosporidium_Meleagridis_GCA_004348035.1_UKMEL3.v0_genomic.f | 0.009 | 2,260 |
| adds/Entamoeba_Dispar_GCF_000209125.1_JCVI_EDISG_1.0_genomic.fna | 0.031 | 12,258 |
| adds/Mucor_Circinelloides_GCA_023629755.1_ASM2362975v1_genomic.fna | 0.033 | 1,948 |
| adds/GCF_000001635.27_GRCm39_genomic.fna (Mus) | 2.728 | 61 |
| | 292.294 | 1480720 |
