## Supplementary Table 11 for "MetaGut: Insights into gut microbiomes in stem cell transplantation by comprehensive shotgun long-read sequencing"

| Kraken2 Assigned Genus | Zymo Gut Microbiome Standard Genus | Reads | Expected Abundance | Expected Reads | Zymo References: Validation Rate | Zymo References: Validated Reads | Zymo References: Non-Validated Reads | FastANI Zymo References vs Comprehensive Database: ANI Estimate | FastANI: Orthologous Matches | FastANI: Sequence Fragments | MetaGut v. 1.0 Database: Validation Rate | MetaGut: Validated Reads | MetaGut: Non-Validated Reads | | |
|---|---|---|---|---|---|---|---|---|---|---|---|---|---|---|---|
| *Bifidobacterium* | TRUE | 239 | 0.060000 | 5422 | 0.937 | 224 | 15 | 99.976 | 701 | 193079 | 0.946 | 226 | 13 | | |
| *Salmonella* | TRUE | 79 | 0.000100 | 9 | 0.734 | 58 | 21 | 99.969 | 1597 | 2310229 | 0.911 | 72 | 7 | **Pearson r Validation Rate vs ANI Esti** | 0.3738722334 |
| *Escherichia* | TRUE | 17230 | 0.140000 | 12651 | 0.984 | 16951 | 279 | 99.925 | 7717 | 4655965 | 0.983 | 16942 | 288 | **t** | 1.453416993 |
| *Akkermansia* | TRUE | 1798 | 0.015000 | 1355 | 0.983 | 1768 | 30 | 99.844 | 956 | 94520 | 0.979 | 1760 | 38 | **p** | 0.1698193994 |
| *Enterococcus* | TRUE | 7 | 0.000010 | 1 | 0.000 | 0 | 7 | 99.820 | 954 | 537258 | 0.286 | 2 | 5 | | |
| *Clostridioides* | TRUE | 1890 | 0.015000 | 1355 | 0.943 | 1783 | 107 | 99.623 | 1412 | 204770 | 0.933 | 1764 | 126 | | |
| *Methanobrevibacter* | TRUE | 34 | 0.001000 | 90 | 0.824 | 28 | 6 | 99.517 | 620 | 7841 | 0.824 | 28 | 6 | | |
| *Bacteroides* | TRUE | 18057 | 0.140000 | 12651 | 0.978 | 17656 | 401 | 99.059 | 1715 | 231969 | 0.942 | 17014 | 1043 | | |
| *Clostridium* | TRUE | 22 | 0.000001 | 0 | 0.000 | 0 | 22 | 98.659 | 1149 | 311472 | 0.182 | 4 | 18 | | |
| *Candida* | TRUE | 449 | 0.015000 | 1355 | 0.497 | 223 | 226 | 98.632 | 2020 | 27264 | 0.940 | 422 | 27 | | |
| *Saccharomyces* | TRUE | 588 | 0.014000 | 1265 | 0.980 | 576 | 12 | 98.432 | 4027 | 11967 | 0.912 | 536 | 52 | | |
| *Faecalibacterium* | TRUE | 19006 | 0.140000 | 12651 | 0.965 | 18332 | 674 | 97.602 | 966 | 23836 | 0.909 | 17285 | 1721 | | |
| *Roseburia* | TRUE | 1796 | 0.140000 | 12651 | 0.961 | 1726 | 70 | 97.349 | 1064 | 10876 | 0.901 | 1618 | 178 | | |
| *Fusobacterium* | TRUE | 4854 | 0.060000 | 5422 | 0.933 | 4530 | 324 | 97.319 | 820 | 49764 | 0.811 | 3937 | 917 | | |
| *Veillonella* | TRUE | 15369 | 0.140000 | 12651 | 0.993 | 15268 | 101 | 91.772 | 698 | 7760 | 0.841 | 12928 | 2441 | | |
| *Prevotella* | TRUE | 4710 | 0.060000 | 5422 | 0.990 | 4661 | 49 | 81.105 | 723 | 81136 | 0.383 | 1803 | 2907 | | |
| *Lactobacillus* | TRUE | 0 | 0.060000 | 5422 | | | | 80.113 | 187 | 114333 | | | | | |
| Nakaseomyces | FALSE | 1 | | | | | | | | | 1.000 | 1 | 0 | | |
| Latilactobacillus | FALSE | 1 | | | | | | | | | 1.000 | 1 | 0 | | |
| Kazachstania | FALSE | 1 | | | | | | | | | 1.000 | 1 | 0 | | |
| Botrytis | FALSE | 1 | | | | | | | | | 1.000 | 1 | 0 | | |
| Weissella | FALSE | 1 | | | | | | | | | 1.000 | 1 | 0 | | |
| Gemella | FALSE | 1 | | | | | | | | | 1.000 | 1 | 0 | | |
| Sugiyamaella | FALSE | 1 | | | | | | | | | 1.000 | 1 | 0 | | |
| Monoglobus | FALSE | 1 | | | | | | | | | 1.000 | 1 | 0 | | |
| Betacoronavirus | FALSE | 3 | | | | | | | | | 1.000 | 3 | 0 | | |
| Marvinbryantia | FALSE | 1 | | | | | | | | | 1.000 | 1 | 0 | | |
| Pseudocitrobacter | FALSE | 5 | | | | | | | | | 1.000 | 5 | 0 | | |
| Shimwellia | FALSE | 1 | | | | | | | | | 1.000 | 1 | 0 | | |
| Yokenella | FALSE | 3 | | | | | | | | | 1.000 | 3 | 0 | | |
| Flintibacter | FALSE | 1 | | | | | | | | | 1.000 | 1 | 0 | | |
| Ignavigranum | FALSE | 1 | | | | | | | | | 1.000 | 1 | 0 | | |
| Pusillibacter | FALSE | 13 | | | | | | | | | 1.000 | 13 | 0 | | |
| Meyerozyma | FALSE | 1 | | | | | | | | | 1.000 | 1 | 0 | | |
| Atlantibacter | FALSE | 4 | | | | | | | | | 1.000 | 4 | 0 | | |
| Anaerobutyricum | FALSE | 1 | | | | | | | | | 1.000 | 1 | 0 | | |
| Naumovozyma | FALSE | 1 | | | | | | | | | 1.000 | 1 | 0 | | |
| Companilactobacillus | FALSE | 1 | | | | | | | | | 1.000 | 1 | 0 | | |
| Cutibacterium | FALSE | 4 | | | | | | | | | 1.000 | 4 | 0 | | |
| Shigella | FALSE | 218 | | | | | | | | | 0.968 | 211 | 7 | | |
| Vescimonas | FALSE | 20 | | | | | | | | | 0.950 | 19 | 1 | | |
| Limosilactobacillus | FALSE | 1948 | | | | | | | | | 0.930 | 1811 | 137 | | |
| Lactococcus | FALSE | 10 | | | | | | | | | 0.900 | 9 | 1 | | |
| Alistipes | FALSE | 65 | | | | | | | | | 0.877 | 57 | 8 | | |
| Leclercia | FALSE | 8 | | | | | | | | | 0.875 | 7 | 1 | | |
| Citrobacter | FALSE | 102 | | | | | | | | | 0.863 | 88 | 14 | | |
| Enterobacter | FALSE | 78 | | | | | | | | | 0.821 | 64 | 14 | | |
| Kosakonia | FALSE | 15 | | | | | | | | | 0.800 | 12 | 3 | | |
| Klebsiella | FALSE | 78 | | | | | | | | | 0.795 | 62 | 16 | | |
| Porphyromonas | FALSE | 51 | | | | | | | | | 0.745 | 38 | 13 | | |
| Simiaoa | FALSE | 22 | | | | | | | | | 0.727 | 16 | 6 | | |
| Mediterraneibacter | FALSE | 7 | | | | | | | | | 0.714 | 5 | 2 | | |
| Erysipelatoclostridium | FALSE | 3 | | | | | | | | | 0.667 | 2 | 1 | | |
| Cronobacter | FALSE | 3 | | | | | | | | | 0.667 | 2 | 1 | | |
| Buttiauxella | FALSE | 3 | | | | | | | | | 0.667 | 2 | 1 | | |
| Lelliottia | FALSE | 9 | | | | | | | | | 0.667 | 6 | 3 | | |
| Raoultella | FALSE | 8 | | | | | | | | | 0.625 | 5 | 3 | | |
| Faecalicatena | FALSE | 8 | | | | | | | | | 0.625 | 5 | 3 | | |
| Flavonifractor | FALSE | 13 | | | | | | | | | 0.615 | 8 | 5 | | |
| Kluyvera | FALSE | 5 | | | | | | | | | 0.600 | 3 | 2 | | |
| Actinomyces | FALSE | 5 | | | | | | | | | 0.600 | 3 | 2 | | |
| Parabacteroides | FALSE | 91 | | | | | | | | | 0.593 | 54 | 37 | | |
| Coprococcus | FALSE | 9 | | | | | | | | | 0.556 | 5 | 4 | | |
| Dysosmobacter | FALSE | 24 | | | | | | | | | 0.542 | 13 | 11 | | |
| Blautia | FALSE | 36 | | | | | | | | | 0.528 | 19 | 17 | | |
| Oscillibacter | FALSE | 4 | | | | | | | | | 0.500 | 2 | 2 | | |
| Jejubacter | FALSE | 6 | | | | | | | | | 0.500 | 3 | 3 | | |
| Wujia | FALSE | 2 | | | | | | | | | 0.500 | 1 | 1 | | |
| Lactiplantibacillus | FALSE | 4 | | | | | | | | | 0.500 | 2 | 2 | | |
| Scandinavium | FALSE | 2 | | | | | | | | | 0.500 | 1 | 1 | | |
| Brettanomyces | FALSE | 2 | | | | | | | | | 0.500 | 1 | 1 | | |
| Moraxella | FALSE | 2 | | | | | | | | | 0.500 | 1 | 1 | | |
| Muribaculum | FALSE | 2 | | | | | | | | | 0.500 | 1 | 1 | | |
| Eubacterium | FALSE | 6 | | | | | | | | | 0.500 | 3 | 3 | | |
| Symbiopectobacterium | FALSE | 4 | | | | | | | | | 0.500 | 2 | 2 | | |
| Suillus | FALSE | 2 | | | | | | | | | 0.500 | 1 | 1 | | |
| Hungatella | FALSE | 16 | | | | | | | | | 0.500 | 8 | 8 | | |
| Amedibacterium | FALSE | 2 | | | | | | | | | 0.500 | 1 | 1 | | |
| Anaerotruncus | FALSE | 8 | | | | | | | | | 0.500 | 4 | 4 | | |
| Phocaeicola | FALSE | 64 | | | | | | | | | 0.391 | 25 | 39 | | |
| Anaerostipes | FALSE | 3 | | | | | | | | | 0.333 | 1 | 2 | | |
| Butyrivibrio | FALSE | 3 | | | | | | | | | 0.333 | 1 | 2 | | |
| Lachnoclostridium | FALSE | 3 | | | | | | | | | 0.333 | 1 | 2 | | |
| Ornithobacterium | FALSE | 3 | | | | | | | | | 0.333 | 1 | 2 | | |
| Tannerella | FALSE | 3 | | | | | | | | | 0.333 | 1 | 2 | | |
| Pectobacterium | FALSE | 3 | | | | | | | | | 0.333 | 1 | 2 | | |
| Homo | FALSE | 28 | | | | | | | | | 0.286 | 8 | 20 | | |
| Streptococcus | FALSE | 32 | | | | | | | | | 0.281 | 9 | 23 | | |
| Neisseria | FALSE | 28 | | | | | | | | | 0.250 | 7 | 21 | | |
| Cedecea | FALSE | 4 | | | | | | | | | 0.250 | 1 | 3 | | |
| Sodaliphilus | FALSE | 15 | | | | | | | | | 0.200 | 3 | 12 | | |
| Lachnospira | FALSE | 11 | | | | | | | | | 0.182 | 2 | 9 | | |
| Bacillus | FALSE | 12 | | | | | | | | | 0.167 | 2 | 10 | | |
| Duncaniella | FALSE | 6 | | | | | | | | | 0.167 | 1 | 5 | | |
| Quercus | FALSE | 15 | | | | | | | | | 0.133 | 2 | 13 | | |
| Juglans | FALSE | 8 | | | | | | | | | 0.125 | 1 | 7 | | |
| Gossypium | FALSE | 17 | | | | | | | | | 0.059 | 1 | 16 | | |
| Synechococcus | FALSE | 1 | | | | | | | | | 0.000 | 0 | 1 | | |
| Telopea | FALSE | 3 | | | | | | | | | 0.000 | 0 | 3 | | |
| Cobetia | FALSE | 3 | | | | | | | | | 0.000 | 0 | 3 | | |
| Spiroplasma | FALSE | 1 | | | | | | | | | 0.000 | 0 | 1 | | |
| Nicotiana | FALSE | 10 | | | | | | | | | 0.000 | 0 | 10 | | |
| Loigolactobacillus | FALSE | 1 | | | | | | | | | 0.000 | 0 | 1 | | |
| Paenibacillus | FALSE | 9 | | | | | | | | | 0.000 | 0 | 9 | | |
| Purpureocillium | FALSE | 1 | | | | | | | | | 0.000 | 0 | 1 | | |
| Sphingobacterium | FALSE | 6 | | | | | | | | | 0.000 | 0 | 6 | | |
| Besnoitia | FALSE | 1 | | | | | | | | | 0.000 | 0 | 1 | | |
| Cellulophaga | FALSE | 1 | | | | | | | | | 0.000 | 0 | 1 | | |
| Lignipirellula | FALSE | 1 | | | | | | | | | 0.000 | 0 | 1 | | |
| Fusarium | FALSE | 2 | | | | | | | | | 0.000 | 0 | 2 | | |
| Mesomycoplasma | FALSE | 2 | | | | | | | | | 0.000 | 0 | 2 | | |
| Cryphonectria | FALSE | 1 | | | | | | | | | 0.000 | 0 | 1 | | |
| Nocardia | FALSE | 2 | | | | | | | | | 0.000 | 0 | 2 | | |
| Fenollaria | FALSE | 2 | | | | | | | | | 0.000 | 0 | 2 | | |
| Mucilaginibacter | FALSE | 3 | | | | | | | | | 0.000 | 0 | 3 | | |
| Lacticaseibacillus | FALSE | 1 | | | | | | | | | 0.000 | 0 | 1 | | |
| Fluviicola | FALSE | 1 | | | | | | | | | 0.000 | 0 | 1 | | |
| Solidesulfovibrio | FALSE | 1 | | | | | | | | | 0.000 | 0 | 1 | | |
| Alkalitalea | FALSE | 1 | | | | | | | | | 0.000 | 0 | 1 | | |
| Anaerococcus | FALSE | 1 | | | | | | | | | 0.000 | 0 | 1 | | |
| Talaromyces | FALSE | 2 | | | | | | | | | 0.000 | 0 | 2 | | |
| Candida/Metschnikowiaceae | FALSE | 2 | | | | | | | | | 0.000 | 0 | 2 | | |
| Paraprevotella | FALSE | 8 | | | | | | | | | 0.000 | 0 | 8 | | |
| Sesamum | FALSE | 5 | | | | | | | | | 0.000 | 0 | 5 | | |
| Aeromicrobium | FALSE | 1 | | | | | | | | | 0.000 | 0 | 1 | | |
| Muricauda | FALSE | 3 | | | | | | | | | 0.000 | 0 | 3 | | |
| Streptomyces | FALSE | 5 | | | | | | | | | 0.000 | 0 | 5 | | |
| Antarcticibacterium | FALSE | 1 | | | | | | | | | 0.000 | 0 | 1 | | |
| Salegentibacter | FALSE | 1 | | | | | | | | | 0.000 | 0 | 1 | | |
| Ureibacillus | FALSE | 1 | | | | | | | | | 0.000 | 0 | 1 | | |
| Methanocaldococcus | FALSE | 1 | | | | | | | | | 0.000 | 0 | 1 | | |
| Micromonospora | FALSE | 1 | | | | | | | | | 0.000 | 0 | 1 | | |
| Odoribacter | FALSE | 2 | | | | | | | | | 0.000 | 0 | 2 | | |
| Elaeis | FALSE | 15 | | | | | | | | | 0.000 | 0 | 15 | | |
| Prunus | FALSE | 1 | | | | | | | | | 0.000 | 0 | 1 | | |
| Pantoea | FALSE | 1 | | | | | | | | | 0.000 | 0 | 1 | | |
| Prescottella | FALSE | 1 | | | | | | | | | 0.000 | 0 | 1 | | |
| Kayvirus | FALSE | 1 | | | | | | | | | 0.000 | 0 | 1 | | |
| Herbinix | FALSE | 1 | | | | | | | | | 0.000 | 0 | 1 | | |
| Heliorestis | FALSE | 1 | | | | | | | | | 0.000 | 0 | 1 | | |
| Spirosoma | FALSE | 2 | | | | | | | | | 0.000 | 0 | 2 | | |
| Polaribacter | FALSE | 3 | | | | | | | | | 0.000 | 0 | 3 | | |
| Acetivibrio | FALSE | 1 | | | | | | | | | 0.000 | 0 | 1 | | |
| Paeniclostridium | FALSE | 1 | | | | | | | | | 0.000 | 0 | 1 | | |
| Malaciobacter | FALSE | 1 | | | | | | | | | 0.000 | 0 | 1 | | |
| Anaplasma | FALSE | 1 | | | | | | | | | 0.000 | 0 | 1 | | |
| Phoenix | FALSE | 2 | | | | | | | | | 0.000 | 0 | 2 | | |
| Geosporobacter | FALSE | 1 | | | | | | | | | 0.000 | 0 | 1 | | |
| Microdochium | FALSE | 1 | | | | | | | | | 0.000 | 0 | 1 | | |
| Paraburkholderia | FALSE | 1 | | | | | | | | | 0.000 | 0 | 1 | | |
| Corallococcus | FALSE | 1 | | | | | | | | | 0.000 | 0 | 1 | | |
| Pseudogymnoascus | FALSE | 1 | | | | | | | | | 0.000 | 0 | 1 | | |
| Shewanella | FALSE | 2 | | | | | | | | | 0.000 | 0 | 2 | | |
| Buchnera | FALSE | 1 | | | | | | | | | 0.000 | 0 | 1 | | |
| Candidatus Pelagibacter | FALSE | 1 | | | | | | | | | 0.000 | 0 | 1 | | |
| Pseudorhodoplanes | FALSE | 1 | | | | | | | | | 0.000 | 0 | 1 | | |
| Helicobacter | FALSE | 2 | | | | | | | | | 0.000 | 0 | 2 | | |
| Kaistella | FALSE | 1 | | | | | | | | | 0.000 | 0 | 1 | | |
| Listeria | FALSE | 1 | | | | | | | | | 0.000 | 0 | 1 | | |
| Sphingobium | FALSE | 5 | | | | | | | | | 0.000 | 0 | 5 | | |
| Phnomibacter | FALSE | 3 | | | | | | | | | 0.000 | 0 | 3 | | |
| Pediococcus | FALSE | 2 | | | | | | | | | 0.000 | 0 | 2 | | |
| Rothia | FALSE | 3 | | | | | | | | | 0.000 | 0 | 3 | | |
| Triticum | FALSE | 31 | | | | | | | | | 0.000 | 0 | 31 | | |
| Taylorella | FALSE | 1 | | | | | | | | | 0.000 | 0 | 1 | | |
| Psychrobacter | FALSE | 3 | | | | | | | | | 0.000 | 0 | 3 | | |
| Gilliamella | FALSE | 1 | | | | | | | | | 0.000 | 0 | 1 | | |
| Scytonema | FALSE | 2 | | | | | | | | | 0.000 | 0 | 2 | | |
| Nosema | FALSE | 1 | | | | | | | | | 0.000 | 0 | 1 | | |
| Rhizobium | FALSE | 2 | | | | | | | | | 0.000 | 0 | 2 | | |
| Enterocloster | FALSE | 2 | | | | | | | | | 0.000 | 0 | 2 | | |
| Neoantrodia | FALSE | 1 | | | | | | | | | 0.000 | 0 | 1 | | |
| Iamia | FALSE | 1 | | | | | | | | | 0.000 | 0 | 1 | | |
| Eremomyces | FALSE | 1 | | | | | | | | | 0.000 | 0 | 1 | | |
| Acinetobacter | FALSE | 9 | | | | | | | | | 0.000 | 0 | 9 | | |
| Ustilaginoidea | FALSE | 2 | | | | | | | | | 0.000 | 0 | 2 | | |
| Algibacter | FALSE | 1 | | | | | | | | | 0.000 | 0 | 1 | | |
| Rummeliibacillus | FALSE | 1 | | | | | | | | | 0.000 | 0 | 1 | | |
| Ziziphus | FALSE | 5 | | | | | | | | | 0.000 | 0 | 5 | | |
| Punica | FALSE | 1 | | | | | | | | | 0.000 | 0 | 1 | | |
| Lasiodiplodia | FALSE | 1 | | | | | | | | | 0.000 | 0 | 1 | | |
| Metallosphaera | FALSE | 1 | | | | | | | | | 0.000 | 0 | 1 | | |
| Coffea | FALSE | 5 | | | | | | | | | 0.000 | 0 | 5 | | |
| Alternaria | FALSE | 1 | | | | | | | | | 0.000 | 0 | 1 | | |
| Pseudoalteromonas | FALSE | 1 | | | | | | | | | 0.000 | 0 | 1 | | |
| Massilistercora | FALSE | 1 | | | | | | | | | 0.000 | 0 | 1 | | |
| Lactuca | FALSE | 15 | | | | | | | | | 0.000 | 0 | 15 | | |
| Haemophilus | FALSE | 1 | | | | | | | | | 0.000 | 0 | 1 | | |
| Subdoligranulum | FALSE | 1 | | | | | | | | | 0.000 | 0 | 1 | | |
| Ipomoea | FALSE | 4 | | | | | | | | | 0.000 | 0 | 4 | | |
| Corynebacterium | FALSE | 2 | | | | | | | | | 0.000 | 0 | 2 | | |
| Desulfurobacterium | FALSE | 1 | | | | | | | | | 0.000 | 0 | 1 | | |
| Dokdonia | FALSE | 1 | | | | | | | | | 0.000 | 0 | 1 | | |
| Acidihalobacter | FALSE | 1 | | | | | | | | | 0.000 | 0 | 1 | | |
| Hordeum | FALSE | 13 | | | | | | | | | 0.000 | 0 | 13 | | |
| Candidatus Profftia | FALSE | 2 | | | | | | | | | 0.000 | 0 | 2 | | |
| Nostoc | FALSE | 2 | | | | | | | | | 0.000 | 0 | 2 | | |
| Lacinutrix | FALSE | 1 | | | | | | | | | 0.000 | 0 | 1 | | |
| Cajanus | FALSE | 3 | | | | | | | | | 0.000 | 0 | 3 | | |
| Alteromonas | FALSE | 3 | | | | | | | | | 0.000 | 0 | 3 | | |
| Nibribacter | FALSE | 1 | | | | | | | | | 0.000 | 0 | 1 | | |
| Cicer | FALSE | 5 | | | | | | | | | 0.000 | 0 | 5 | | |
| Cupriavidus | FALSE | 1 | | | | | | | | | 0.000 | 0 | 1 | | |
| Rhizophagus | FALSE | 1 | | | | | | | | | 0.000 | 0 | 1 | | |
| Proteiniphilum | FALSE | 1 | | | | | | | | | 0.000 | 0 | 1 | | |
| Exophiala | FALSE | 2 | | | | | | | | | 0.000 | 0 | 2 | | |
| Ananas | FALSE | 8 | | | | | | | | | 0.000 | 0 | 8 | | |
| Tetrapisispora | FALSE | 2 | | | | | | | | | 0.000 | 0 | 2 | | |
| Lolium | FALSE | 7 | | | | | | | | | 0.000 | 0 | 7 | | |
| Flammeovirga | FALSE | 4 | | | | | | | | | 0.000 | 0 | 4 | | |
| Mycobacteroides | FALSE | 1 | | | | | | | | | 0.000 | 0 | 1 | | |
| Gramella | FALSE | 2 | | | | | | | | | 0.000 | 0 | 2 | | |
| Avibacterium | FALSE | 1 | | | | | | | | | 0.000 | 0 | 1 | | |
| Mammaliicoccus | FALSE | 1 | | | | | | | | | 0.000 | 0 | 1 | | |
| Leuconostoc | FALSE | 1 | | | | | | | | | 0.000 | 0 | 1 | | |
| Asparagus | FALSE | 4 | | | | | | | | | 0.000 | 0 | 4 | | |
| Brevibacillus | FALSE | 1 | | | | | | | | | 0.000 | 0 | 1 | | |
| Macadamia | FALSE | 3 | | | | | | | | | 0.000 | 0 | 3 | | |
| Ilyonectria | FALSE | 1 | | | | | | | | | 0.000 | 0 | 1 | | |
| Pseudescherichia | FALSE | 2 | | | | | | | | | 0.000 | 0 | 2 | | |
| Carya | FALSE | 5 | | | | | | | | | 0.000 | 0 | 5 | | |
| Elizabethkingia | FALSE | 1 | | | | | | | | | 0.000 | 0 | 1 | | |
| Methylocystis | FALSE | 2 | | | | | | | | | 0.000 | 0 | 2 | | |
| Yarrowia | FALSE | 1 | | | | | | | | | 0.000 | 0 | 1 | | |
| Phaeoacremonium | FALSE | 1 | | | | | | | | | 0.000 | 0 | 1 | | |
| Puccinia | FALSE | 2 | | | | | | | | | 0.000 | 0 | 2 | | |
| Microcella | FALSE | 1 | | | | | | | | | 0.000 | 0 | 1 | | |
| Kingella | FALSE | 1 | | | | | | | | | 0.000 | 0 | 1 | | |
| Caminibacter | FALSE | 1 | | | | | | | | | 0.000 | 0 | 1 | | |
| Salidesulfovibrio | FALSE | 1 | | | | | | | | | 0.000 | 0 | 1 | | |
| Oscillatoria | FALSE | 2 | | | | | | | | | 0.000 | 0 | 2 | | |
| Caulobacter | FALSE | 4 | | | | | | | | | 0.000 | 0 | 4 | | |
| Pisum | FALSE | 9 | | | | | | | | | 0.000 | 0 | 9 | | |
| Dorea | FALSE | 2 | | | | | | | | | 0.000 | 0 | 2 | | |
| Sulfurovum | FALSE | 1 | | | | | | | | | 0.000 | 0 | 1 | | |
| Brassica | FALSE | 9 | | | | | | | | | 0.000 | 0 | 9 | | |
| Vibrio | FALSE | 7 | | | | | | | | | 0.000 | 0 | 7 | | |
| Mariniblastus | FALSE | 1 | | | | | | | | | 0.000 | 0 | 1 | | |
| Chryseobacterium | FALSE | 11 | | | | | | | | | 0.000 | 0 | 11 | | |
| Dolosigranulum | FALSE | 2 | | | | | | | | | 0.000 | 0 | 2 | | |
| Ascochyta | FALSE | 3 | | | | | | | | | 0.000 | 0 | 3 | | |
| Methanonatronarchaeum | FALSE | 1 | | | | | | | | | 0.000 | 0 | 1 | | |
| Daucus | FALSE | 1 | | | | | | | | | 0.000 | 0 | 1 | | |
| Dysgonomonas | FALSE | 10 | | | | | | | | | 0.000 | 0 | 10 | | |
| Papaver | FALSE | 14 | | | | | | | | | 0.000 | 0 | 14 | | |
| Brachyspira | FALSE | 2 | | | | | | | | | 0.000 | 0 | 2 | | |
| Qiania | FALSE | 1 | | | | | | | | | 0.000 | 0 | 1 | | |
| Desulfuromonas | FALSE | 2 | | | | | | | | | 0.000 | 0 | 2 | | |
| Burkholderia | FALSE | 1 | | | | | | | | | 0.000 | 0 | 1 | | |
| Rhodococcus | FALSE | 1 | | | | | | | | | 0.000 | 0 | 1 | | |
| Phaeobacter | FALSE | 1 | | | | | | | | | 0.000 | 0 | 1 | | |
| Chitinophaga | FALSE | 1 | | | | | | | | | 0.000 | 0 | 1 | | |
| Dioscorea | FALSE | 5 | | | | | | | | | 0.000 | 0 | 5 | | |
| Sorghum | FALSE | 5 | | | | | | | | | 0.000 | 0 | 5 | | |
| Proteus | FALSE | 1 | | | | | | | | | 0.000 | 0 | 1 | | |
| Aquiflexum | FALSE | 1 | | | | | | | | | 0.000 | 0 | 1 | | |
| Marinomonas | FALSE | 1 | | | | | | | | | 0.000 | 0 | 1 | | |
| Zea | FALSE | 4 | | | | | | | | | 0.000 | 0 | 4 | | |
| Fluviispira | FALSE | 1 | | | | | | | | | 0.000 | 0 | 1 | | |
| Pluralibacter | FALSE | 1 | | | | | | | | | 0.000 | 0 | 1 | | |
| Lupinus | FALSE | 7 | | | | | | | | | 0.000 | 0 | 7 | | |
| Oryza | FALSE | 6 | | | | | | | | | 0.000 | 0 | 6 | | |
| Miniphocaeibacter | FALSE | 1 | | | | | | | | | 0.000 | 0 | 1 | | |
| Pseudomonas | FALSE | 10 | | | | | | | | | 0.000 | 0 | 10 | | |
| Tetrahymena | FALSE | 3 | | | | | | | | | 0.000 | 0 | 3 | | |
| Virgibacillus | FALSE | 2 | | | | | | | | | 0.000 | 0 | 2 | | |
| Candidatus Sulfidibacterium | FALSE | 1 | | | | | | | | | 0.000 | 0 | 1 | | |
| Carnobacterium | FALSE | 1 | | | | | | | | | 0.000 | 0 | 1 | | |
| Glycine | FALSE | 6 | | | | | | | | | 0.000 | 0 | 6 | | |
| Eucalyptus | FALSE | 2 | | | | | | | | | 0.000 | 0 | 2 | | |
| Campylobacter | FALSE | 3 | | | | | | | | | 0.000 | 0 | 3 | | |
| Trichoderma | FALSE | 2 | | | | | | | | | 0.000 | 0 | 2 | | |
| Pedobacter | FALSE | 2 | | | | | | | | | 0.000 | 0 | 2 | | |
| Halalkaliarchaeum | FALSE | 1 | | | | | | | | | 0.000 | 0 | 1 | | |
| Brucella | FALSE | 1 | | | | | | | | | 0.000 | 0 | 1 | | |
| Priestia | FALSE | 1 | | | | | | | | | 0.000 | 0 | 1 | | |
| Kosmotoga | FALSE | 1 | | | | | | | | | 0.000 | 0 | 1 | | |
| Camelina | FALSE | 2 | | | | | | | | | 0.000 | 0 | 2 | | |
| Blattabacterium | FALSE | 2 | | | | | | | | | 0.000 | 0 | 2 | | |
| Leptosphaeria | FALSE | 1 | | | | | | | | | 0.000 | 0 | 1 | | |
| Salvia | FALSE | 5 | | | | | | | | | 0.000 | 0 | 5 | | |
| Aeromonas | FALSE | 7 | | | | | | | | | 0.000 | 0 | 7 | | |
| Aggregatibacter | FALSE | 2 | | | | | | | | | 0.000 | 0 | 2 | | |
| Paracoccus | FALSE | 1 | | | | | | | | | 0.000 | 0 | 1 | | |
| Hymenobacter | FALSE | 7 | | | | | | | | | 0.000 | 0 | 7 | | |
| Staphylococcus | FALSE | 6 | | | | | | | | | 0.000 | 0 | 6 | | |
| Venustampulla | FALSE | 1 | | | | | | | | | 0.000 | 0 | 1 | | |
| Jeotgalibaca | FALSE | 1 | | | | | | | | | 0.000 | 0 | 1 | | |
| Thiomicrorhabdus | FALSE | 2 | | | | | | | | | 0.000 | 0 | 2 | | |
| Acaryochloris | FALSE | 2 | | | | | | | | | 0.000 | 0 | 2 | | |
| Myroides | FALSE | 2 | | | | | | | | | 0.000 | 0 | 2 | | |
| Bipolaris | FALSE | 1 | | | | | | | | | 0.000 | 0 | 1 | | |
| Ligilactobacillus | FALSE | 2 | | | | | | | | | 0.000 | 0 | 2 | | |
| Arthrobacter | FALSE | 2 | | | | | | | | | 0.000 | 0 | 2 | | |
| Planococcus | FALSE | 5 | | | | | | | | | 0.000 | 0 | 5 | | |
| Guyanagaster | FALSE | 1 | | | | | | | | | 0.000 | 0 | 1 | | |
| Apibacter | FALSE | 1 | | | | | | | | | 0.000 | 0 | 1 | | |
| Panicum | FALSE | 6 | | | | | | | | | 0.000 | 0 | 6 | | |
| Benincasa | FALSE | 4 | | | | | | | | | 0.000 | 0 | 4 | | |
| Pasteurella | FALSE | 6 | | | | | | | | | 0.000 | 0 | 6 | | |
| Anoxybacillus | FALSE | 1 | | | | | | | | | 0.000 | 0 | 1 | | |
| Hirsutella | FALSE | 21 | | | | | | | | | 0.000 | 0 | 21 | | |
| Olea | FALSE | 6 | | | | | | | | | 0.000 | 0 | 6 | | |
| Manihot | FALSE | 5 | | | | | | | | | 0.000 | 0 | 5 | | |
| Bordetella | FALSE | 1 | | | | | | | | | 0.000 | 0 | 1 | | |
| Mercurialis | FALSE | 1 | | | | | | | | | 0.000 | 0 | 1 | | |
| Brachypodium | FALSE | 2 | | | | | | | | | 0.000 | 0 | 2 | | |
| Methylacidiphilum | FALSE | 1 | | | | | | | | | 0.000 | 0 | 1 | | |
| Legionella | FALSE | 1 | | | | | | | | | 0.000 | 0 | 1 | | |
| Erigeron | FALSE | 3 | | | | | | | | | 0.000 | 0 | 3 | | |
| Aquimarina | FALSE | 2 | | | | | | | | | 0.000 | 0 | 2 | | |
| Eimeria | FALSE | 2 | | | | | | | | | 0.000 | 0 | 2 | | |
| Colwellia | FALSE | 1 | | | | | | | | | 0.000 | 0 | 1 | | |
| Rosa | FALSE | 3 | | | | | | | | | 0.000 | 0 | 3 | | |
| Wickerhamomyces | FALSE | 2 | | | | | | | | | 0.000 | 0 | 2 | | |
| Mergibacter | FALSE | 1 | | | | | | | | | 0.000 | 0 | 1 | | |
| Providencia | FALSE | 2 | | | | | | | | | 0.000 | 0 | 2 | | |
| Citrus | FALSE | 3 | | | | | | | | | 0.000 | 0 | 3 | | |
| Mycena | FALSE | 1 | | | | | | | | | 0.000 | 0 | 1 | | |
| Sphaerotilus | FALSE | 1 | | | | | | | | | 0.000 | 0 | 1 | | |
| Arachis | FALSE | 13 | | | | | | | | | 0.000 | 0 | 13 | | |
| Ktedonosporobacter | FALSE | 1 | | | | | | | | | 0.000 | 0 | 1 | | |
| Halothiobacillus | FALSE | 1 | | | | | | | | | 0.000 | 0 | 1 | | |
| Agrococcus | FALSE | 1 | | | | | | | | | 0.000 | 0 | 1 | | |
| Pseudarthrobacter | FALSE | 1 | | | | | | | | | 0.000 | 0 | 1 | | |
| Chroococcidiopsis | FALSE | 2 | | | | | | | | | 0.000 | 0 | 2 | | |
| Kribbella | FALSE | 1 | | | | | | | | | 0.000 | 0 | 1 | | |
| Potentilla | FALSE | 3 | | | | | | | | | 0.000 | 0 | 3 | | |
| Cucurbita | FALSE | 3 | | | | | | | | | 0.000 | 0 | 3 | | |
| Tatlockia | FALSE | 1 | | | | | | | | | 0.000 | 0 | 1 | | |
| Maricaulis | FALSE | 1 | | | | | | | | | 0.000 | 0 | 1 | | |
| Pseudonocardia | FALSE | 1 | | | | | | | | | 0.000 | 0 | 1 | | |
| Plasmodium | FALSE | 6 | | | | | | | | | 0.000 | 0 | 6 | | |
| Oceanobacillus | FALSE | 1 | | | | | | | | | 0.000 | 0 | 1 | | |
| Brevibacterium | FALSE | 1 | | | | | | | | | 0.000 | 0 | 1 | | |
| Beta | FALSE | 2 | | | | | | | | | 0.000 | 0 | 2 | | |
| Pseudoclavibacter | FALSE | 1 | | | | | | | | | 0.000 | 0 | 1 | | |
| Paraclostridium | FALSE | 5 | | | | | | | | | 0.000 | 0 | 5 | | |
| Mycobacterium | FALSE | 4 | | | | | | | | | 0.000 | 0 | 4 | | |
| Cytobacillus | FALSE | 2 | | | | | | | | | 0.000 | 0 | 2 | | |
| Helianthus | FALSE | 12 | | | | | | | | | 0.000 | 0 | 12 | | |
| Mesorhizobium | FALSE | 1 | | | | | | | | | 0.000 | 0 | 1 | | |
| Algoriphagus | FALSE | 1 | | | | | | | | | 0.000 | 0 | 1 | | |
| Phaseolus | FALSE | 2 | | | | | | | | | 0.000 | 0 | 2 | | |
| Blastomyces | FALSE | 1 | | | | | | | | | 0.000 | 0 | 1 | | |
| Entamoeba | FALSE | 2 | | | | | | | | | 0.000 | 0 | 2 | | |
| Theobroma | FALSE | 1 | | | | | | | | | 0.000 | 0 | 1 | | |
| Scheffersomyces | FALSE | 3 | | | | | | | | | 0.000 | 0 | 3 | | |
| Medicago | FALSE | 8 | | | | | | | | | 0.000 | 0 | 8 | | |
| Viridibacillus | FALSE | 2 | | | | | | | | | 0.000 | 0 | 2 | | |
| Vigna | FALSE | 10 | | | | | | | | | 0.000 | 0 | 10 | | |
| Zymoseptoria | FALSE | 1 | | | | | | | | | 0.000 | 0 | 1 | | |
| Thermosynechococcus | FALSE | 1 | | | | | | | | | 0.000 | 0 | 1 | | |
| Megasphaera | FALSE | 1 | | | | | | | | | 0.000 | 0 | 1 | | |
| Ichthyophthirius | FALSE | 2 | | | | | | | | | 0.000 | 0 | 2 | | |
| Weeksella | FALSE | 1 | | | | | | | | | 0.000 | 0 | 1 | | |
| Flavobacterium | FALSE | 13 | | | | | | | | | 0.000 | 0 | 13 | | |
| Cynara | FALSE | 4 | | | | | | | | | 0.000 | 0 | 4 | | |
| Ricinus | FALSE | 3 | | | | | | | | | 0.000 | 0 | 3 | | |
| Flavisolibacter | FALSE | 3 | | | | | | | | | 0.000 | 0 | 3 | | |
| Sorangium | FALSE | 1 | | | | | | | | | 0.000 | 0 | 1 | | |
| Paramecium | FALSE | 1 | | | | | | | | | 0.000 | 0 | 1 | | |
| Daldinia | FALSE | 1 | | | | | | | | | 0.000 | 0 | 1 | | |
| Paraurantiacibacter | FALSE | 1 | | | | | | | | | 0.000 | 0 | 1 | | |
| Setaria | FALSE | 4 | | | | | | | | | 0.000 | 0 | 4 | | |
| Physcomitrium | FALSE | 3 | | | | | | | | | 0.000 | 0 | 3 | | |
| Coniosporium | FALSE | 1 | | | | | | | | | 0.000 | 0 | 1 | | |
| Borrelia | FALSE | 1 | | | | | | | | | 0.000 | 0 | 1 | | |
| Perkinsus | FALSE | 1 | | | | | | | | | 0.000 | 0 | 1 | | |
| Treponema | FALSE | 1 | | | | | | | | | 0.000 | 0 | 1 | | |
| Anaeromyxobacter | FALSE | 1 | | | | | | | | | 0.000 | 0 | 1 | | |
| Fonsecaea | FALSE | 1 | | | | | | | | | 0.000 | 0 | 1 | | |
| Amycolatopsis | FALSE | 2 | | | | | | | | | 0.000 | 0 | 2 | | |
| Cannabis | FALSE | 3 | | | | | | | | | 0.000 | 0 | 3 | | |
| Megamonas | FALSE | 2 | | | | | | | | | 0.000 | 0 | 2 | | |
| Tardiphaga | FALSE | 1 | | | | | | | | | 0.000 | 0 | 1 | | |
| Pseudoprevotella | FALSE | 4 | | | | | | | | | 0.000 | 0 | 4 | | |
| Aspergillus | FALSE | 6 | | | | | | | | | 0.000 | 0 | 6 | | |
| Magnetospira | FALSE | 1 | | | | | | | | | 0.000 | 0 | 1 | | |
| Stutzerimonas | FALSE | 4 | | | | | | | | | 0.000 | 0 | 4 | | |
| Francisella | FALSE | 2 | | | | | | | | | 0.000 | 0 | 2 | | |
| Salipiger | FALSE | 1 | | | | | | | | | 0.000 | 0 | 1 | | |
| Fragaria | FALSE | 3 | | | | | | | | | 0.000 | 0 | 3 | | |
| Zunongwangia | FALSE | 1 | | | | | | | | | 0.000 | 0 | 1 | | |
| Impatiens | FALSE | 2 | | | | | | | | | 0.000 | 0 | 2 | | |
| Gordonia | FALSE | 1 | | | | | | | | | 0.000 | 0 | 1 | | |
| Candidatus Nitrosomarinus | FALSE | 2 | | | | | | | | | 0.000 | 0 | 2 | | |
| Mus | FALSE | 116 | | | | | | | | | 0.000 | 0 | 116 | | |
| Yersinia | FALSE | 1 | | | | | | | | | 0.000 | 0 | 1 | | |
| Metarhizium | FALSE | 1 | | | | | | | | | 0.000 | 0 | 1 | | |
| Sphingopyxis | FALSE | 1 | | | | | | | | | 0.000 | 0 | 1 | | |
| Fictibacillus | FALSE | 1 | | | | | | | | | 0.000 | 0 | 1 | | |
| Zingiber | FALSE | 15 | | | | | | | | | 0.000 | 0 | 15 | | |
| Solanum | FALSE | 17 | | | | | | | | | 0.000 | 0 | 17 | | |
| Cucumis | FALSE | 5 | | | | | | | | | 0.000 | 0 | 5 | | |
| Leptotrichia | FALSE | 1 | | | | | | | | | 0.000 | 0 | 1 | | |
| Sulfuricella | FALSE | 2 | | | | | | | | | 0.000 | 0 | 2 | | |
| Nymphaea | FALSE | 3 | | | | | | | | | 0.000 | 0 | 3 | | |
| Bathymodiolus thermophilus th | FALSE | 3 | | | | | | | | | 0.000 | 0 | 3 | | |
| Toxoplasma | FALSE | 1 | | | | | | | | | 0.000 | 0 | 1 | | |
| Seonamhaeicola | FALSE | 1 | | | | | | | | | 0.000 | 0 | 1 | | |
| Duganella | FALSE | 1 | | | | | | | | | 0.000 | 0 | 1 | | |
| Mycolicibacterium | FALSE | 2 | | | | | | | | | 0.000 | 0 | 2 | | |
| Musa | FALSE | 4 | | | | | | | | | 0.000 | 0 | 4 | | |
| Capsicum | FALSE | 22 | | | | | | | | | 0.000 | 0 | 22 | | |
| Gluconobacter | FALSE | 2 | | | | | | | | | 0.000 | 0 | 2 | | |
| Actinoalloteichus | FALSE | 1 | | | | | | | | | 0.000 | 0 | 1 | | |
| Mangrovivirga | FALSE | 1 | | | | | | | | | 0.000 | 0 | 1 | | |
| Malus | FALSE | 8 | | | | | | | | | 0.000 | 0 | 8 | | |
| Pontibacter | FALSE | 1 | | | | | | | | | 0.000 | 0 | 1 | | |
| Leptothermofonsia | FALSE | 1 | | | | | | | | | 0.000 | 0 | 1 | | |
| Lacunisphaera | FALSE | 1 | | | | | | | | | 0.000 | 0 | 1 | | |
| Trifolium | FALSE | 3 | | | | | | | | | 0.000 | 0 | 3 | | |
| Arachidicoccus | FALSE | 1 | | | | | | | | | 0.000 | 0 | 1 | | |
| Debaryomyces | FALSE | 2 | | | | | | | | | 0.000 | 0 | 2 | | |
| Sporosarcina | FALSE | 2 | | | | | | | | | 0.000 | 0 | 2 | | |
| Mangifera | FALSE | 3 | | | | | | | | | 0.000 | 0 | 3 | | |
| Vitis | FALSE | 5 | | | | | | | | | 0.000 | 0 | 5 | | |
| Lysinibacillus | FALSE | 1 | | | | | | | | | 0.000 | 0 | 1 | | |
| Tenacibaculum | FALSE | 1 | | | | | | | | | 0.000 | 0 | 1 | | |
| Sum | | 90361 | | | | | | | | | | | | | |
