## Supplementary Table 12 for "MetaGut: Insights into gut microbiomes in stem cell transplantation by comprehensive shotgun long-read sequencing"

| Zymo Genome | Weighted FastANI |
| --- | --- |
| ../../zymo_genomes/Akkermansia_muciniphila.fasta | 99.8943 |
| ../../zymo_genomes/Bacteroides_fragilis.fasta | 99.7164 |
| ../../zymo_genomes/Bifidobacterium_adolescentis.fasta | 99.8568024354723 |
| ../../zymo_genomes/Candida_albican.fasta | 96.1765469526184 |
| ../../zymo_genomes/Clostridioides_difficile.fasta | 99.9472 |
| ../../zymo_genomes/Escherichia_coli_B1109.fasta | 99.7628675875021 |
| ../../zymo_genomes/Escherichia_coli_B3008.fasta | 99.5789948864936 |
| ../../zymo_genomes/Escherichia_coli_B766.fasta | 99.6691412256221 |
| ../../zymo_genomes/Escherichia_coli_JM109.fasta | 99.6824412541286 |
| ../../zymo_genomes/Escherichia_coli_b2207.fasta | 99.8536193911511 |
| ../../zymo_genomes/Faecalibacterium_prausnitzii.fasta | 99.9206853141039 |
| ../../zymo_genomes/Fusobacterium_nucleatum.fasta | 99.9362 |
| ../../zymo_genomes/Lactobacillus_fermentum.fasta | 99.8957 |
| ../../zymo_genomes/Methanobrevibacter_smithii.fasta | 99.7695 |
| ../../zymo_genomes/Prevotella_corporis.fasta | 99.8694310892808 |
| ../../zymo_genomes/Roseburia_hominis.fasta | 99.9604 |
| ../../zymo_genomes/Saccharomyces_cerevisiae.fasta | 99.6749754533036 |
| ../../zymo_genomes/Salmonella_enterica.fasta | 96.5858 |
| ../../zymo_genomes/Veillonella_rogosae.fasta | 99.9602 |
